# An Integrative Single-Cell Epigenomic Atlas Annotates the Regulatory Genome of the Adult Mouse Brain

**DOI:** 10.64898/2026.02.07.704075

**Authors:** Zhaoning Wang, Songpeng Zu, Ethan J. Armand, Timothy H. Loe, Jonathan A. Rink, Wanying Wu, Yang Xie, Lei Chang, Chenxu Zhu, Nicholas D. Johnson, Jasper Lee, Jackson K. Willier, Silvia Cho, Stella Cao, Ariana S. Barcoma, Nora Emerson, Hanqing Liu, Kangli Wang, Bharath Saravanan, Zane A. Gibbs, Xiaomeng Gao, Sunan Xu, David Guo, Zhuowen Tu, Yang E. Li, Joseph R. Ecker, M. Margarita Behrens, Bing Ren

## Abstract

Histone modifications underpin the cell-type-specific gene regulatory programs that drive the remarkable cellular diversity of the mammalian brain. Here, we jointly profiled four histone modifications and the transcriptome in 2.5 million nuclei from the adult mouse brain. By integrating these data with chromatin accessibility, DNA methylation, and three-dimensional genome organization, we constructed a unified epigenomic atlas spanning over 100 brain cell subclasses, and assigned active, primed, and repressive chromatin states across 81% of the genome. Active chromatin states marked by combinatorial histone modifications better nominate candidate enhancers with functional support than chromatin accessibility alone, while H3K27me3- and H3K9me3-marked chromatin states delineate distinct modes of cell-type-specific gene repression. Finally, this multimodal resource enables deep learning models that predict cell-type-specific epigenomic features and gene expression from DNA sequence, providing a comprehensive framework for annotating the brain regulatory genome and interpreting non-coding disease risk variants.

**HIGHLIGHTS:**

- A single-cell epigenome atlas of transcription and four histone modifications across 2.5M mouse brain cells
- Multimodal integration maps chromatin states across ∼81% of the adult mouse brain genome
- Cell-type-resolved chromatin landscapes reveal regulatory programs mediated by enhancers, Polycomb and H3K9me3
- Deep learning models predict cell-type-specific epigenomic features and gene expression from DNA sequence

## INTRODUCTION

The mammalian brain contains an exceptionally diverse array of cell types that differ in developmental origin, molecular identity, anatomical distribution, and function^1,2^. Large-scale single-cell and spatial omics studies, including those from the BRAIN Initiative Cell Census Network (BICCN) and Allen Brain Atlas, have established comprehensive taxonomies of brain cell types and revealed extensive cellular heterogeneity across brain regions and species^3–31^. These studies have provided a foundational framework for understanding brain organization, but transcriptional profiles alone offer limited insight into the regulatory mechanisms that establish and maintain cell-type-specific gene expression programs.

Gene regulation in all mammalian cells is governed by a multilayered epigenome that includes chromatin accessibility, histone modifications, DNA methylation, and three-dimensional (3D) genome organization^32,33^. In the mammalian brain, the extraordinary diversity of cell types, coupled with their precise spatial organization and developmental lineages, greatly amplifies regulatory complexity, necessitating coordinated interrogation of multiple epigenomic layers to resolve cell-type-specific regulatory mechanisms. Recent single-cell epigenomic studies have begun to map chromatin accessibility, DNA methylation and chromatin conformation at cell-type resolution in the mouse and human brain, substantially expanding catalogs of candidate *cis-*regulatory elements (cCREs) and revealing their cell-type-specific deployment^34–44^. However, histone modification landscapes, which represent a central regulatory layer, remain poorly resolved at single-cell resolution across brain cell types. Consequently, it remains unclear how active and repressive chromatin states contribute to the establishment and maintenance of cell-type-specific regulatory programs in the mammalian brain.

Histone modifications play essential roles in gene regulation by modulating the accessibility of regulatory elements to transcription machinery and facilitating communication between distal enhancers and target gene promoters^45–49^. Acetylation and methylation of specific histone residues distinguish active, primed, poised, and repressed regulatory states, and these signatures have been extensively characterized in bulk tissues and cultured cell lines^50–54^. For example, acetylation of lysine 27 on histone H3 (H3K27ac) marks both active promoters and enhancers, trimethylation of lysine 27 on histone H3 (H3K27me3) marks facultative heterochromatin regions, monomethylation of lysine 4 on histone H3 (H3K4me1) is associated with both active and primed enhancers, and trimethylation of lysine 9 on histone H3 (H3K9me3) demarcates constitutive heterochromatin^55–59^. In complex tissues such as the brain, however, bulk profiling obscures cell-type-specific chromatin states, while technical challenges have constrained the scale and scope of single-cell histone modification mapping^60^. As a result, how active and repressive histone programs are deployed across diverse brain cell types, and how they integrate with other epigenomic layers, remains largely unknown.

Recent methodological advances have enabled single-cell profiling of histone modifications, either as individual modalities or jointly with the transcriptome^61–70^. In particular, we previously developed Paired-Tag (parallel analysis of individual cells for RNA expression and DNA from targeted tagmentation by sequencing), a multi-omic strategy that simultaneously profiles nuclear transcriptome and histone modifications in single cells using combinatorial indexing^62^. This approach enables direct linkage of chromatin state and transcriptional output in heterogeneous tissues and is scalable to atlas-level studies. However, a comprehensive application of this strategy to systematically map histone modification landscapes across the adult brain has not yet been achieved.

To define how multiple chromatin regulatory programs cooperate to establish brain cell identity, we generated a large-scale single-cell epigenomic atlas of the adult mouse brain by jointly profiling transcription and histone modifications in over 2.5 million nuclei sampled from multiple brain regions. By integrating these data with existing single-cell maps of chromatin accessibility, DNA methylation, and 3D genome organization^34,35^, and anchoring all modalities to a unified brain cell-type taxonomy, we constructed a unified regulatory framework spanning more than 100 neuronal and non-neuronal subclasses. This integrative atlas defines active, primed, open, and repressive chromatin states across the genome in each cell type, reveals complementary contributions of enhancer activation, H3K27me3-mediated repression, and H3K9me3-associated heterochromatin to cell-type-specific gene regulation, and demonstrates that combinatorial histone modification signatures more accurately identify candidate enhancers with functional support than chromatin accessibility alone. Finally, we show that this single-cell multimodal epigenome atlas enables sequence-based deep learning models to predict cell-type-specific epigenomic features and gene expression directly from DNA sequence, providing a framework for interpreting the regulatory genome of complex mammalian tissues.

## RESULTS

### A unified multimodal epigenomic atlas of the adult mouse brain

To define how multiple chromatin regulatory programs cooperate to establish cell-type-specific gene expression in the mammalian brain, we generated a large-scale multimodal epigenomic atlas by applying Paired-Tag to nuclei isolated from nine anatomically and functionally distinct regions of the adult mouse brain, including amygdala (AMY), caudate putamen (CPU), entorhinal area (ERC), hippocampus-anterior (HCa), hippocampus-posterior (HCp), hypothalamus (HYP), nucleus accumbens (NAC), prefrontal cortex (PFC, including dorsal anterior cingulate cortex, prelimbic and infralimbic regions), and ventral tegmental area-substantia nigra (VTA-SnR)^71^. These regions were selected for their roles in cognition and memory, and their vulnerabilities in neurodegenerative diseases^72,73^ (Figure 1A, Figures S1A-C). To capture complementary chromatin regulatory programs, we profiled four histone modifications (H3K27ac, H3K4me1, H3K27me3, and H3K9me3), representing major active and repressive chromatin states, jointly with RNA at single cell resolution. We implemented an ultra-high-throughput Paired-Tag workflow with expanded combinatorial barcodes, and performed over 100 independent Paired-Tag reactions across biological replicates from both sexes (Figure 1A, Table 1 and Methods). Following stringent quality control (Figure S1D, Figures S2A, B, and Methods), we obtained over 2.5 million paired RNA-histone modification profiles, representing one of the largest single-cell epigenomic datasets generated for a complex mammalian tissue (Figure 1A and Table S2). The resulting dataset exhibited high complexity, excellent reproducibility across biological replicates, and minimal batch effects. Pseudobulk transcriptomic and histone modification profiles were highly correlated between biological replicates, and low-dimensional embeddings demonstrated robust integration across experiments despite the large scale of the dataset (Figure S3). Together, these results establish a high-quality multimodal resource for systematic analysis of chromatin regulation across diverse brain cell types.

**Figure 1.**
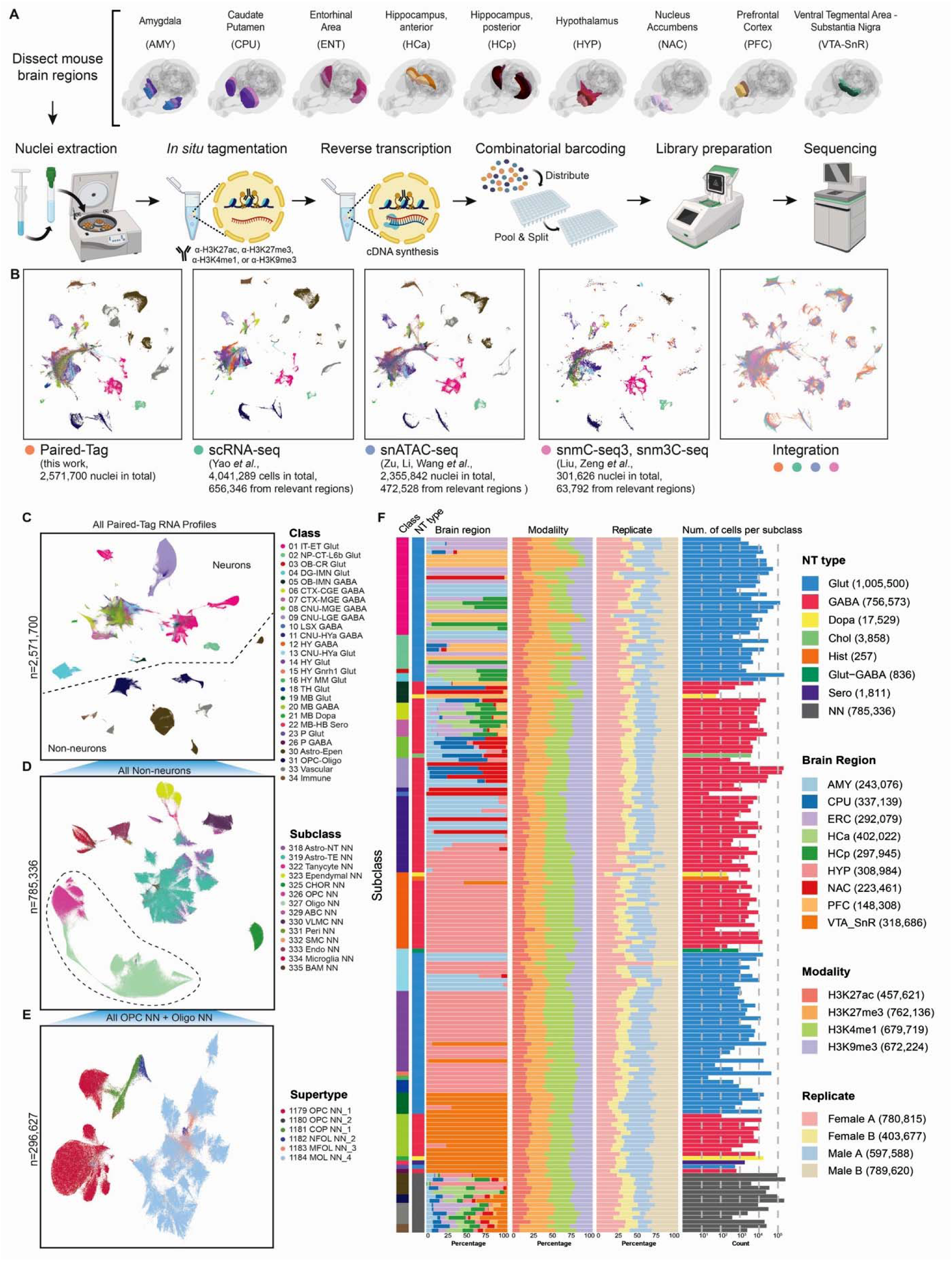
Single-cell co-analysis of gene expression and histone modifications in the adult mouse brain. A. Schematic overview of the Paired-Tag workflow used to jointly profile nuclear transcription and histone modifications in single cells from nine adult mouse brain regions. Brain regions analyzed include amygdala (AMY), caudate putamen (CPU), entorhinal area (ERC), hippocampus-anterior (HCa), hippocampus-posterior (HCp), hypothalamus (HYP), nucleus accumbens (NAC), prefrontal cortex (PFC), and ventral tegmental area-substantia nigra (VTA-SnR). For each region, two biological replicates were generated for both male and female mice by pooling the same region from 2 to 8 animals. Brain region schematics were generated using coordinates from the Allen Mouse Brain Common Coordinate Framework (CCF) v3^71^. B. Integration and joint embedding of Paired-Tag RNA profiles with reference scRNA-seq, snATAC-seq, and snmC-seq3 datasets generated from matched brain regions, enabling unified cell-type annotation. C. Uniform manifold approximation projection (UMAP) embedding of all single-nucleus Paired-Tag RNA profiles analyzed in this study, colored by class labels. D. UMAP visualization of reclustered non-neuronal cells, including the following classes: 30 Astro-Epen, 31 OPC-Oligo, 33 Vascular, and 34 Immune, colored by subclass labels. E. UMAP visualization of single-nucleus Paired-Tag RNA profiles from subclasses 326 OPC NN and 327 Oligo NN, showing progressive differentiation from oligodendrocyte precursor cells (OPCs) to mature oligodendrocytes at supertype resolution. F. Summary bar plots showing, from left to right, class labels, major neurotransmitter (NT) types, major brain region distribution of cells categorized, histone modification modality co-profiled with single-nucleus RNA-seq profiles, biological replicate composition, and the number of nuclei for each subclass present in the dataset.

We next constructed a unified cellular framework by anchoring Paired-Tag transcriptomic profiles to the Allen Brain Cell Atlas taxonomy^6^. First, Paired-Tag snRNA-seq profiles were analyzed by five rounds of iterative clustering using SnapATAC2^74^. This procedure yielded 4,247 L5-level cell clusters, which are independent of the “cluster” annotations of Allen Brain Cell Atlas’ whole mouse brain taxonomy. We then integrated Paired-Tag profiles with the Allen reference, annotated each L5 cluster to supertype resolution, and mapped subclass and class labels using the hierarchical relationships defined in the Allen taxonomy^6^ (Figure 1B). Neuronal and non-neuronal profiles were integrated separately, with neuronal profiles from each brain region aligned to comparable reference regions, resulting in strong correspondence across datasets (Figure S2C). Overall, our dataset provides joint RNA-histone modification profiles for 27 out of 34 major cell classes, 164 out of 338 subclasses, and 519 out of 1201 supertypes represented in the current mouse brain cell taxonomy (Figures 1C-F, Table S2). Cell types absent from the dataset primarily originate from brain regions that were not sampled in this study, such as the thalamus, cerebellum, olfactory bulb, and portions of the midbrain.

Because all Paired-Tag profiles are anchored to a common transcriptomic reference, we next integrated complementary single-cell epigenomic datasets generated by the BICCN consortium from comparable regions, including chromatin accessibility, DNA methylation, and 3D chromatin organization. This integration established a unified multimodal framework spanning transcription, chromatin accessibility, four histone modifications, DNA methylation, and chromatin architecture across more than one hundred neuronal and non-neuronal subclasses^34,35^ (Figure 1B). Representative loci illustrate how this integrated framework captures coordinated regulatory features at canonical marker genes. Cell-type-specific gene expression is consistently accompanied by coordinated chromatin accessibility, active histone marks, depletion of repressive histone marks, reduced DNA methylation, and cell-type-specific chromatin interactions. At loci such as *Satb2* and *Aqp4*, the integrated atlas further identifies distal regulatory elements whose activity is supported simultaneously by active chromatin signatures, three-dimensional chromatin contacts, and coordinated covariance between H3K27ac enrichment and transcription, providing a unified view of gene regulation at single-cell resolution (Figure 2). Although transcriptomic profiles were used to establish the common cellular framework, individual histone modifications independently captured distinct aspects of cellular identity (Figure S2D). The active marks H3K27ac and H3K4me1 robustly resolved neuronal and non-neuronal cell classes, whereas H3K27me3 achieved comparable discrimination despite marking transcriptionally repressed chromatin. In contrast, H3K9me3 primarily distinguished broad neuronal and non-neuronal lineages while unable to resolve finer neuronal classes, which likely reflects both the broad-domain patterns of H3K9me3-mediated constitutive heterochromatin and the greater sparsity of single-cell H3K9me3 profiles in compact heterochromatin regions. Together, these results indicate that individual histone modifications capture cell identity information through distinct regulatory features, with different degrees of resolution under the current profiling depth and analytical framework. Because RNA is co-profiled in Paired-Tag, these results also highlight the value of joint RNA profiling for assigning cell-type-specific epigenomic features in complex tissues.

**Figure 2.**
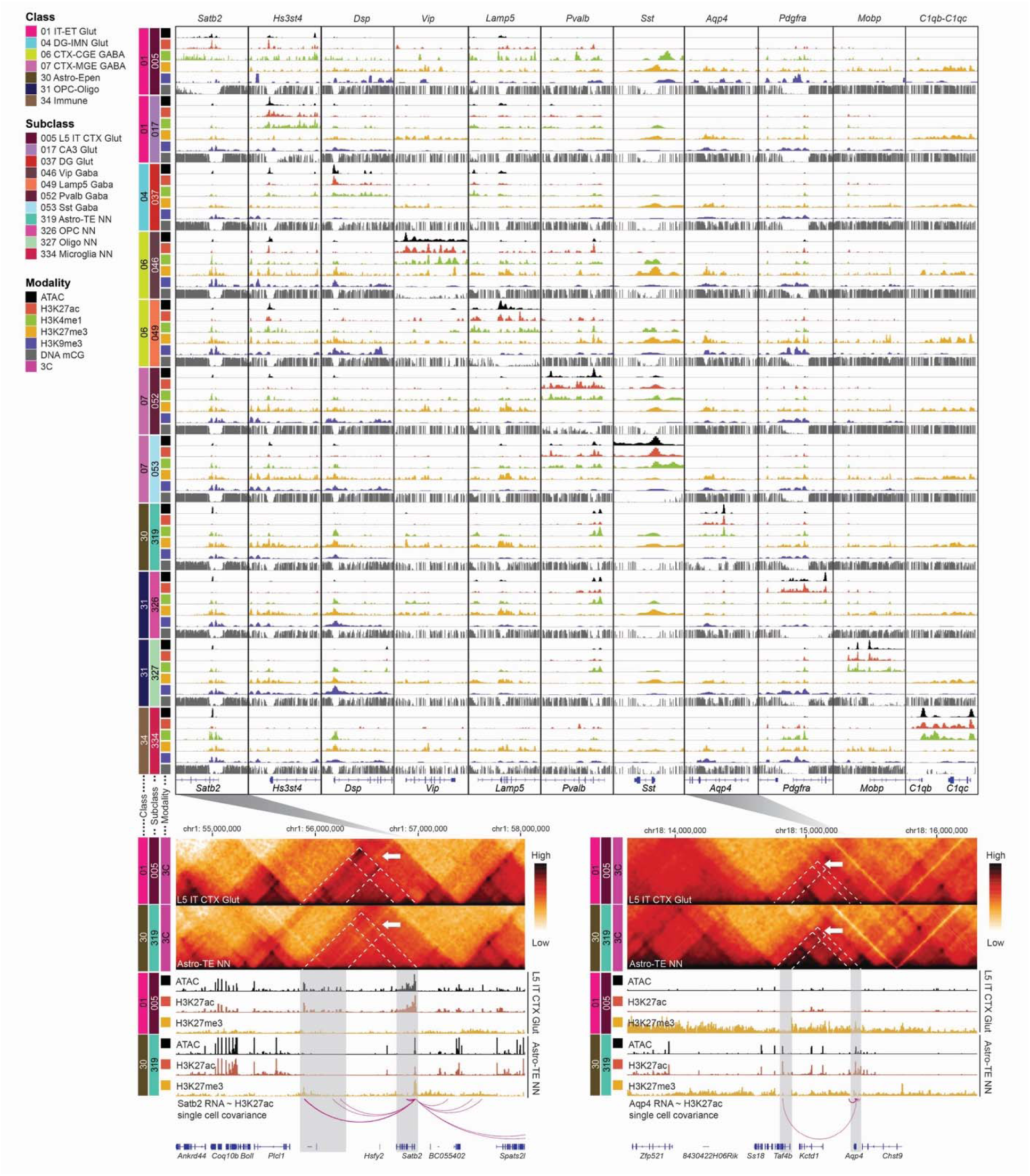
A comprehensive, cell-type-resolved epigenome map of the adult mouse brain. Genome browser views illustrating integrated epigenomic features across representative brain cell subclasses. (Top) Aggregated signals from snATAC-seq, H3K27ac Paired-Tag, H3K27me3 Paired-Tag, H3K4me1 Paired-Tag, H3K9me3 Paired-Tag, and DNA methylation at CG sites (mCG) for selected brain cell subclasses at cell-type-specific marker gene loci. These marker genes include: *Satb2*, *Hs3st4*, *Dsp*, *Vip*, *Lamp5*, *Pvalb*, *Sst*, *Aqp4*, *Pdgfra*, *Mobp*, and *C1qb*. (Bottom) Cell-type-specific chromatin contact maps and Paired-Tag single-cell joint RNA-H3K27ac covariance analysis showing enhancer-promoter interactions at the *Satb2* gene locus in 005 L5 IT CTX Glut, and at the *Aqp4* gene locus in 319 Astro-TE NN, highlighting coordinated activation of distal regulatory elements and target gene expression.

The atlas also captures the remarkable regional specialization of cell types in the adult mouse brain. Consistent with previous transcriptomic studies, most neuronal subclasses exhibited strong anatomical enrichment, while non-neuronal populations were broadly distributed across brain regions^6,35^ (Figure 1F, Figure S4 and Table S2). In total, 117 of 164 subclasses had more than 75% of cells from one dominant region, with the hypothalamus, ventral midbrain, and amygdala contributing the largest numbers of region-specific populations (Figure S4). Furthermore, comparison of anterior and posterior hippocampus revealed spatial enrichment of specific glutamatergic subclasses, including anterior enrichment of 025 CA2-FC-IG Glut and posterior enrichment of 023 SUB-ProS Glut and 033 NP SUB Glut (Figure S4), consistent with the known spatial organization of these hippocampal neuron populations^6,27^.

Collectively, these analyses establish a unified multimodal epigenomic framework that directly links transcription with complementary layers of chromatin regulation across the adult mouse brain. Beyond representing independent epigenomic measurements, the integrated atlas provides a common regulatory framework through which active and repressive chromatin programs can be interpreted together. This framework enables systematic identification of functional chromatin states, regulatory elements, and gene regulatory programs across diverse brain cell types. To facilitate community access, the complete multimodal atlas is available through the CATlas portal (https://catlas.org/catlas/amb-pt), which enables interactive exploration of transcriptomic, epigenomic, and chromatin-state features across all profiled brain cell types.

### Integrated chromatin states reveal the regulatory architecture of the adult mouse brain

A central goal of multimodal epigenomics is to determine how individual regulatory layers collectively specify functional chromatin states. To address this question, we integrated chromatin accessibility with H3K27ac, H3K27me3, H3K4me1, and H3K9me3 profiles using ChromHMM to generate subclass-specific chromatin-state models across the adult mouse brain^75^. Model optimization identified 18 recurrent chromatin signatures that were consolidated into eight biologically interpretable regulatory states: active chromatin (Chr-A), poised chromatin (Chr-Po), primed chromatin (Chr-Pr), open chromatin (Chr-O), repressed chromatin (Chr-R), Polycomb-associated heterochromatin (Hc-P), H3K9me3-associated heterochromatin (Hc-H), and regions lacking sufficient information for confident annotation (Not Determined, ND) (Figures S5A-F; see Methods). Saturation analysis showed that ND coverage decreases with cell sampling depth, indicating that chromatin-state annotation improves with greater per-subclass coverage (Figure S5G). State annotations were consistent between biological replicates (Figures S5D, E), and generally concordant with published chromatin-state models (Figure S6A)^50,54,62,76^.

These integrated chromatin states capture distinct combinations of active and repressive epigenomic features and provide a unified functional annotation of the mouse brain regulatory genome (Figures 3A, B). ND regions occupied the largest fraction of the genome, whereas Chr-Pr, Hc-P, Chr-A, and Hc-H covered substantially more genomic sequence (>10 Mb) than Chr-Po and Chr-R (Figure 3C). Across brain cell subclasses, we found that Chr-Po and Chr-R states are the most variable (Figure S6B). Integration with DNA methylation further revealed that each chromatin state possesses a characteristic epigenetic signature. Active and poised chromatin (Chr-A and Chr-Po) exhibited the lowest levels of both CpG (mCG) and non-CpG methylation (mCH, where H denotes A, C, or T) sites, whereas unannotated regions (ND) and Polycomb-associated (Hc-P) showed the highest mCG and mCH fractions (Figures 3D, E). On CpG islands, we found an inverse correlation of DNA methylation with H3K27me3 signal, consistent with previous reports that DNA methylation counteracts the H3K27me3 writer PRC2 at large CpG islands^77–80^ (Figures S6C). Hc-H also showed a considerable fraction of mCG, and the actual mCG fraction on *bona fide* Hc-H could be underestimated due to the open chromatin bias of Tn5 tagmentation that the DNA modality analysis of Paired-Tag (i.e., CUT&Tag) is based on^81^ (Figure 3D). Previous studies have shown that mCH preferentially accumulates in neurons compared with glia^82,83^. In line with this pattern, we found overall higher mCH fraction in all chromatin states in neuronal subclasses compared with non-neuronal subclasses, especially in Hc-P, suggesting differential involvement of mCH in neuronal gene regulation^34^ (Figures S6D, E). These observations demonstrate that DNA methylation and histone modifications cooperate to define regulatory chromatin states across brain cell types.

**Figure 3.**
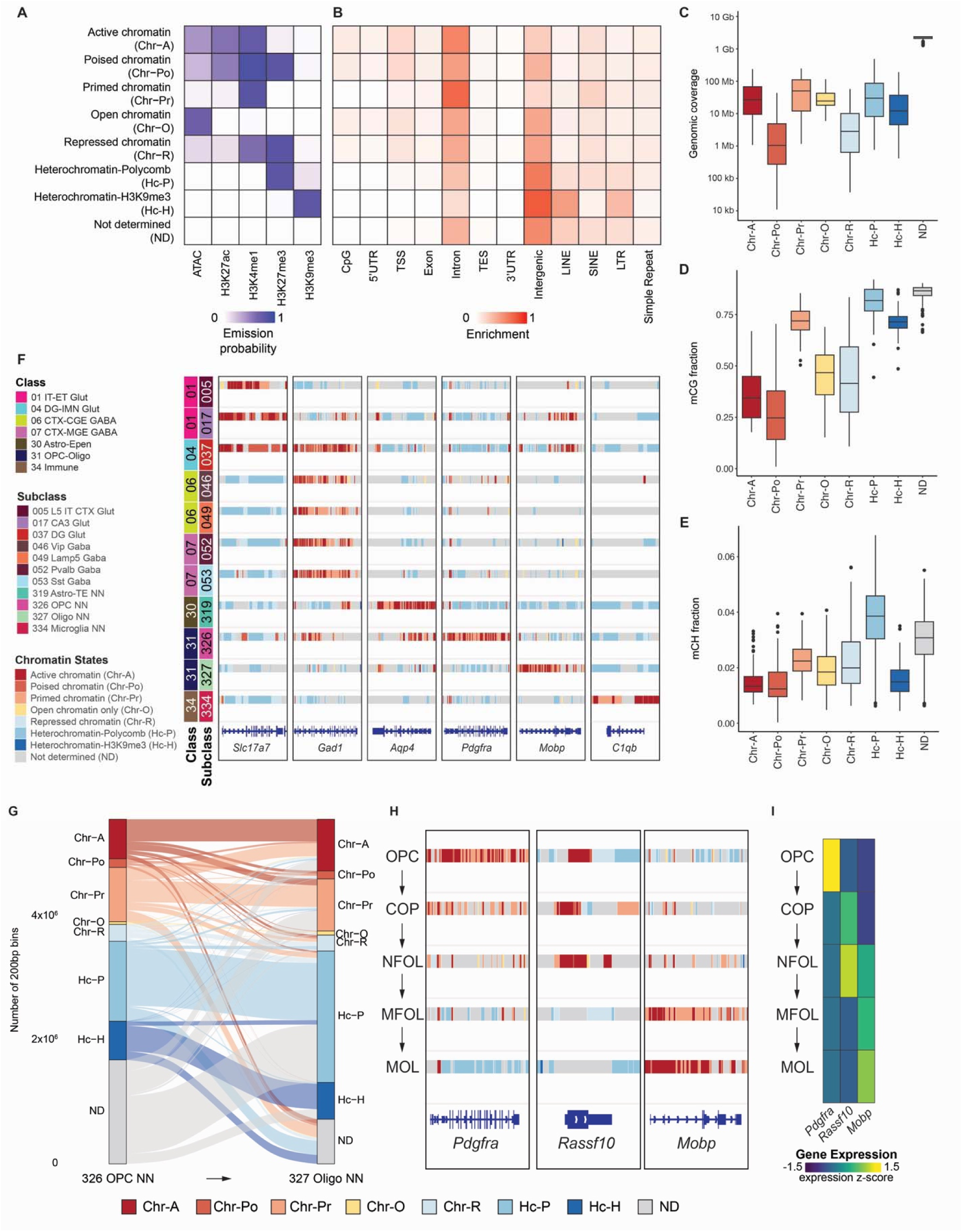
Chromatin state modeling functionally annotates the brain regulatory genome. A. Heatmap showing the emission probability of each epigenomic mark across the 8 chromatin states. B. Heatmap showing the relative enrichment of each chromatin state across different genomic annotations. C. Box plot showing the distribution of genomic coverage of each chromatin state across brain cell subclasses. D. Box plot showing fraction of CG methylation (mCG) associated with each chromatin state across brain cell subclasses. E. Box plot showing fraction of non-CG methylation (mCH) associated with each chromatin state across brain cell subclasses. F. Chromatin state landscapes at the *Slc17a7*, *Gad1*, *Aqp4*, *Pdgfra*, *Mobp*, and *C1qb* loci across selected brain cell subclasses, including 005 L5 IT CTX Glut, 017 CA3 Glut, 037 DG Glut, 046 Vip Gaba, 049 Lamp5 Gaba, 052 Pvalb Gaba, 053 Sst Gaba, 319 Astro-TE NN, 326 OPC NN, 327 Oligo NN, and 334 Microglia NN. G. Alluvial plot illustrating chromatin state transitions between 326 OPC NN and 327 Oligo NN. Chromatin regions that are annotated as ND for both 326 OPC NN and 327 Oligo NN subclasses are not shown. H. Chromatin state landscapes at the *Pdgfra*, *Rassf10*, and *Mobp* loci across selected cell supertypes reflecting cellular state transitions during the oligodendrocyte precursor cell (OPC) to oligodendrocyte differentiation process, including OPC, committed oligodendrocyte precursors (COP), newly formed oligodendrocytes (NFOL), myelin-forming oligodendrocytes (MFOL), and mature oligodendrocytes (MOL). I. Heatmap showing the relative expression of *Pdgfra*, *Rassf10*, and *Mobp* in OPC, COP, NFOL, MFOL and MOLs.

Representative genomic loci illustrate how integrated chromatin states provide a more comprehensive view of gene regulation than any individual modality alone. Cell-type-specific marker genes were consistently associated with active or primed chromatin states in expressing subclasses and with Hc-P or Hc-H states in non-expressing populations (Figure 3F). Rather than simply cataloging individual histone modifications, the integrated chromatin-state framework directly links combinations of epigenomic features to regulatory activity, thereby providing a unified interpretation of chromatin state across diverse cellular contexts.

Across all profiled brain cell subclasses, chromatin-state modeling substantially expanded functional annotation of the mouse genome. Approximately 29% of the genome was assigned Chr-A in at least one subclass, while 81% was covered by at least one non-ND chromatin state assignment across all profiled subclasses under the current sampling depth (Figure S6F). This represents a more than fourfold expansion beyond previous annotations based solely on chromatin accessibility^35^, and demonstrates that integration of complementary epigenomic modalities greatly increases the fraction of the genome that can be functionally annotated.

Together, these analyses establish integrated chromatin states as a common regulatory framework that unifies multiple epigenomic modalities. Instead of treating chromatin accessibility and individual histone modifications as independent measurements, this framework captures their coordinated deployment to define active, primed, and repressive regulatory programs across the adult mouse brain, providing the foundation for systematic investigation of chromatin dynamics, enhancer function, and gene regulatory mechanisms.

### Chromatin-state remodeling accompanies lineage specification and regional specialization

A key advantage of an integrated chromatin-state framework is that it enables regulatory programs to be followed across related cellular states. To illustrate this capability, we examined chromatin-state remodeling during oligodendrocyte differentiation, a well-characterized process accompanied by extensive transcriptional and epigenomic reprogramming. Comparison of oligodendrocyte precursor cells (OPCs, 326 OPC NN) and mature oligodendrocytes (327 Oligo NN) revealed widespread chromatin-state transitions accompanying lineage progression (Table S3). Nearly half of the Chr-A regions identified in OPCs adopted alternative chromatin states following differentiation, while more than 50% of Chr-A regions in mature oligodendrocytes were newly established during lineage maturation (Figure 3G). Most newly acquired Chr-A regions originated from previously primed or unannotated chromatin, indicating progressive activation of lineage-specific regulatory elements rather than global resetting of the regulatory landscape.

Extending this analysis across successive stages of oligodendrocyte maturation, from OPCs to committed oligodendrocyte precursors (COPs), newly formed oligodendrocytes (NFOLs), myelin-forming oligodendrocytes (MFOLs), and mature oligodendrocytes (MOLs)^6^, revealed coordinated chromatin-state transitions that closely paralleled transcriptional progression (Figures 3H,I). Genes associated with progenitor identity, such as *Pdgfra,* progressively lost active and primed chromatin states and acquired Polycomb-associated repression during differentiation^84,85^ (Figure 3H, left), whereas genes required for myelination, including *Mobp*^86^, underwent the reciprocal transition from repressive or primed chromatin to active regulatory states (Figure 3H, right). This analysis also highlighted genes not previously characterized during oligodendrocyte differentiation, such as *Rassf10*, which showed transient chromatin activation during intermediate developmental stages before returning to an inactive chromatin state in mature oligodendrocytes, consistent with its known function in regulating cell proliferation^87,88^ (Figure 3H, middle). These chromatin transitions paralleled the gene expression dynamics of these key lineage genes (Figure 3I), indicating that coordinated remodeling of active and repressive states accompanies OPC-to-oligodendrocyte differentiation. Together, these observations demonstrate that integrated chromatin-state maps can resolve sequential regulatory transitions that accompany lineage commitment and cellular maturation.

The chromatin-state framework also faithfully recapitulated regulatory principles established by previous oligodendrocyte studies. Immune-related loci, including *Tnfrsf1a*, *Psmb9-Tap2*, and *H2-Ab1/H2-Aa*, exhibited preferential Chr-A or Chr-Pr in OPCs consistent with reported primed accessibility at selected immune-related genes^89^ (Figure S7A), whereas canonical OPC regulator genes, including *Sox5*, *Sox6* and *Ptprz1*, progressively accumulated H3K27me3 during oligodendrocyte maturation, consistent with Polycomb-mediated silencing of progenitor programs^64^ (Figure S7B). Together, these analyses demonstrate that chromatin-state transitions provide a mechanistic framework linking dynamic epigenomic remodeling to key lineage-specific transcriptional patterns.

We next asked whether the same framework could resolve regulatory differences between closely related mature cell populations. Comparison of telencephalic astrocytes (Astro-TE, subclass 319) and non-telencephalic astrocytes (Astro-NT, subclass 318) revealed extensive chromatin-state divergence despite their shared cellular identity, with more than one-fifth of the genome assigned to different chromatin states between the two subclasses (Figure S8A). These differences were apparent at subclass-enriched gene loci (Figure S8B, C), and coincided with regional transcriptional specialization and were concentrated at genes that define telencephalic or non-telencephalic astrocyte identity^6,90–92^ (Figure S8C). These results indicate that chromatin-state remodeling continues beyond developmental lineage specification to support regional specialization within mature glial populations. One striking example was provided by the *HoxA* locus, which was shown to retain region-specific open or active chromatin signatures across astrocyte populations despite limited expression of canonical Hox transcription factor genes^64,93^ (Figure S8D). Although the locus remained broadly repressed by H3K27me3 and H3K9me3 in both astrocyte populations, non-telencephalic astrocytes (Astro-NT) retained a focal domain with increased accessibility, H3K27ac, and H3K4me1 that coincided with selective expression of the long non-coding RNA *Hotairm1* (Figure S8D). Single-cell covariance analyses further demonstrated coordinated activation of this regulatory region specifically within Hotairm1-expressing astrocytes (Figure S8E-H), suggesting that localized remodeling of active chromatin can occur within otherwise repressed chromosomal domains. These observations are consistent with the idea that chromatin-state remodeling preserves regional developmental memory while maintaining mature astrocyte identity.

Collectively, these analyses demonstrate that integrative mapping of chromatin states provides a cell-type-resolved view of regulatory genome activity. Beyond annotating regulatory elements, the framework captures progressive activation and repression of lineage-specific regulatory programs during differentiation, and reveals how closely related mature cell populations deploy distinct chromatin states to maintain specialized biological functions.

### Active chromatin states better nominate functional enhancers than open chromatin

Chromatin accessibility is widely used to nominate candidate cis regulatory elements, yet chromatin accessibility alone does not distinguish functional enhancers from other open genomic regions with structural or regulatory roles. We therefore asked whether integrating histone modifications with chromatin accessibility could better nominate functional enhancers across the adult mouse brain.

Using the chromatin-state framework established above, we classified previously identified brain cCREs based on their chromatin states^35^. Across brain cell subclasses, approximately 42% of cCREs were assigned to Chr-A, whereas 54% were Chr-O regions with chromatin accessibility but lacking active histone marks (Figure 4A). Although both classes are accessible, Chr-A regions exhibited substantially stronger signatures of functional regulatory activity. Several independent analyses consistently supported this conclusion. Chr-A regions showed significantly greater evolutionary conservation, measured using the PhastCons conservation scores^94^, than Chr-O regions (Figure 4B), were preferentially enriched among human-conserved candidate regulatory elements (cCREs) ^35,40^ (Figures S9A, B), and were more frequently identified as subclass-specific regulatory elements across the mouse brain (Figure S9C). Chr-A regions also showed shorter pairwise genomic distances than Chr-O regions (Figure 4C), consistent with local clustering of active regulatory elements. Notably, the Chr-A distribution displays a pronounced density at very short distances, which likely reflects groups of densely packed enhancer elements commonly known as “super-enhancers”^95^ (Figure 4C), which we examine later in this section (Figure S10).

**Figure 4.**
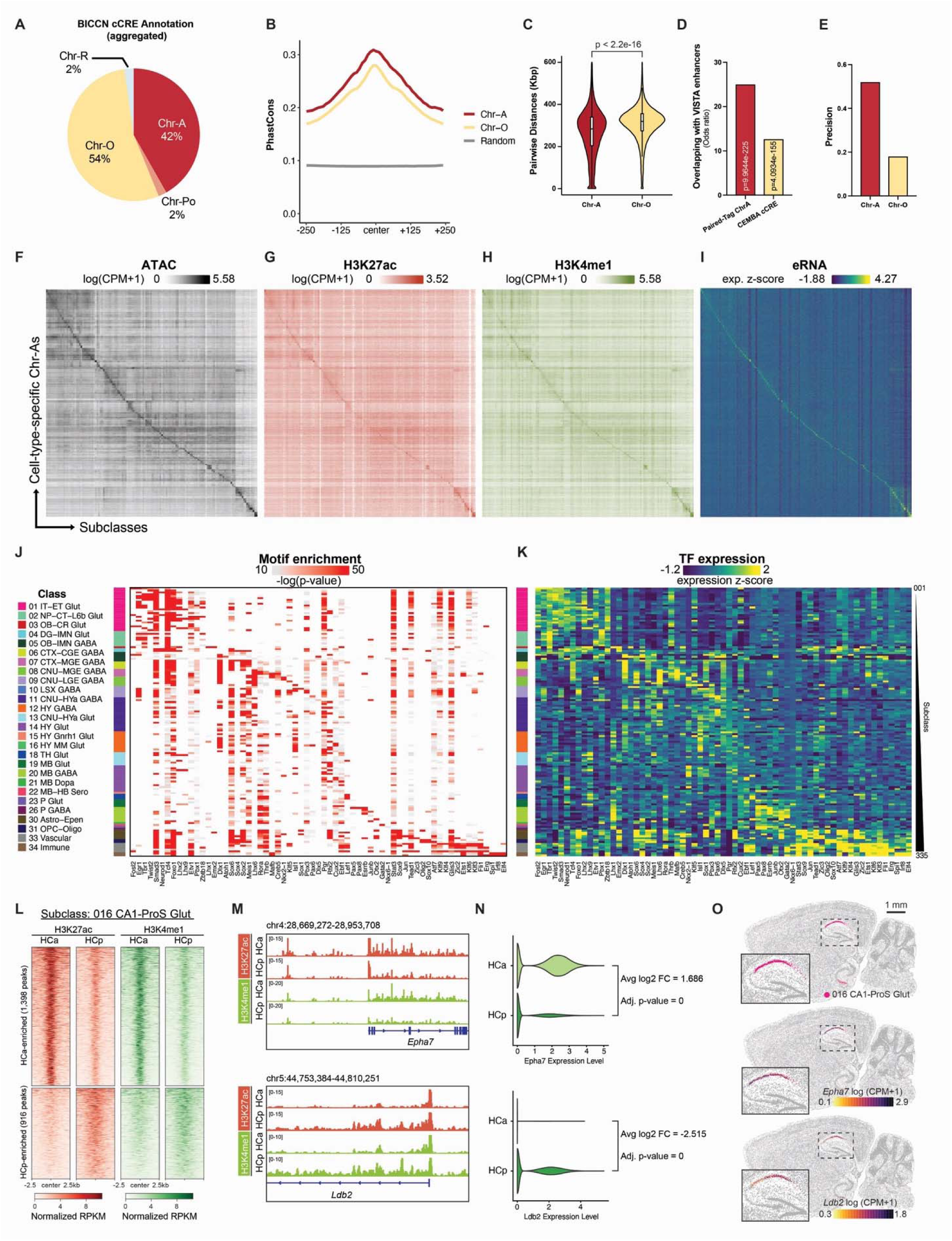
Functional annotation and epigenomic dissection of brain cCREs. A. Pie chart summarizing the overall chromatin state composition of cCREs aggregated across all brain cell subclasses. Brain cCREs were annotated by chromatin state separately within each subclass, and state assignments were then pooled across all subclasses to quantify the global distribution of chromatin states. Brain cCREs were previously defined using snATAC-seq^35^. B. Average PhastCons conservation scores of cCREs annotated as Chr-A (red) or Chr-O (yellow), compared with random genomic background (control) regions (grey). C. Violin plot showing the distribution of pairwise genomic distances between Chr-As or between Chr-Os within 1 Mb genomic bins across brain cell subclasses. Statistical significance was assessed using the Wilcoxon rank-sum test. D. Bar plot showing the odds ratios for overlap between Chr-As identified by Paired-Tag (red) or cCREs identified using snATAC-seq (yellow) and validated enhancers with brain activities from the VISTA enhancer browser database. Statistical significance of overlap was evaluated using Fisher’s exact test. E. Bar plot showing the precision of Chr-A or Chr-O annotation in predicting validated cell-type-specific enhancers from the Allen Institute Enhancer (AiE) collection. Precision is defined as the fraction of predicted enhancer cCREs that were experimentally validated using Allen Institute AAV reporter assays. F. Heatmap showing chromatin accessibility (ATAC-seq signal) at cell-type-specific Chr-As across brain cell subclasses. G. Heatmap showing H3K27ac signal at cell-type-specific Chr-As across brain cell subclasses. H. Heatmap showing H3K4me1 signal at cell-type-specific Chr-As across brain cell subclasses. I. Heatmap showing enhancer RNA (eRNA) signal at cell-type-specific Chr-As across brain cell subclasses. J. Heatmap showing enrichment of cell-type-specific transcription factor binding motifs within Chr-As across brain cell subclasses. K. Heatmap showing the relative expression levels of corresponding cell-type-specific transcription factors across brain cell subclasses. L. Coverage heatmaps of H3K27ac and H3K4me1 signals at differentially enriched H3K27ac peaks between anterior (HCa) and posterior (HCp) hippocampus in subclass 016 CA1-ProS Glut neurons. M. Genome browser tracks showing differential H3K27ac and H3K4me1 signals in 016 CA1-ProS Glut neurons from HCa and HCp at *Epha7* (top) and *Ldb2* (bottom) genomic loci. N. Violin plots showing differential expression of *Epha7* (top) and *Ldb2* (bottom) in 016 CA1-ProS Glut neurons between HCa and HCp. Statistical significance was assessed using the Wilcoxon rank-sum test followed by Bonferroni correction. O. MERFISH analysis of the adult mouse brain showing (top) the spatial distribution of subclass 016 CA1-ProS Glut neurons in a sagittal brain section; (middle) anterior-enriched expression of *Epha7* in these neurons; and (bottom) posterior-enriched expression of *Ldb2*. MERFISH data were obtained from the Allen Brain Cell (ABC) Atlas web portal (https://brain-map.org/bkp/explore/abc-atlas). Scale bar, 1 mm.

We next asked whether Chr-A regions better enrich experimentally validated enhancers. In the VISTA Enhancer Browser^96^, Chr-A regions showed substantially stronger enrichment for validated brain enhancers than accessible chromatin alone (Figure 4D). Because most VISTA enhancers were validated in embryonic mice, this comparison is conservative and may miss adult-specific enhancers. Similar results were obtained using the Allen Institute Enhancer (AiE) collection, in which Chr-A annotations consistently achieved higher precision, recall and specificity for predicting cell-type-specific enhancer activity than Chr-O regions^97^ (Figure 4E, Figures S9D-F). These complementary validation datasets demonstrate that active chromatin states provide a substantially more accurate representation of functional enhancer activity than chromatin accessibility alone.

To infer potential regulators of different chromatin states, we performed motif analysis on Chr-A, Chr-Pr, and Chr-O regions of each brain cell subclass. We detected the largest number of transcription factor (TF) binding motifs enriched in Chr-A regions. In contrast, fewer motifs were enriched in Chr-Pr and Chr-O, especially in non-neuronal Chr-O regions (Figure S9G). In addition, we found lineage-specific TF binding motifs to be most significantly enriched in Chr-A regions, including various NeuroD and Neurogenin subfamily members of basic helix-loop-helix (bHLH) TFs in dentate gyrus glutamatergic neurons (subclass 037 DG Glut) and ETS family member TFs in microglia (subclass 334 Microglia NN) (Figure S9H). We also identified certain TF motifs enriched in Chr-Pr with lower significance, possibly regulating the primed but not yet activated gene programs in response to stress or cellular signaling (Figure S9H). Interestingly, we found that CTCF-related motifs were repeatedly discovered in Chr-O regions of almost every cell subclass we examined (Figures S9H, I). In contrast, motifs from the MEF2 transcription factor family, such as MEF2A, MEF2C, and MEF2D, were predominantly enriched in Chr-A regions in most subclasses (Figure S9J). These findings are consistent with the established role of MEF2 family members in regulating gene regulatory networks across diverse lineages, including both neuronal and non-neuronal cells^98,99^, and further support the notion that Chr-O regions are enriched for structural elements involved in chromatin looping and genome topology. Together, these data highlight differences in sequence conservation, functional outcomes, and motif enrichment between Chr-A and Chr-O, supporting the interpretation that Paired-Tag-defined active chromatin state enriches enhancer elements.

The integrated chromatin-state framework further enabled systematic identification of approximately 500,000 cell-type-specific Chr-A elements across more than one hundred neuronal and non-neuronal subclasses (Figure 4F, Table S4). These regions displayed coordinated chromatin accessibility, H3K27ac enrichment, H3K4me1 deposition, and enhancer RNA transcription (Figures 4G-I), consistent with active enhancer function. Motif enrichment analysis identified 839 significantly enriched TF binding motifs (Figure S9K, Table S5), while integration with subclass-specific TF expression nominated 65 high-confidence lineage regulators whose expression coincided with enrichment of their cognate motifs (Figures 4J, K). Many correspond to well-established regulators of neuronal and glial identity, including TBR1 and NeuroD1 in multiple glutamatergic neuronal subclasses^100–102^, Lhx6 and Dlx1 in GABAergic interneurons^103–106^, Olig2 and Sox10 in oligodendrocyte^107–111^, and Fli1 and Irf8 in vascular cells and immune cell populations^112–117^, supporting the biological coherence of the inferred regulatory networks.

Because active enhancers frequently operate in higher-order regulatory clusters, we next examined whether a subset of Chr-A regions can be organized into super-enhancer domains, which have been strongly implicated in the regulation of key cell identity genes^118–122^. Rank Ordering of Super-Enhancers (ROSE) analysis identified more than 100,000 putative super-enhancers distributed across brain cell subclasses (Figure S10A). Compared with other active regions, super-enhancer-associated Chr-A regions exhibited significantly shorter pairwise genomic distances than other Chr-A regions, strong subclass-specific H3K27ac enrichment, increased enhancer RNA production, and enhanced coupling between chromatin activity and gene expression (Figures S10B-E). Together, these analyses show that the integrative chromatin-state framework captures both individual active regulatory elements and their higher-order organization into super-enhancer domains.

Finally, we asked whether the integrative chromatin-state framework also captures regulatory specialization within shared neuronal subclasses across different brain regions. Comparison of anterior and posterior hippocampal glutamatergic neurons, including 016 CA1-ProS Glut, 017 CA3 Glut, and 037 DG Glut, identified numerous region-specific Chr-A elements where their H3K27ac variations correlated with regional differences in gene expression (Figure 4L, Figures S10F, G). For example, *Epha7* and *Ldb2* displayed reciprocal region-specific active chromatin states within CA1-ProS glutamatergic neurons from HCa and HCp (Figure 4M, N). Consistent with these findings, these patterns were supported by an independent MERFISH analysis (Figure 4O) ^27^. These results suggest that active chromatin landscapes encode regional regulatory specialization even within shared neuronal subclasses.

Taken together, these analyses demonstrate that functional enhancers are defined not simply by chromatin accessibility alone, but by integrative chromatin states that combine accessibility with active histone modifications. By distinguishing active regulatory elements from structurally accessible chromatin, this framework substantially improves functional annotation of the brain regulatory genome and provides a foundation for identifying lineage-specific enhancers, reconstructing transcriptional regulatory networks, and developing cell-type-specific genetic tools.

### H3K27me3 and H3K9me3 define complementary modes of gene repression

Cell identity is determined not only by activation of lineage-appropriate regulatory programs but also by stable repression of genes that specify alternative cellular identities. We therefore used the integrated chromatin-state atlas to investigate how distinct repressive chromatin mechanisms contribute to gene regulation across the adult mouse brain.

Comparison of H3K27me3 and H3K9me3 landscapes revealed fundamentally different patterns of chromatin repression. H3K27me3, deposited by the Polycomb Repressive Complex 2 (PRC2), is a hallmark of facultative heterochromatin^123–125^. By contrast, H3K9me3 marks constitutive heterochromatin and is catalyzed by histone methyltransferases such as SUV39H1 (KMT1A), SUV39H2 (KMT1B), and SETDB1 (KMT1E), exhibiting genomic deposition patterns that are distinct from those of H3K27me3^126,127^. In our integrative single-cell epigenomic atlas, H3K27me3 profiles exhibited substantially greater variation among closely related brain cell subclasses than H3K9me3, consistent with their stronger ability to resolve fine cellular identities under the current profiling depth and feature-calling framework (Figures 5A, B, Figure S2D). Differential analysis likewise identified more than three times as many subclass-specific H3K27me3 features (289,341) as H3K9me3 features (89,025) (Table S4), indicating that Polycomb-mediated repression contributes prominently to regulatory diversification among closely related cell populations.

**Figure 5.**
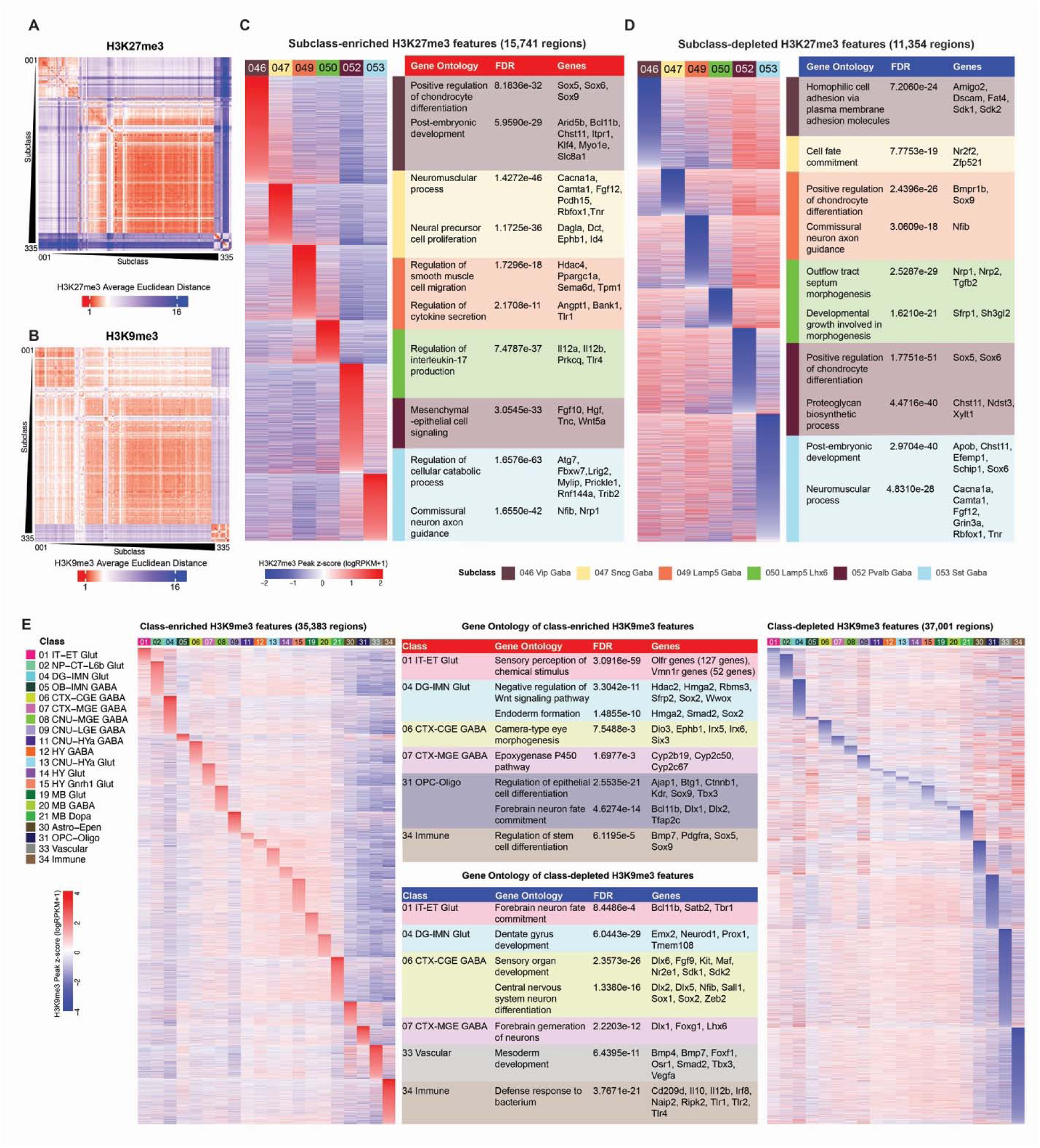
Cell-type-specific repressive histone modification features of the mouse brain. A. Heatmap showing the pairwise Euclidean distance between brain cell subclasses computed from low-dimensional embeddings of single-cell H3K27me3 Paired-Tag profiles. B. Heatmap showing the pairwise Euclidean distance between brain cell subclasses computed from low-dimensional embeddings of single-cell H3K9me3 Paired-Tag profiles. C. Left, heatmap showing normalized H3K27me3 signal over subclass-enriched H3K27me3 features in selected CGE- and MGE-derived GABAergic neuronal subclasses. Right, table showing Gene Ontology terms enriched among these features and their nearest gene targets. D. Left, heatmap showing normalized H3K27me3 signal over subclass-depleted H3K27me3 features in selected CGE- and MGE-derived GABAergic neuronal subclasses. Right, table showing Gene Ontology terms enriched among these features and their nearest gene targets. E. Left, heatmap showing normalized H3K9me3 signal over class-enriched H3K9me3 features across major brain cell classes. Right, heatmap showing normalized H3K9me3 signal over class-depleted H3K9me3 features across major brain cell classes. Middle, table showing Gene Ontology terms enriched among these features and their nearest gene targets in selected cell classes.

To understand how H3K27me3-mediated repression shapes neuronal identity, we examined cortical GABAergic interneuron subclasses representing distinct developmental lineages. H3K27me3 landscapes selectively marked developmental regulators, signaling molecules, and transcription factors that distinguish closely related interneuron populations (Figures 5C,D). For example, the medial ganglionic eminence determinant Lhx6 remained associated with active chromatin only in MGE-derived interneurons, whereas CGE-derived interneurons acquired H3K27me3-associated repression at the same locus (Figure S11C, left). Within the MGE lineage, Sox6, encoding a key transcription factor in regulating the development, migration, and function of Pvalb and Sst interneurons^128,129^, exhibited selective depletion of H3K27me3 specifically in Pvalb and Sst interneurons, illustrating how H3K27me3-mediated repression further diversifies neuronal identity after lineage commitment (Figure S11C, middle). Likewise, differential H3K27me3 occupancy across the *Fgf12* locus, which encodes a key regulator of voltage-gated sodium channels involved in neuronal action potential initiation and propagation^130–132^, corresponded to subclass-specific expression of alternative transcript isoforms (Figure S11C, right), a regulatory pattern that was independently supported by comparative single-cell analyses across mouse, marmoset, macaque, and human cortex^37^ (Figure S11D). Together, these observations demonstrate that H3K27me3-mediated repression refines transcriptional programs among closely related neuronal subtypes.

Analysis of the same cortical GABAergic interneuron subclasses revealed far fewer H3K9me3 features at the same resolution (Figures S11A, B). In contrast, H3K9me3 exhibited a markedly different regulatory organization by demarcating broad cellular lineages and silencing genes incompatible with a given cellular identity (Figure 5E). H3K9me3-enriched domains encompassed sensory receptor clusters, developmental regulators, metabolic genes, and neuronal lineage determinants within non-neuronal populations, consistent with constitutive repression of lineage-inappropriate transcriptional programs. Conversely, genes defining major neuronal, vascular, and immune lineages, including *Bcl11b*, *Satb2*, *Tbr1*, *Emx2*, *Neurod1*, and numerous endothelial and immune regulators, were selectively depleted of H3K9me3 within their corresponding cellular classes, permitting lineage-specific gene expression (Figure 5E).

These complementary patterns suggest that H3K27me3- and H3K9me3-associated repression programs operate at distinct regulatory scales. H3K27me3 preferentially marks developmental regulators to distinguish closely related cellular identities, whereas H3K9me3 establishes broader lineage boundaries by stably silencing incompatible transcriptional programs. Together with active chromatin states, these distinct repressive mechanisms provide a hierarchical regulatory framework for understanding how multiple chromatin states contribute to the specification and maintenance of brain cell identity.

### Sex-dependent histone modification features and gene expression in brain cell types

We investigated sex differences in histone mark deposition and in gene expression patterns across brain cell subclasses (Table S6). Sex chromosomes were excluded from differential histone modification peak analysis because they differ in copy number and are subject to dosage compensation, X-chromosome inactivation, and extensive Y-chromosome heterochromatin. Across most subclasses, fewer than 100 autosomal differential peaks were identified for any histone modification (Figure S12A, left), indicating broadly similar autosomal epigenomic landscapes between males and females. A subset of subclasses showed more than 100 differential peaks for at least one histone modification, including 006 L4/5 IT CTX Glut, 007 L2/3 IT CTX Glut, 008 L2/3 IT ENT Glut, 014 LA-BLA-BMA-PA Glut, 022 L5 ET CTX Glut, several striatal GABAergic subclasses, and 318 Astro-NT NN (Figure S12A, left). We next performed differential gene expression analysis using the snRNA-seq profiles of the Paired-Tag dataset (Figure S12A, right). As expected, X-linked genes such as *Xist* and *Kdm6a* were consistently detected as the top sex-biased transcripts in nearly all subclasses (Figures S12B, C). Beyond these, most subclasses contained relatively few sex-differential genes, and the effect sizes were generally modest. Nevertheless, we identified genes whose sex-biased expression was accompanied by concordant active or repressive histone mark differences, including *Kcnc2* in 006 L4/5 IT CTX Glut, *St6galnac3* in 008 L2/3 IT ENT Glut, *Sptbn2* in 014 LA-BLA-BMA-PA Glut, and *Adarb2* in 063 STR D1 Sema5a Gaba (Figures S12D-G).

Because subclass-level aggregation may mask finer sex-associated cell states, we next examined sex imbalance among independently defined Paired-Tag L5 clusters beyond Allen Brain Cell atlas’ taxonomy framework. Testing each cluster against the overall dataset sex ratio identified a substantial number of female-biased clusters within our dataset that were robustly detected in both biological replicates (Figures S12H, I, Table S6). The only male-biased cluster identified in this analysis was mapped to 016 CA1-ProS Glut, whereas female-biased clusters were distributed across neuronal and non-neuronal subclasses (Figure S12J). STR D1/D2 GABA neurons from CPU and NAC and DG Glut neurons from HCa and HCp contained the largest numbers of sex-biased cells, with additional contributions from astrocytes, oligodendrocytes, OPCs, and microglia (Figure S12J). These data are consistent with previous studies showing that sex-related differences in the brain may involve both neuronal and glial components^133^. We further examined the top 2 most abundant female-specific L5 clusters (cluster 1-2-1-0-0/L5A, and cluster 1-1-1-0-0/L5B), which were mainly mapped to STR D2 Gaba and STR D1 Gaba subclasses, respectively (Figure S12H). In the L5A cluster, when compared with other cells in the STR D2 Gaba subclass, top autosomal upregulated genes were implicated in synaptic signaling, neural circuitry, and neurological traits and disorders^134–136^ (Figure S12K). In the L5B cluster, when compared with other cells in the STR D1 Gaba subclass, differentially expressed genes suggested altered GPCR, purinergic, and sphingolipid-related signaling, as well as cyclic nucleotide and extracellular matrix regulation^137–142^ (Figure S12L). Consistent with these transcriptional differences, differentially expressed genes in these female-specific clusters exhibited concordant changes in active and repressive histone marks at their genomic loci, providing cross-modality support for these distinct female-associated cell states captured by Paired-Tag (Figure S12M). Together, these analyses indicate that sex-associated differences in the adult mouse brain are generally modest at the level of subclass-aggregated autosomal chromatin and transcriptomic profiles, but become more apparent at finer Paired-Tag-defined cluster resolution. These results highlight sex-imbalanced neuronal and glial cell states that may underlie sex-dependent variation in the adult mouse brain epigenome and transcriptome.

### Active and repressive chromatin programs differentially regulate transposable elements

The preceding analyses demonstrate that active and repressive chromatin programs cooperate to establish cell-type-specific gene expression across the mammalian brain. We next asked how these complementary regulatory mechanisms operate within one of the most challenging components of the genome, transposable elements (TEs), which comprise approximately 37.5% of the mouse genome^143–146^. Although most TEs are epigenetically silenced through H3K9me3-mediated heterochromatin formation to preserve genome stability, accumulating evidence suggests that a small subset has been co-opted as lineage-specific regulatory elements^147^. The integrated chromatin-state framework provides a unique opportunity to distinguish these alternative regulatory fates across diverse brain cell types.

Consistent with their canonical role in genome defense, H3K9me3-associated heterochromatin (Hc-H) represented the predominant chromatin state across long interspersed nuclear elements (LINEs) and long terminal repeat (LTR) families, supporting widespread constitutive silencing of these TEs (Figure 3B). However, not all TEs were regulated equivalently. Using the cell-type-resolved epigenomic landscapes of the mouse brain, we first examined H3K27ac and H3K9me3 signals across genomic regions of each TE subfamily as proxies of their overall activation and repression states, and evaluated their associated H3K4me1 and H3K27me3 levels. This analysis identified two distinct modules of TE subfamilies that feature the highest average H3K27ac (active) and H3K9me3 (silenced) signals among brain cell subclasses, respectively, suggesting that different subfamilies of TEs are subject to distinct modes of regulation mediated by histone modifications (Figures 6A-D). Specifically, TE subfamilies in the H3K27ac-enriched module also exhibited elevated H3K4me1 signals (Figure 6B), consistent with features of enhancer-like, active chromatin states. In contrast, top TE subfamilies with the highest H3K9me3 signal showed uniformly low H3K27ac and H3K4me1 levels (Figures 6A, B, C), suggesting an H3K9me3-mediated constitutive heterochromatin state. Thus, the integrated chromatin-state atlas distinguishes two fundamentally different regulatory modes: pervasive H3K9me3-mediated repression of most repetitive elements and selective activation of a small subset with potential regulatory function.

**Figure 6.**
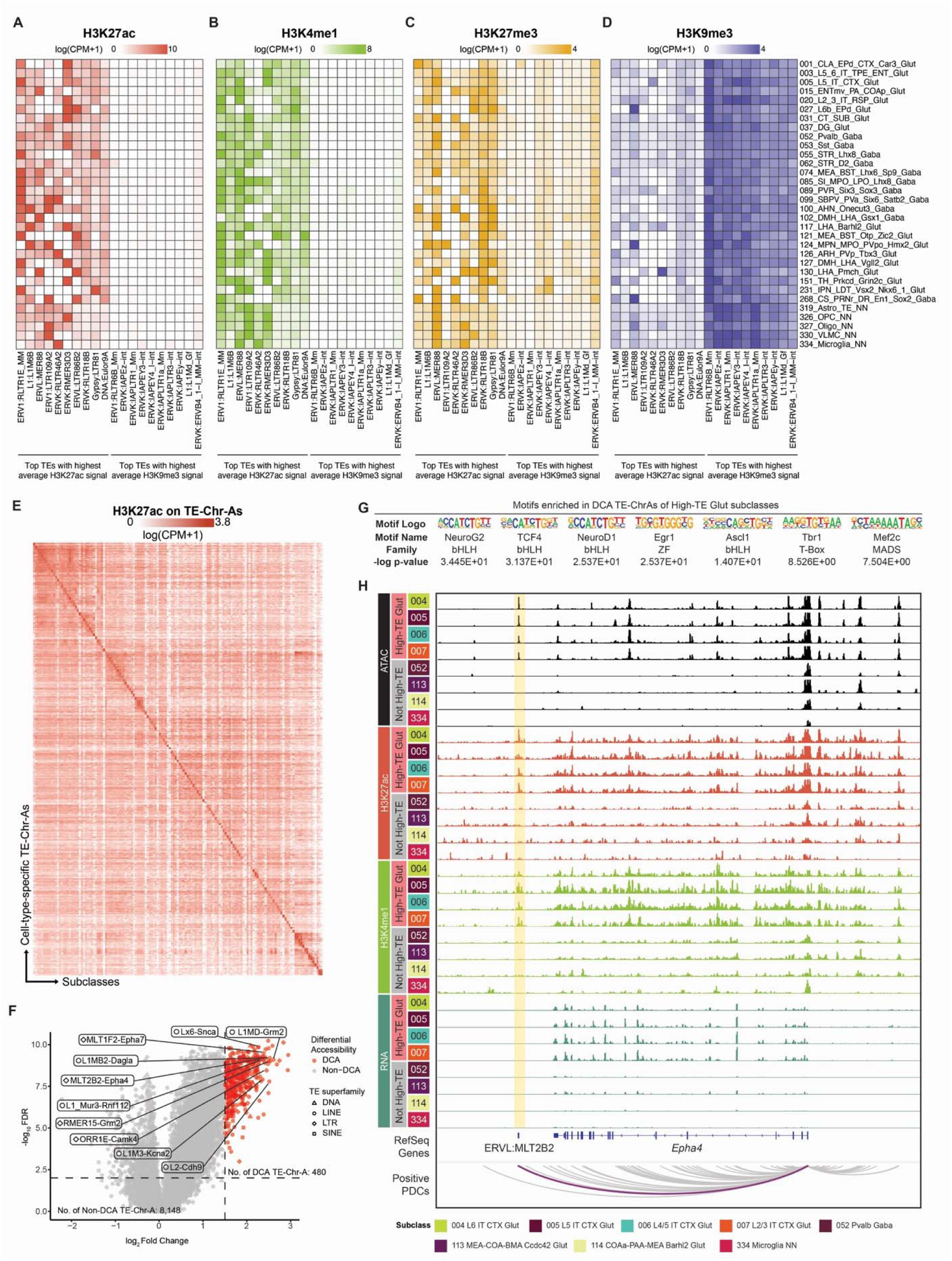
Transposable elements as potential gene regulatory elements. A–D. Heatmaps showing average H3K27ac (A), H3K4me1 (B), H3K27me3 (C), and H3K9me3 (D) signals across selected brain cell subclasses at the same set of transposable elements (TEs). TEs are grouped by those exhibiting the highest average H3K27ac or H3K9me3 signal across subclasses. E. Heatmap showing normalized H3K27ac signal at cell-type-specific TE-Chr-As across brain cell subclasses. F. Volcano plot showing differential chromatin accessible (DCA) TE-Chr-As in High-TE Glut subclasses compared with other subclasses. The top ten DCA TE-Chr-As correlated with synaptic-related genes are shown. G. Motifs enriched in DCA TE-Chr-As of High-TE Glut subclasses. Statistical significance was assessed using Fisher’s exact test. H. Genome browser tracks showing ATAC, H3K27ac, H3K4me1, and RNA signals for selected High-TE Glut subclasses (004 L6 IT CTX Glut, 005 L5 IT CTX Glut, 006 L4/5 IT CTX Glut, and 007 L2/3 IT CTX Glut) and representative subclasses that are not High-TE (052 Pvalb Gaba, 113 MEA-COA-BMA Ccdc42 Glut, 114 COAa-PAA-MEA Barhl2 Glut, and 334 Microglia NN), at the *Epha4* locus. An ERVL:MLT2B2 TE-Chr-A is highlighted, with specific ATAC, H3K27ac, and H3K4me1 signals in High-TE Glut subclasses, and a positive proximal-distal connection (PDC) to the *Epha4* transcriptional start site.

To investigate the functional significance of these active TE populations, we integrated chromatin-state annotations with previously identified TE-overlapping cCREs in the mouse brain (TE-cCREs) ^35^. TE-derived active regulatory elements (TE-Chr-A) displayed markedly stronger H3K27ac and H3K4me1 signals than TE-Chr-O elements (Figures S13A, B), and exhibited pronounced cell-type-specific H3K27ac patterns (Figure 6E, Table S7). Consistent with our previous observation that certain glutamatergic neuronal subclasses preferentially utilize TE-derived regulatory elements, which we refer to as High-TE Glut subclasses, these neurons contained substantially larger fractions of TE-Chr-A regions than other brain cell types (Figure S13C, D) ^35^. Differential chromatin accessibility analysis identified hundreds of TE-derived active regulatory elements enriched in these neuronal populations, with LINE/L1 families contributing prominently ^35^ (Figure 6F).

These active TE-derived regulatory elements also displayed the molecular characteristics expected of functional enhancers. Motif enrichment analysis on differential Chr-As of High-TE Glut subclasses revealed binding sites for neuronal lineage regulators, including NeuroG2, TCF4, NeuroD1, Egr1, Ascl1, Tbr1 and Mef2C (Figure 6G) ^99,102,148–153^, suggesting their involvement in neuronal transcriptional regulatory networks. Proximal-distal connection analyses further linked individual TE-derived enhancers to nearby neuronal genes^35^. For example, an MLT2B2-derived regulatory element located downstream of *Epha4* exhibited strong proximal-distal connection and cell-type-specific active chromatin signatures, consistent with a role in promoting preferential *Epha4* expression in High-TE Glut neurons (Figure 6H).

Together, these analyses demonstrate that active and repressive chromatin programs govern distinct regulatory fates of repetitive DNA. Whereas H3K9me3-mediated heterochromatin enforces widespread silencing of transposable elements to maintain genome integrity, a highly selected subset escapes constitutive repression, acquires active chromatin states, and becomes incorporated into lineage-specific gene regulatory networks. These findings extend the chromatin-state framework established throughout this study by showing that the same regulatory principles governing conventional enhancers also explain how repetitive genomic sequences are selectively repurposed as functional regulatory elements in the mammalian brain.

### Multimodal epigenomic atlases enable the prediction of regulatory programs from DNA sequence

Finally, we asked whether the regulatory programs revealed by our integrative epigenome atlas can be learned and predicted from DNA sequence using deep learning. Recent advances in sequence-to-function (S2F) modeling have demonstrated remarkable progress in predicting individual epigenomic features from genomic sequence^154,155^, interpreting both coding and non-coding variation^156–158^, and designing functional DNA sequences *de novo*^159,160^. We reasoned that the integrated multimodal atlas generated here provides an unprecedented training resource for learning the regulatory logic underlying cell-type-specific gene regulation.

To test this idea, we fine-tuned the Borzoi S2F framework^155^ using approximately 1,400 transcriptomic and epigenomic tracks spanning more than 130 brain cell subclasses and seven complementary epigenomic modalities along with stranded gene expression (Figure 7A). Unlike previous models trained primarily on chromatin accessibility or bulk functional genomic data, our framework simultaneously learns multiple layers of chromatin regulation, including active and repressive histone modifications, chromatin accessibility, DNA methylation, and gene expression, from a unified single-cell reference atlas.

**Figure 7.**
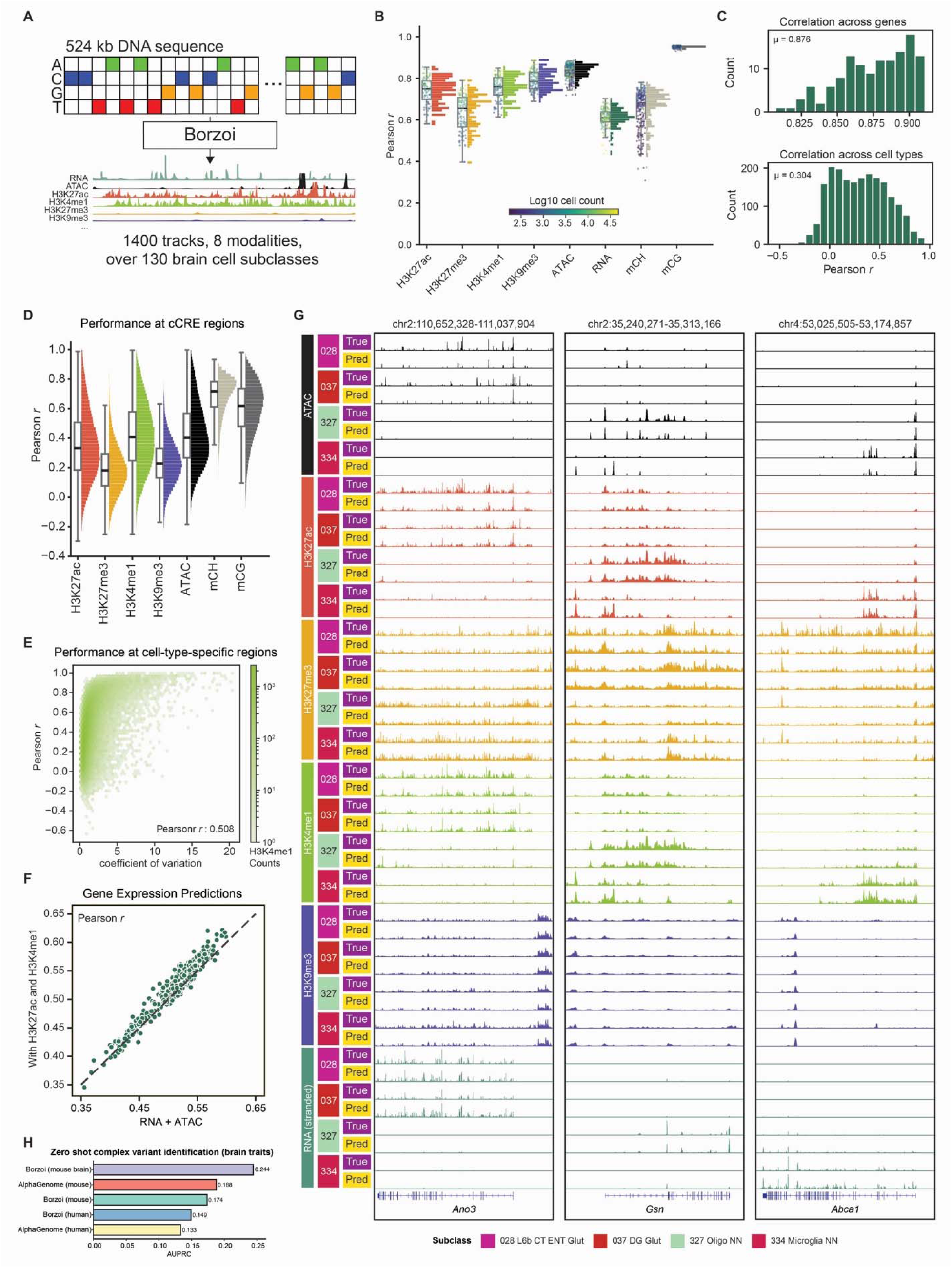
Deep learning prediction of brain epigenomic features and gene expression from DNA sequences. A. Schematic of the mouse brain Borzoi model, which takes as input 524 kilobases of DNA sequence and predicts 1400 tracks composed of 8 modalities in over 130 brain cell subclasses. Tracks are predicted as the average over every 32 bp. B. Model performance in an unseen test set for each modality was measured by the Pearson correlation between predicted values and measured values for each subclass. Boxplots show the mean, interquartile range, and data extent of Pearson correlation across subclasses. Histograms show the distribution of correlation values, with point clouds highlighting individual subclasses. Each point is colored by the number of cells measured in each subclass. C. Top, histogram of the Pearson correlation between predicted expression and measured expression across all genes in the test set for each subclass. Bottom, histogram shows the Pearson correlation between the predicted expression pattern across cells and the measured expression pattern across cells for each gene in the unseen test set. D. Model performance in the unseen test set predicting patterns of epigenomes across cell types at mouse brain cCRE regions^35^. Boxplots show mean, interquartile range, and data extent across cCRES with histograms showing the distribution. E. Model performance across cCREs with different levels of variability. Pearson R measures the model performance predicting patterns of H3K4me1 across cells, while the coefficient of variation measures the specificity of measured H3K4me1 across subclasses. F. A comparison of gene expression prediction with ATAC alone or when adding H3K27ac and H3K4me1 when training from scratch. Points show the correlation between predicted and measured tracks in an unseen test set, with the x-axis position showing the value for a model trained with ATAC alone, and the y-axis position showing the value for a model trained with active histone modifications as well. G. Genome browser tracks showing predicted and measured ATAC, H3K27ac, H3K27me3, H3K4me1, H3K9me3, and RNA signals across representative brain cell subclasses (028 L6b CT ENT Glut, 037 DG Glut, 327 Oligo NN, and 334 Microglia NN) at selected cell-type-specific genes (*Ano3*, *Gsn*, and *Abca1*) in the unseen test set. H. Zero-shot performance of the mouse brain fine-tuned Borzoi model developed in this study on all neurological-trait variants in the TraitGym benchmark, compared with AlphaGenome mouse and human models and untuned Borzoi mouse and human models.

The resulting models predicted cell-type-specific epigenomic landscapes directly from DNA sequence. Across held-out genomic regions, predicted epigenomic profiles largely matched experimentally measured chromatin states, with particularly strong performance for DNA methylation and active chromatin features (Figure 7B). We also identified prediction accuracy to be dependent on cell number, which serves as a proxy for evaluating genome coverage and data quality (Figures S13E-G). While RNA coverage was more challenging to predict than other epigenomic modalities, our model accurately predicted the relative gene expression within individual cell types, with an average Pearson correlation of 0.876 (Figure 7C). Predicting gene expression differences across cell types was more challenging, with an average correlation per gene of 0.3 (Figure 7C). This behavior likely reflects a bias in model training to attend to highly expressed genes, which incur larger loss penalties, given that we found the model’s ability to predict expression differences across cells is highly correlated with the expression of a gene in its most highly expressed cell type (Figure S13H).

To assess our model’s ability to predict cell-type-specific regulatory elements, we measured the correlation between our model’s predicted epigenomes and the measured epigenomes at regions of our previously defined brain cCREs (Figure 7D)^35^. Our model demonstrated exceptional performance in predicting DNA methylation differences across cell types. While predictions for active histone modifications were generally robust, our model’s performance was reduced for predicting differences in repressive histone marks, likely due to lower biological variability and signal intensity (Figure S13I). For active histone marks, we found that our model’s ability to predict epigenome patterns at peaks across cell types was highly correlated with their variability (Figure 7E, Figure S13J).

Importantly, models trained from scratch using active histone modifications together with chromatin accessibility outperformed models trained using chromatin accessibility alone in predicting RNA expression, demonstrating that complementary epigenomic features provide richer information about gene regulation than individual modality (Figure 7F). To demonstrate the interpretability of our model’s predictions, we visualized representative genomic loci encoding *Ano3*, *Gsn*, and *Abca1*, and demonstrated that predicted epigenomic landscapes and gene expression closely recapitulated experimentally observed cell-type-specific regulatory patterns. These data suggest that our model successfully captured coordinated relationships among multiple layers of epigenomic regulation and their impacts on the transcriptional output (Figure 7G).

We further evaluated whether our brain-specific model could generalize to independent assessments of noncoding variant function using the TraitGym benchmark, which includes fine-mapped, high-probability causal variants paired with matched control variants with similar properties^156^. In this zero-shot analysis, when restricted to neurological traits, our model outperformed Borzoi and AlphaGenome models trained on mouse or human seuqnece with no additional tuning (Figure 7H). In contrast, across the full benchmark, AlphaGenome showed the strongest overall performance, whereas our brain-specific model showed a decreased ability to detect causal variants, likely reflecting the small contribution of neurological traits within the full benchmark (Figure S13K). To facilitate broad access to these models, we provide an interactive online predictor that predicts subclass-specific epigenomes and transcriptomes for arbitrary DNA sequences at https://seqnn.org.

Together, these findings demonstrate that our integrative epigenomic atlas serves not only as a comprehensive reference for genome annotation but also as a powerful training resource for predictive models of gene regulation. By learning coordinated patterns of enhancer activation, chromatin repression, DNA methylation, and transcription from DNA sequence, these models move beyond predicting individual molecular features toward learning the regulatory grammar that specifies cell-type identity. This framework establishes an important step toward predictive interpretation of noncoding variation and rational engineering of cell-type-specific regulatory elements.

## DISCUSSION

The remarkable cellular diversity of the mammalian brain arises from precisely coordinated gene regulatory programs shaped by multiple layers of epigenomic regulation. A longstanding challenge has been to understand how these regulatory layers act together to establish and maintain cell identity. In this study, we address this challenge by constructing an integrated single-cell epigenomic atlas that combines transcription, four histone modifications, chromatin accessibility, DNA methylation, and three-dimensional genome organization into a unified regulatory framework spanning more than 100 brain cell subclasses^6,34,35^.

A central finding of this study is that cell-type-specific regulatory information emerges from the integration of multiple chromatin modalities rather than from any single epigenomic layer. Whereas chromatin accessibility catalogs cCREs, integrating active and repressive histone modifications resolves their functional states, allowing active, primed, poised, and repressed chromatin states to be distinguished across the majority of the mouse brain genome. This expanded annotation coverage suggests that cell-type identity is encoded not only through selective activation of regulatory elements, but also through widespread deployment and resolution of repressive chromatin programs.

Our analyses also clarify the distinction between open chromatin and active regulatory elements. Chr-A regions, defined by chromatin accessibility together with H3K27ac and H3K4me1, exhibit greater evolutionary conservation, stronger enrichment for validated brain enhancers, higher cell-type specificity, enhancer RNA transcription, and stronger lineage TF motif enrichment than Chr-O regions. In contrast, Chr-O regions were enriched for CTCF-related motifs, consistent with architectural roles in genome organization. These findings suggest that chromatin accessibility should no longer be viewed as a sufficient proxy for enhancer activity. Rather, enhancer-associated regulatory activity is best represented by combinatorial chromatin states that integrate accessibility with active histone modifications. Although the integrative framework presented here prioritizes candidate regulatory elements using multiple orthogonal lines of evidence, direct perturbation of candidate enhancers and experimental validations of predicted regulatory interactions will be important next steps. The approximately half million active chromatin regions identified in this study therefore provide a rich, cell-type-resolved resource of candidate enhancers for future high-throughput validation of regulatory activity and cell-type specificity, as well as mechanistic characterization through perturbation-based approaches.

Our analyses further highlight distinct and complementary roles for different repressive chromatin programs in shaping brain cell identity, particularly those marked by H3K27me3 and H3K9me3. H3K27me3-mediated repression refines closely related cellular identities by selectively silencing developmental regulators, whereas H3K9me3 demarcates broader lineage boundaries through constitutive repression of incompatible gene programs. Together, these complementary mechanisms provide a hierarchical framework for stabilizing cellular identity. Our atlas also provides a framework for investigating sex-dependent chromatin states that may contribute to sexually dimorphic brain function and disease susceptibility.

Finally, the scale and diversity of the atlas enabled sequence-based deep learning models that predict cell-type-specific epigenomic features and gene expression from DNA sequence. By learning multiple layers of chromatin regulation simultaneously, these models move beyond predicting individual epigenomic features toward learning the regulatory grammar that links DNA sequence to cell-type-specific gene regulation. As larger multimodal atlases become available across tissues and species, this framework should facilitate increasingly accurate interpretation of noncoding variation and rational design of regulatory DNA. That said, the models presented in this work should be interpreted within the scope of the training atlas: they capture sequence-encoded regulatory potential but cannot directly infer expression changes driven by altered trans-acting environments, such as changes in TF abundance, signaling state, aging, or disease when DNA sequence is unchanged. Looking forward, applying emerging foundation models and newer sequence-based architectures to this atlas may further improve predictive accuracy and reveal additional regulatory information from these multimodal profiles.

In summary, this work illustrates a transition in functional genomics from descriptive cataloging of regulatory elements toward predictive modeling of genome function. By integrating complementary layers of epigenomic regulation at single-cell resolution, our study provides both a comprehensive annotation of the mammalian brain regulatory genome and a foundation for learning the regulatory logic that connects DNA sequence to cell identity. We anticipate that similar multimodal reference atlases across human tissues and disease states will accelerate interpretation of noncoding genetic variation, enable increasingly precise engineering of cell-type-specific regulatory elements, and ultimately support predictive models of genome function in health and disease.

### Limitations of the study

Our study provides a comprehensive multimodal epigenomic atlas of the adult mouse brain, but several limitations should be considered. First, although integration of multiple epigenomic modalities enables functional annotation of regulatory elements and inference of gene regulatory programs, most regulatory relationships remain correlative and will require perturbation-based validation to establish causality. Second, our atlas represents the healthy adult brain and therefore does not capture the dynamic chromatin remodeling that occurs during brain development, aging, or neurological disease. Extending this framework across developmental stages and disease models will provide important insights into the regulatory mechanisms underlying brain function and dysfunction. Third, while the atlas integrates transcriptomes with four histone modifications, chromatin accessibility, DNA methylation, and three-dimensional genome organization, additional regulatory layers, including transcription factor occupancy, RNA modifications, chromatin-associated proteins, and spatially resolved epigenomic measurements, will further refine our understanding of cell-type-specific gene regulation. Fourth, while reference-based annotation enables robust integration with existing brain cell atlases and multimodal resources, it may bias analyses toward previously defined cellular identities. To mitigate this limitation, we additionally incorporated an unbiased iterative clustering analysis of Paired-Tag snRNA-seq data, which captures fine-grained cellular heterogeneity beyond the current reference taxonomy for future characterization. Fifth, although our atlas includes biological replicates from both sexes, the limited number of pooled replicates may constrain sample-level inference and reduce power to detect subtle sex-dependent differences. Finally, although the predictive models developed here demonstrate the potential of multimodal atlases for sequence-to-function learning, improving prediction accuracy for rare cell types and repressive chromatin states will likely require larger datasets and continued advances in model architecture.

---

**Figure S1.**
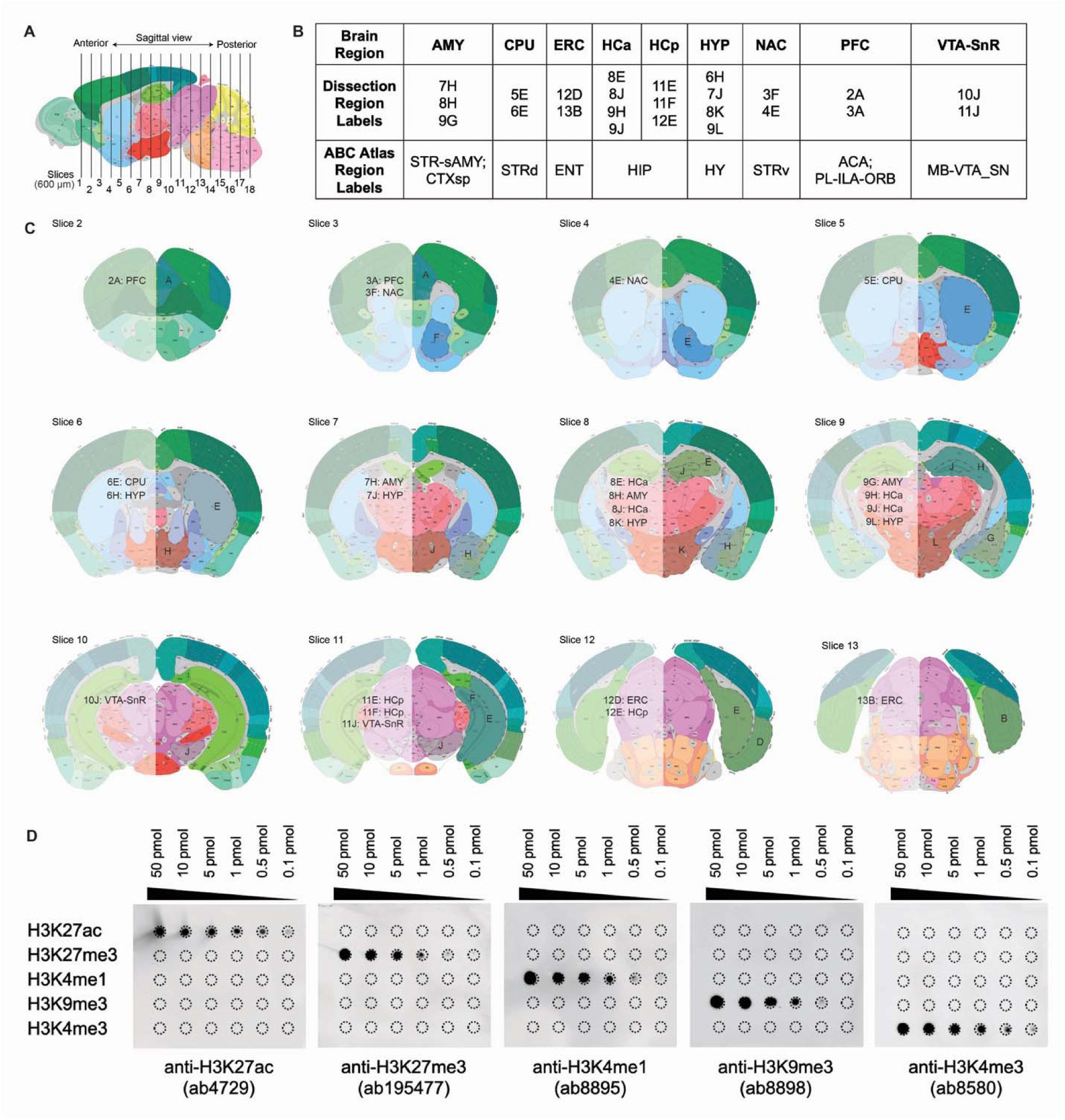
Sample collection strategy, related to Figure 1. A. Schematic overview of the mouse brain tissue dissection strategy. Mouse brains were sectioned into 600 µm thick coronal slices. B. Brain regions analyzed in this study, their corresponding dissection region labels, and the matched brain region annotations registered in the Allen Brain Cell Atlas. The dissection region labels used here are consistent with those defined in prior work from the Center for Epigenomics of the Mouse Brain Atlas (CEMBA), generated as part of the Brain Research through Advancing Innovative Neurotechnologies (BRAIN) Initiative - Cell Census Network (BICCN). C. Brain regions dissected from each coronal slice, annotated according to the Allen Brain Reference Atlas. The frontal view of slices 2-13 are shown, with the dissected region labels indicated on the left, and corresponding anatomical region annotations on the right. D. Dot blot assays demonstrating the specificity and reactivity of antibodies used in this study against recombinant histone H3 carrying various histone modifications. H3K4me3 antibody and recombinant peptides were used here as control. All brain maps shown in this figure were generated using coordinates from the Allen Mouse Brain Common Coordinate Framework (CCF) v3^71^.

**Figure S2.**
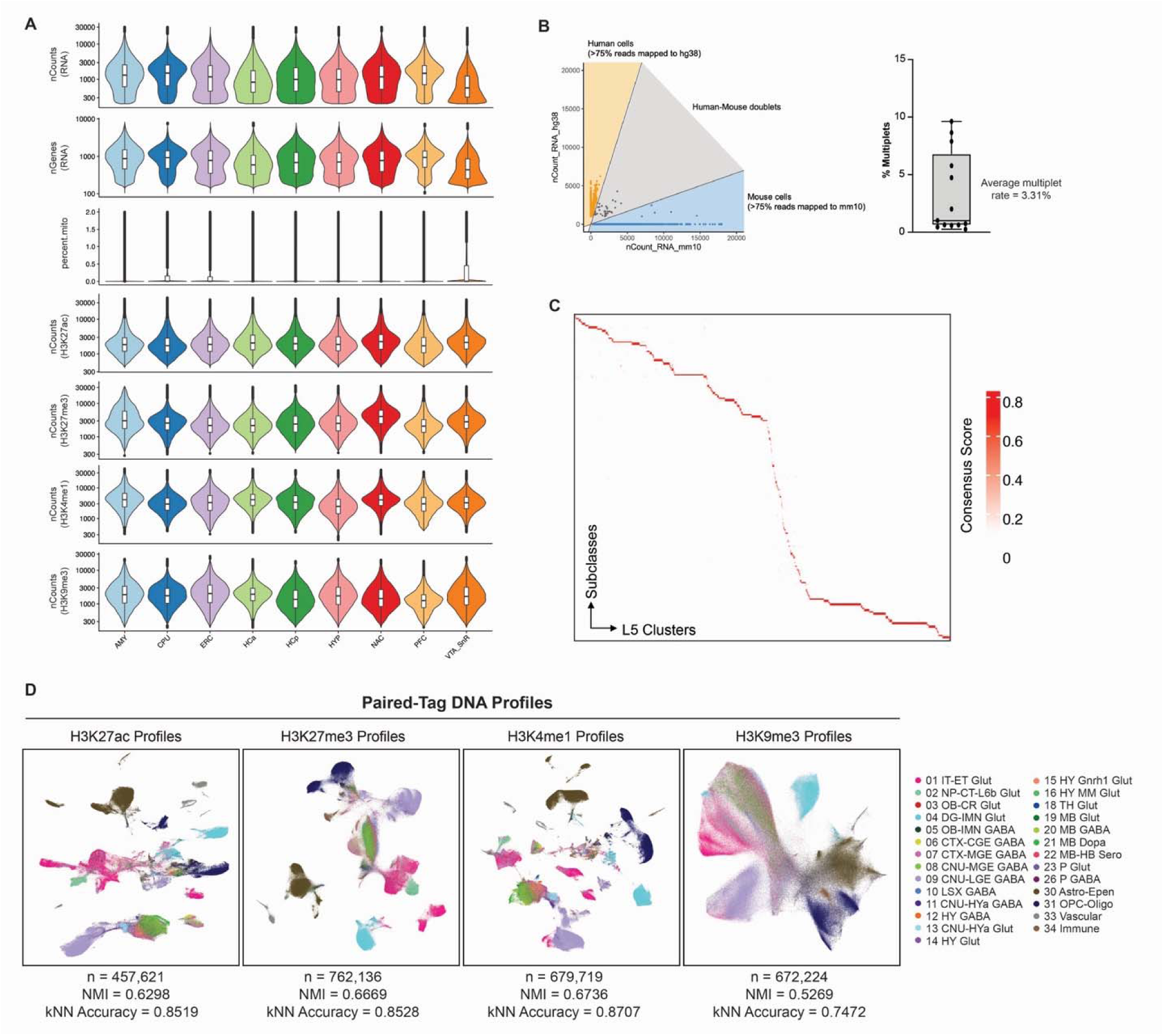
Quality control metrics, related to Figure 1. A. Violin plots, from top to bottom, showing per-cell quality of number of RNA unique read counts, number of genes detected, percentage of mitochondrial reads, number of H3K27ac unique read counts, number of H3K27me3 unique read counts, number of H3K4me1 unique read counts, and number of H3K9me3 unique read counts, for each brain region analyzed. B. Left, scatter plot showing the fraction of human and mouse RNA reads in each cell from the species-mixing experiment. Barcodes with fewer than 75% of reads from a single species were classified as doublets. Right, box plot showing the distribution and mean multiplet rate of the Paired-Tag dataset. C. Heatmap showing the consensus scores between scRNA-seq subclasses and L5-level clusters derived from Paired-Tag RNA profiles. D. Uniform manifold approximation project (UMAP) visualization of single cell Paired-Tag DNA profiles for each histone modification, from left to right: H3K27ac, H3K27me3, H3K4me1, and H3K9me3, colored by class labels. Normalized mutual information (NMI) and k-nearest neighbor label transfer accuracy values were used to evaluate clustering quality, with the Paired-Tag RNA modality achieving an NMI value of 0.7692 (cap).

**Figure S3.**
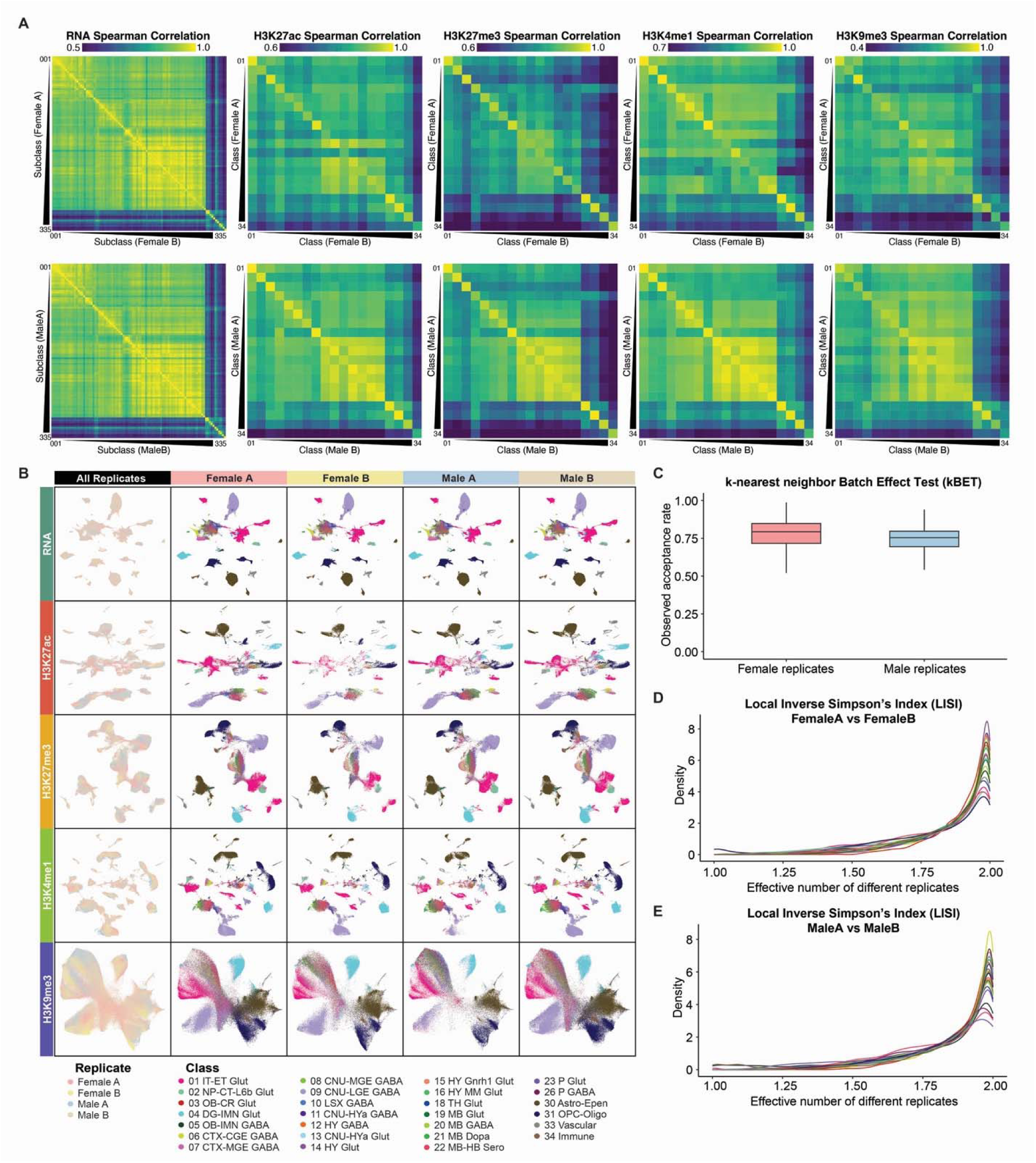
Reproducibility and batch effect assessment across biological replicates. A. Pseudobulk Spearman correlation heatmaps for Paired-Tag RNA-seq and histone modification profiles across female and male biological replicates. RNA correlations were computed at the subclass level, whereas histone modification correlations were computed at the class level. B. Joint UMAP visualizations of Paired-Tag RNA-seq and histone modification profiles from all cells, shown for all replicates together and colored by biological replicate, with corresponding panels for each individual replicate colored by class annotation, to assess the reproducibility of low-dimensional manifold structure across biological replicates. C. k-nearest neighbor Batch Effect Test (kBET) results for female and male biological replicates in the RNA modality. Median observed acceptance rates exceeded 0.75 in both groups, suggesting limited batch effects across replicates. D, E. Distributions of Local Inverse Simpson’s Index (LISI) scores for cells from each class, comparing two female replicates (D) and two male replicates (E). Across classes, LISI values were enriched near 2, indicating effective mixing of the two biological replicates for each sex.

**Figure S4.**
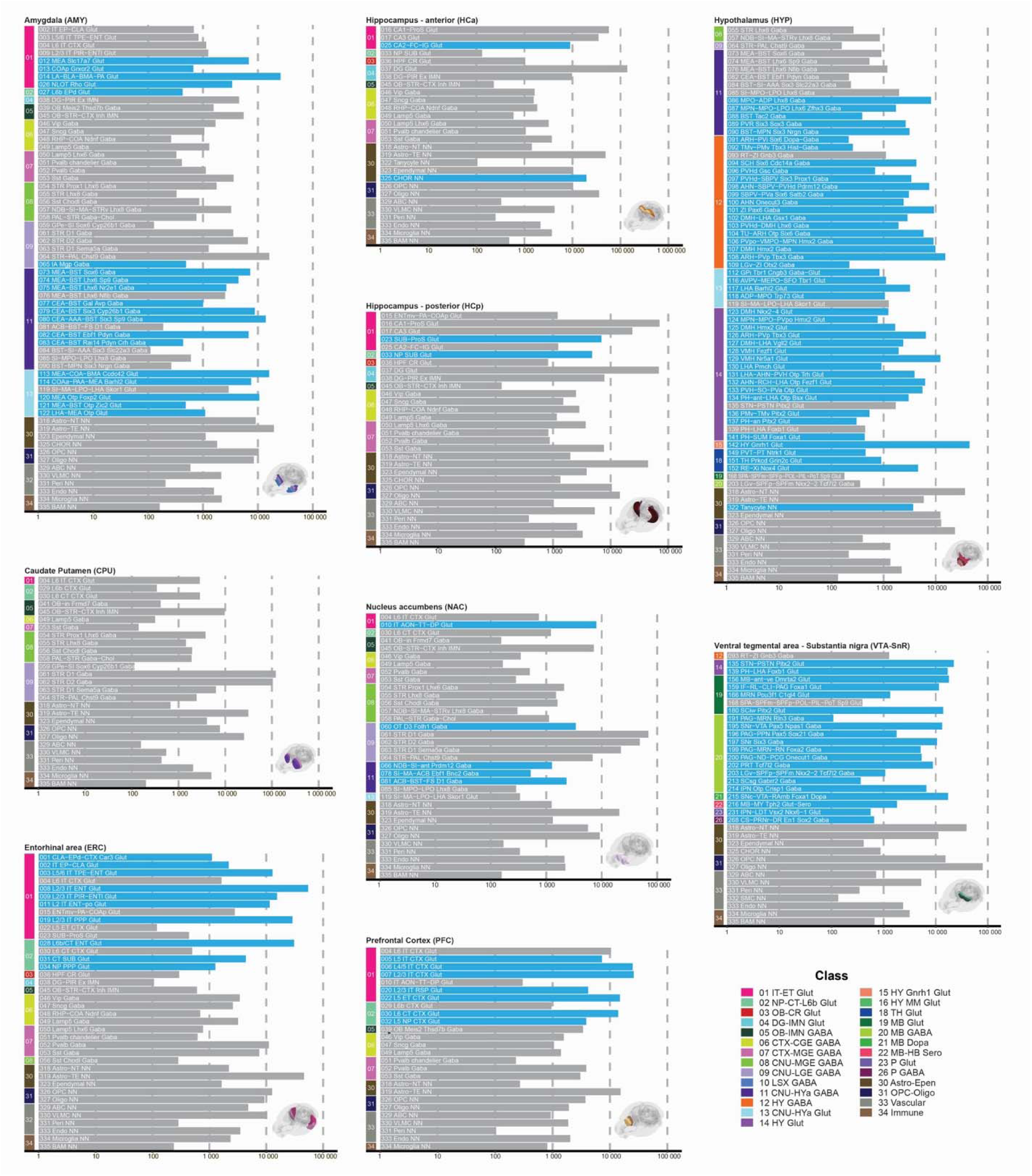
Cellular composition of the analyzed brain regions, related to Figure 1. Bar plots depict the cellular composition of the nine analyzed brain regions. Subclasses with over 100 cells for each region are shown. Region-specific subclasses, defined as subclasses for which more than 75% of cells in the Paired-Tag dataset originate from a single dominant brain region, are shown in blue. Brain region schematics were generated using coordinates from the Allen Mouse Brain Common Coordinate Framework (CCF) v3^71^.

**Figure S5.**
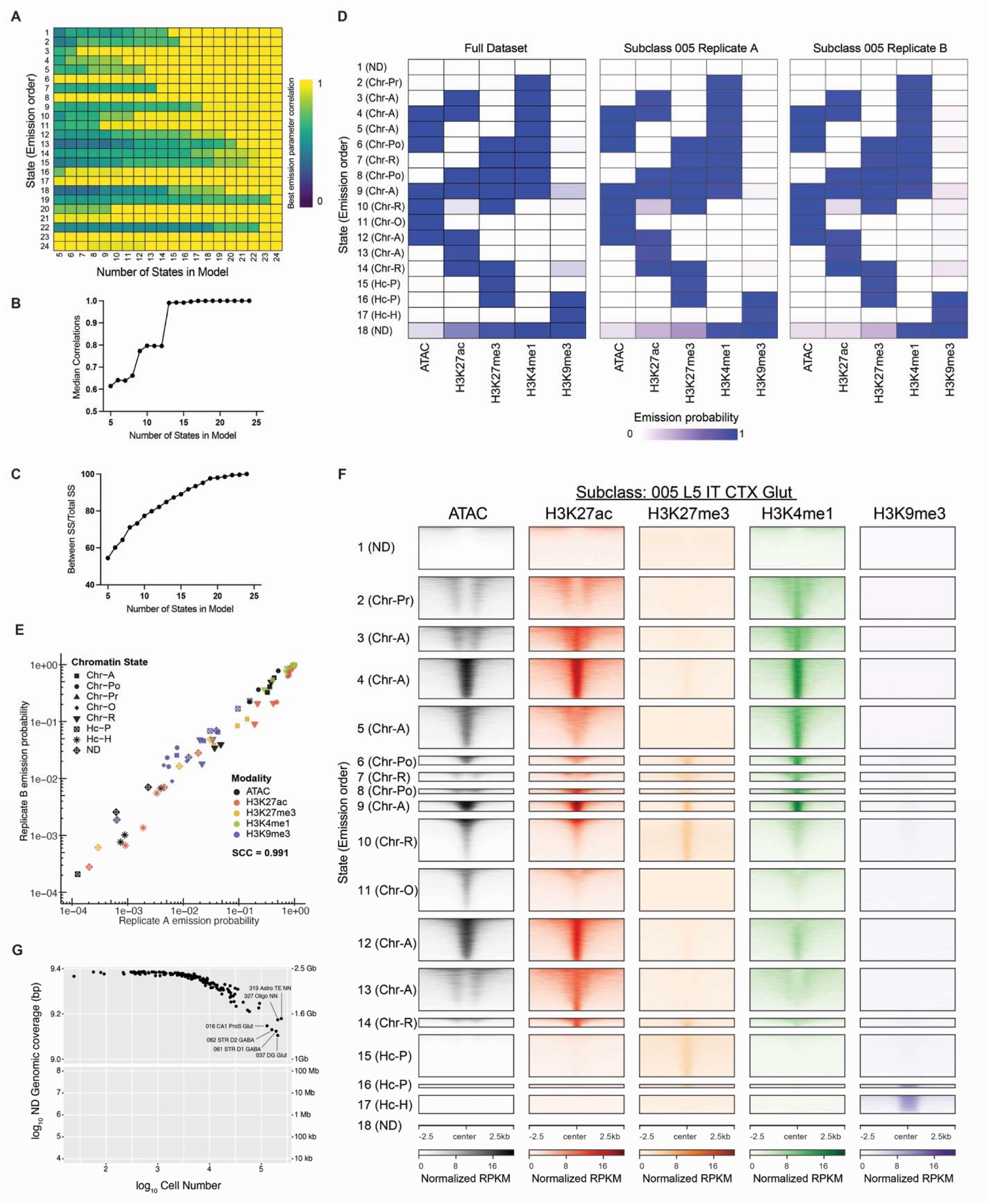
ChromHMM modeling of chromatin states, related to Figure 3. A. Heatmap showing the maximum Pearson’s correlation of each state in the full ChromHMM model (y-axis) with its best-matching state in each reduced model (x-axis). B. Median Pearson correlation of all 24 states in each reduced ChromHMM model (x-axis). Models with 18 states and more showed a median Pearson correlation of > 99%. C. Evaluation of k-means clustering using the ratio of between-cluster variance to total variance (Between SS/Total SS) across ChromHMM models with varying numbers of states. Models with 18 states and more showed a Between SS/Total SS of > 95%. D. Emission probabilities for each chromatin mark across the 18 states in the ChromHMM model, shown for the full dataset (left), replicate A (middle) and replicate B (right). Subclass 005 L5 IT CTX Glut is shown as an example to illustrate the consistency of state-specific emission profiles between biological replicates. E. Spearman correlation of emission probabilities from ChromHMM models derived from two replicates. Emission probabilities from three representative subclasses (005 L5 IT CTX Glut, 318 Astro-NT NN, and 319 Astro-TE NN) are shown. Dot colors indicate epigenomic modalities, and dot shapes indicate chromatin states. F. Coverage heatmap showing the ATAC, H3K27ac, H3K27me3, H3K4me1, and H3K9me3 signals from the 005 L5 IT CTX Glut subclass across states in the 18-state ChromHMM model. G. Relationship between genomic coverage of Not Determined (ND) chromatin state and cell number. Selected high coverage cell subclasses are labeled.

**Figure S6.**
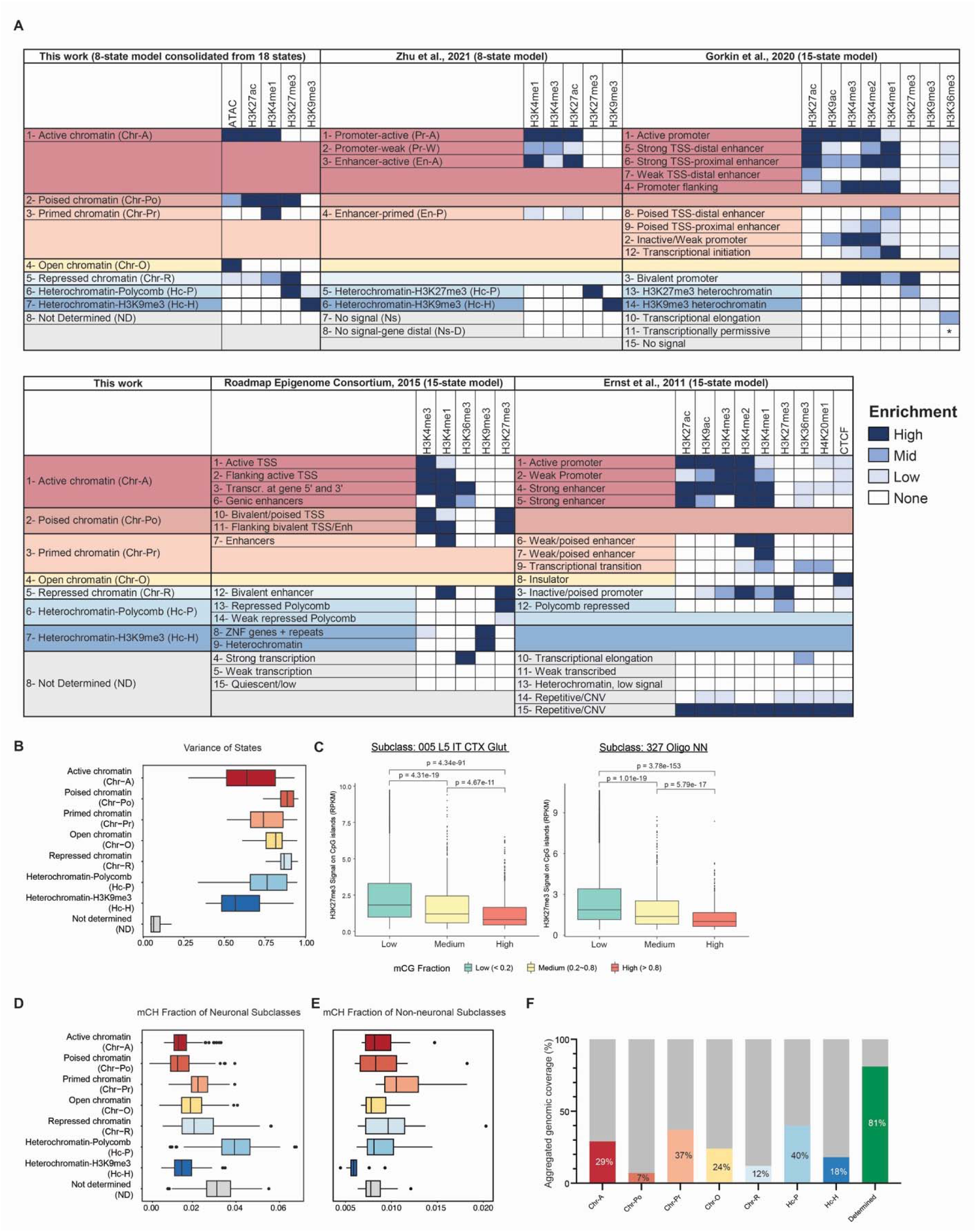
Characterization of the 8-state chromatin state model, related to Figure 3. A. Comparison of the ChromHMM model from this study with representative published ChromHMM models. Published chromatin states were aligned to the most similar state category in this work on the basis of relative enrichment patterns of profiled epigenomic marks. Because different studies used different combinations of chromatin features, these correspondences should be interpreted as approximate functional matches rather than one-to-one equivalences. Notably, our model incorporates chromatin accessibility and therefore identifies an open chromatin state (Chr-O) that is not explicitly represented in most previous models. In cases where no confident correspondence could be assigned, states were associated with the closest higher-level category or grouped with the Not Determined state for visualization purposes. *, in the 15-state ChromHMM model by Gorkin et al., 2020, although the H3K36me3 enrichment in state “11 - Transcriptionally permissive” is classified as “None”, the authors report that it was >30-fold higher than genomic background (state “15 - No signal”)^54^. B. Box plot showing fraction of variable bases associated with each chromatin state across brain cell subclasses. C. Box plots showing H3K27me3 signal on CpG islands stratified by Low (< 0.2), Medium (0.2∼0.8) and High (> 0.8) mCG fraction in representative brain cell subclasses 005 L5 IT CTX Glut (left) and 327 Oligo NN (right). Statistical testing was performed using *t* test followed by Benjamini-Hochberg correction. D. Box plot showing fraction of DNA non-CG methylation (mCH) associated with each chromatin state across neuronal brain cell subclasses. E. Box plot showing fraction of DNA non-CG methylation (mCH) associated with each chromatin state across non-neuronal brain cell subclasses. F. Bar plot showing aggregated genomic coverage of each chromatin state, along with genomic regions assigned functional chromatin state annotations, across all brain cell subclasses.

**Figure S7.**
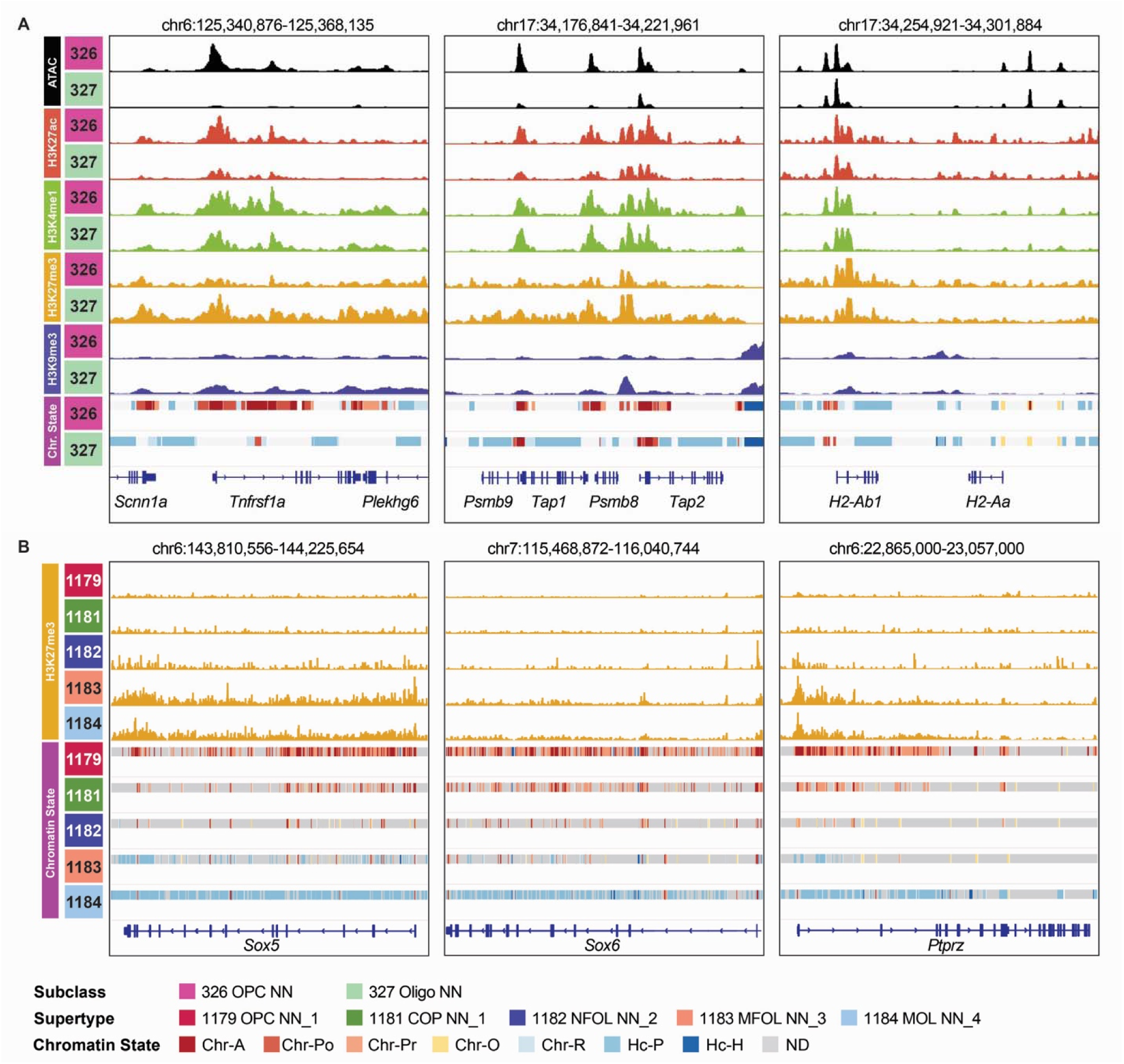
Epigenomic landscapes of previously reported immune and differentiation-associated loci in the oligodendrocyte lineage, related to Figure 3. A. Genome browser tracks showing chromatin accessibility, histone modification profiles, and chromatin state annotations at representative immune-related loci, including *Tnfrsf1a*, the *Psmb9 - Tap2* MHC-I-associated locus, and the *H2-Ab1/H2-Aa* MHC-II-associated locus, in subclasses 326 OPC NN and 327 Oligo NN. B. H3K27me3 signal and chromatin-state annotations across oligodendrocyte lineage supertypes at canonical OPC-associated genes, including *Sox5*, *Sox6*, and *Ptprz1*.

**Figure S8.**
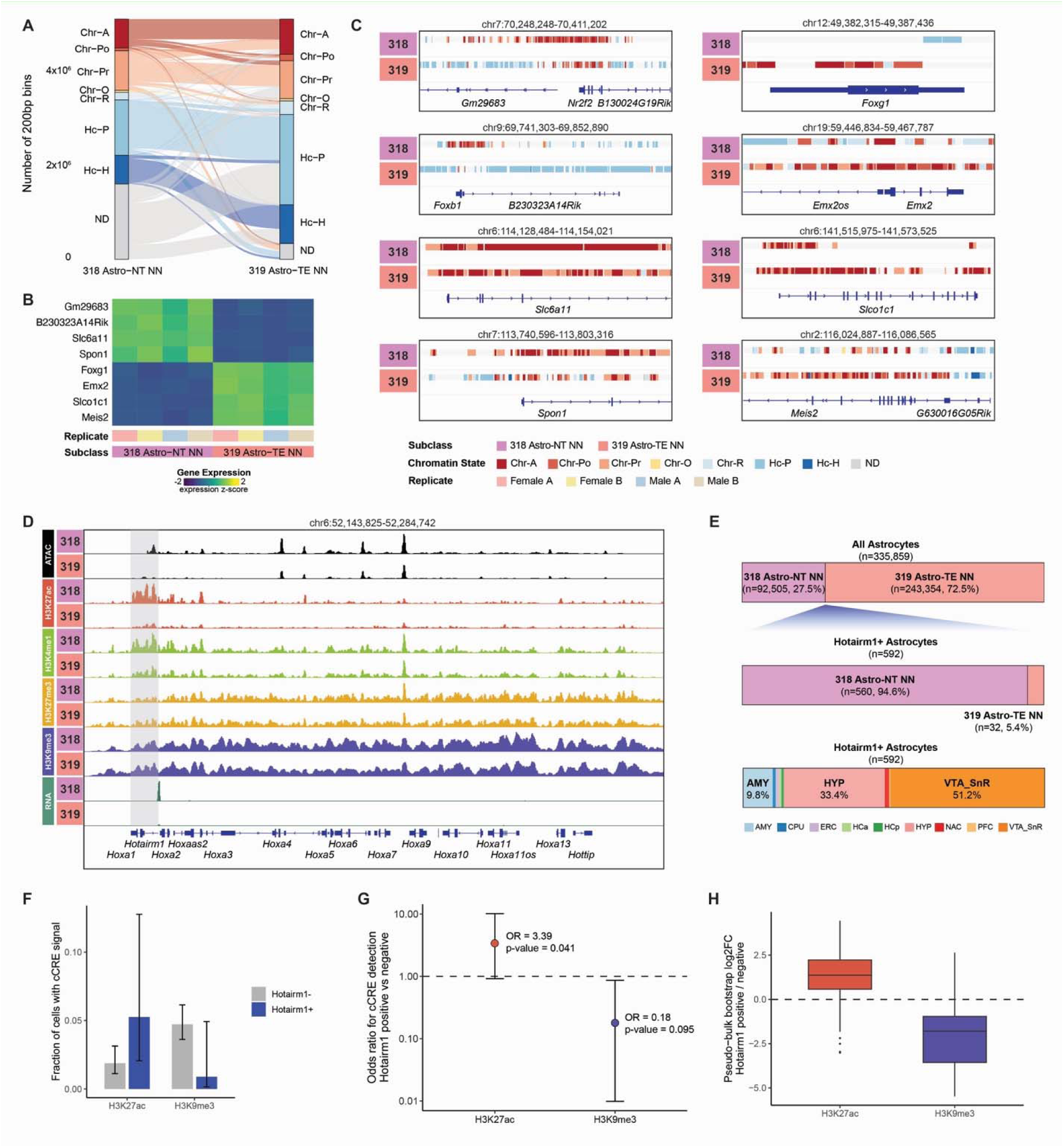
Chromatin-state differences between telencephalic and non-telencephalic astrocytes, related to Figure 3. A. Alluvial plot showing genome-wide chromatin state differences between Astro-NT and Astro-TE. Chromatin regions that are annotated as ND for both 318 Astro-NT NN and 319 Astro-TE NN subclasses are not shown. B. Heatmap showing scaled expression of representative Astro-NT and Astro-TE enriched genes across biological replicates. C. Chromatin state annotations at representative subclass-enriched genes in Astro-NT (left) and Astro-TE (right). D. Genome browser tracks showing chromatin accessibility, histone modification signals, RNA expression, and chromatin state annotations at the HoxA locus in Astro-NT and Astro-TE. A candidate Astro-NT-specific regulatory region at the 3’ end of the locus is highlighted. E. Distribution of *Hotairm1*-expressing astrocytes across subclasses and brain regions. The top bar shows subclass composition of all astrocytes. The middle bar shows *Hotairm1*-expressing astrocytes by subclasses. The bottom bar shows the brain region composition of Hotairm1-expressing astrocytes. F. Fraction of *Hotairm1*-positive and matched *Hotairm1*-negative astrocytes with H3K27ac or H3K9me3 signal at the candidate regulatory region. Error bars indicate Wilson confidence intervals. G. Logistic regression analysis testing the association between *Hotairm1* detection and histone modification detection at the candidate regulatory region at the single cell level. Points indicate odds ratios, and error bars indicate confidence intervals. The dashed line indicates an odds ratio of 1. H. Pseudobulk comparison of H3K27ac and H3K9me3 signal at the candidate regulatory region between *Hotairm1*-positive and matched *Hotairm1*-negative astrocyte groups. Values are shown as log2 fold changes.

**Figure S9.**
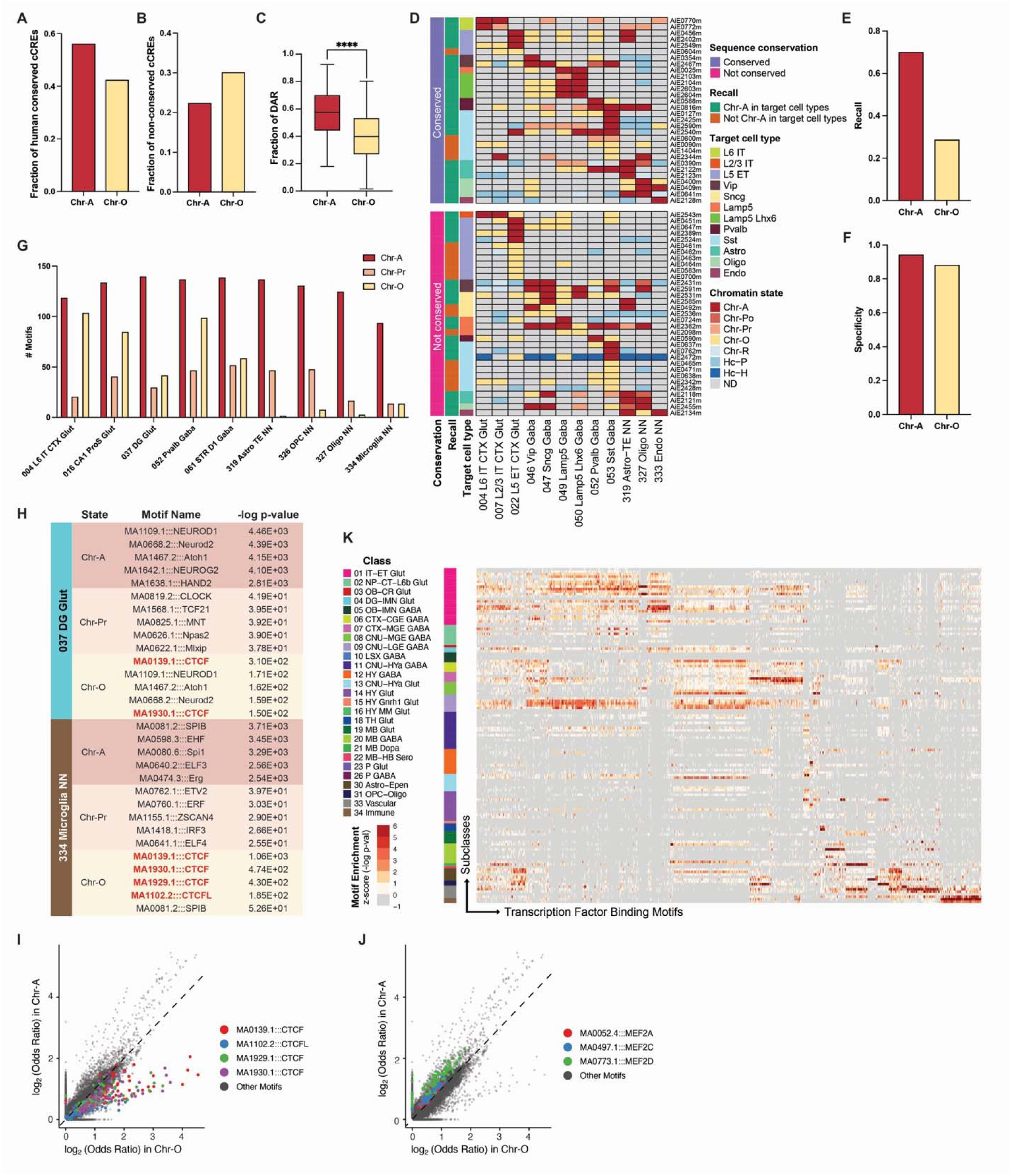
Functional and epigenomic comparison of Chr-A and Chr-O, related to Figure 4. A. Bar plot showing the fraction of human conserved cCREs in Chr-A and Chr-O. B. Bar plot showing the fraction of non-conserved cCREs in Chr-A and Chr-O. C. Box plot showing the fraction of differentially accessible regions (DAR) assigned as Chr-As and Chr-Os across different brain cell subclasses. Statistical significance was assessed using the Wilcoxon signed-rank test. ****, p-value < 0.0001. D. Heatmap showing the dominant chromatin state annotation of validated enhancers from Allen Institute Enhancer (AiE) collection, with verified on-target cell-type-specific activity, across various relevant brain cell subclasses. These enhancers are stratified based on sequence conservation (presence of conserved sequence in the human genome), recall (whether Chr-A chromatin state annotation is detected in the validated target cell type), and target cell type (experimentally validated using AAV-enhancer reporter). E. Bar plots showing the recall of Chr-A and Chr-O annotations in predicting validated cell type-specific enhancers from the Allen Institute Enhancer (AiE) AAV reporter collection. Recall is defined as the fraction of AAV reporter–validated enhancers that are annotated as Chr-A or Chr-O in the corresponding cell types. F. Bar plots showing the specificity of Chr-A and Chr-O annotations in predicting validated cell type-specific enhancers from the Allen Institute Enhancer (AiE) AAV reporter collection. Specificity is defined as the fraction of cell types in which Allen AAV reporters tested negative and in which the corresponding genomic regions are not annotated as Chr-A or Chr-O. G. Number of non-redundant TF motifs that are significantly enriched in Chr-A, Chr-Pr and Chr-O across selected brain cell subclasses. Non-redundant TF motifs were defined as described previously^161^. H. Table summarizing the top five enriched TF motifs in Chr-A, Chr-Pr and Chr-O for brain cell subclasses 037 DG Glut and 334 Microglia NN. I. Scatter plot showing the log2 Odds Ratio of TF motif enrichment in Chr-A compared with Chr-O across subclasses, with CTCF and CTCFL motifs highlighted. J. Scatter plot showing the log2 Odds Ratio of TF motif enrichment in Chr-A compared with Chr-O across subclasses, with MEF2A, MEF2C and MEF2D motifs highlighted. K. Heatmap showing enrichment of transcription factor binding motifs across all brain cell subclasses.

**Figure S10.**
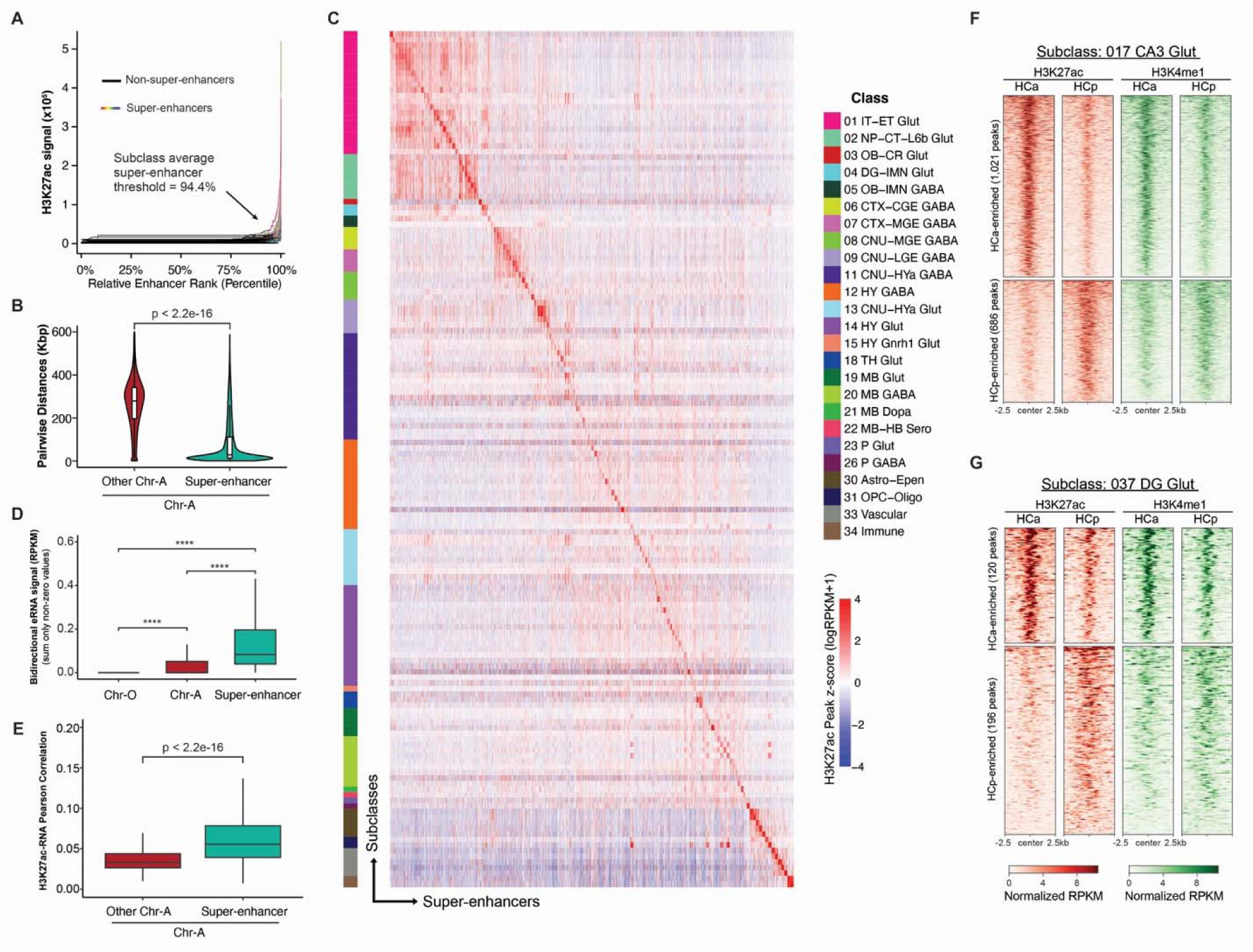
Identification and characterization of super-enhancers and brain-region specific epigenomic features, related to Figure 4. A. Rank Ordering of Super-Enhancers (ROSE) analysis of super-enhancers in brain cell subclasses. Subclass-level H3K27ac peaks were ranked by H3K27ac signal intensity, and super-enhancers were defined using the inflection point of the rank curve. The average super-enhancer threshold across subclasses is 94.4% (top 5.6%). B. Pairwise genomic distance between super-enhancer-overlapping Chr-A regions and other Chr-A regions. Statistical significance was assessed using the Wilcoxon rank sum test. C. Heatmap showing H3K27ac signal across identified super-enhancers across brain cell subclasses. Super-enhancers are grouped by the subclass in which they were identified, highlighting their subclass-specific activity patterns. D. Enhancer RNA signal associated with Chr-O regions, Chr-A regions, and super-enhancers. Super-enhancers showed the strongest enhancer RNA signal among the three regulatory element categories. Statistical significance was assessed using pairwise Wilcoxon rank-sum test with Benjamini-Hochbery correction. E. Box plot showing distribution of single-cell H3K27ac-RNA Pearson correlation of super-enhancers and non-super-enhancer Chr-A regions. Statistical significance was assessed using the Wilcoxon rank sum test. F. Coverage heatmaps of H3K27ac and H3K4me1 signals at differentially enriched H3K27ac peaks between anterior (HCa) and posterior (HCp) hippocampus in subclass 017 CA3 Glut neurons. G. Coverage heatmaps of H3K27ac and H3K4me1 signals at differentially enriched H3K27ac peaks between HCa and HCp in subclass 037 DG Glut neurons.

**Figure S11.**
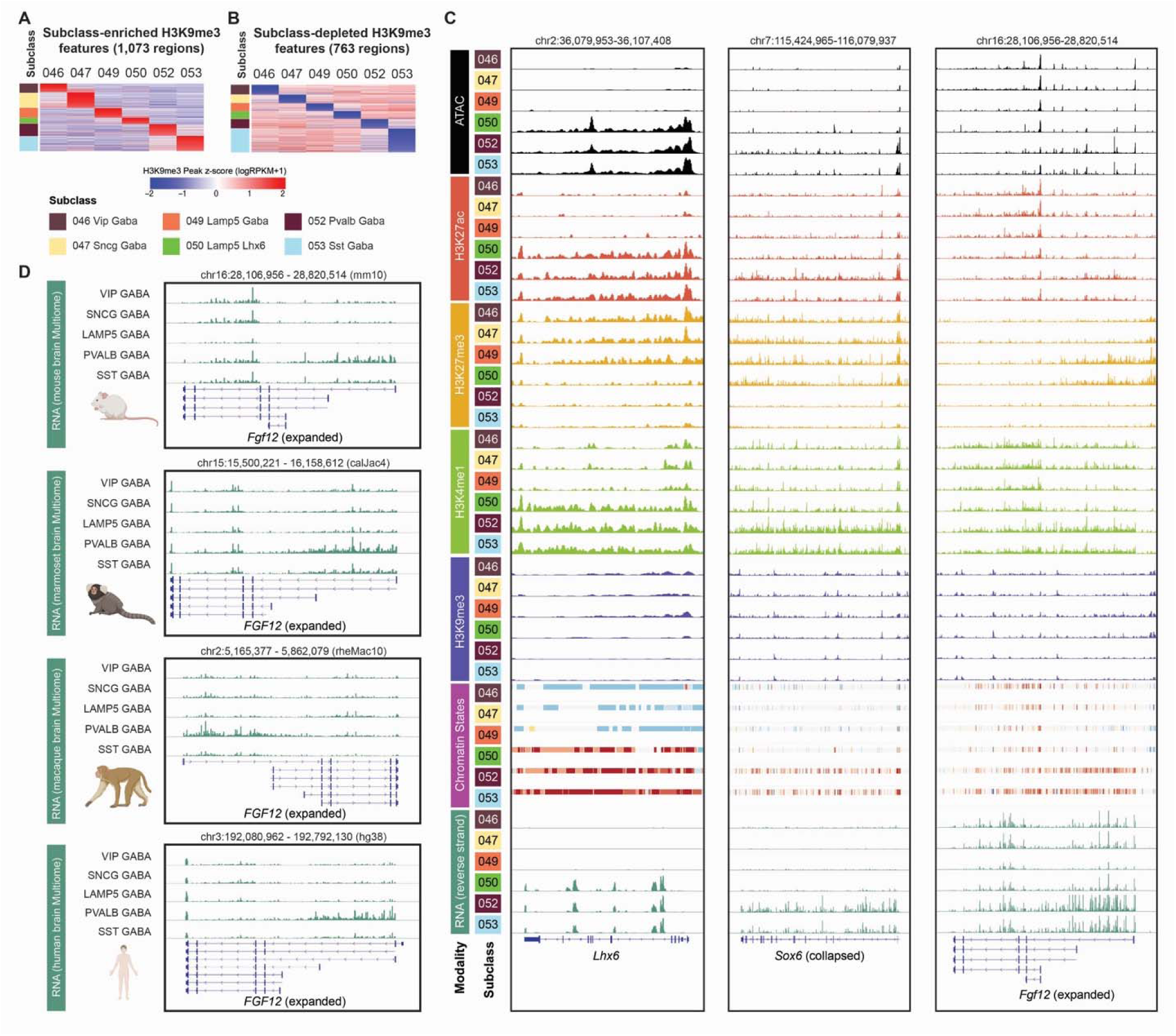
Cell-type-specific repressive features in the mouse brain, related to Figure 5. A. Heatmap showing normalized H3K9me3 signal over subclass-enriched H3K9me3 features in selected CGE- and MGE-derived GABAergic neuronal subclasses. B. Heatmap showing normalized H3K9me3 signal over subclass-depleted H3K9me3 features in selected CGE- and MGE-derived GABAergic neuronal subclasses. C. Genome browser tracks showing ATAC, H3K27ac, H3K27me3, H3K4me1, H3K9me3, chromatin states, and RNA signals, for selected CGE- and MGE-derived GABAergic neuronal subclasses, at *Lhx6*, *Sox6* and *Fgf12* genomic loci. D. Genome browser tracks showing RNA signals at *Fgf12* genomic loci for VIP, SNCG, LAMP5, PVALB and SST GABAergic neurons in mouse, marmoset, macaque, and human brains. Cell-type-resolved RNA-seq tracks of the brain from various species were generated using 10xGenomics Multiome from our previous work^37^.

**Figure S12.**
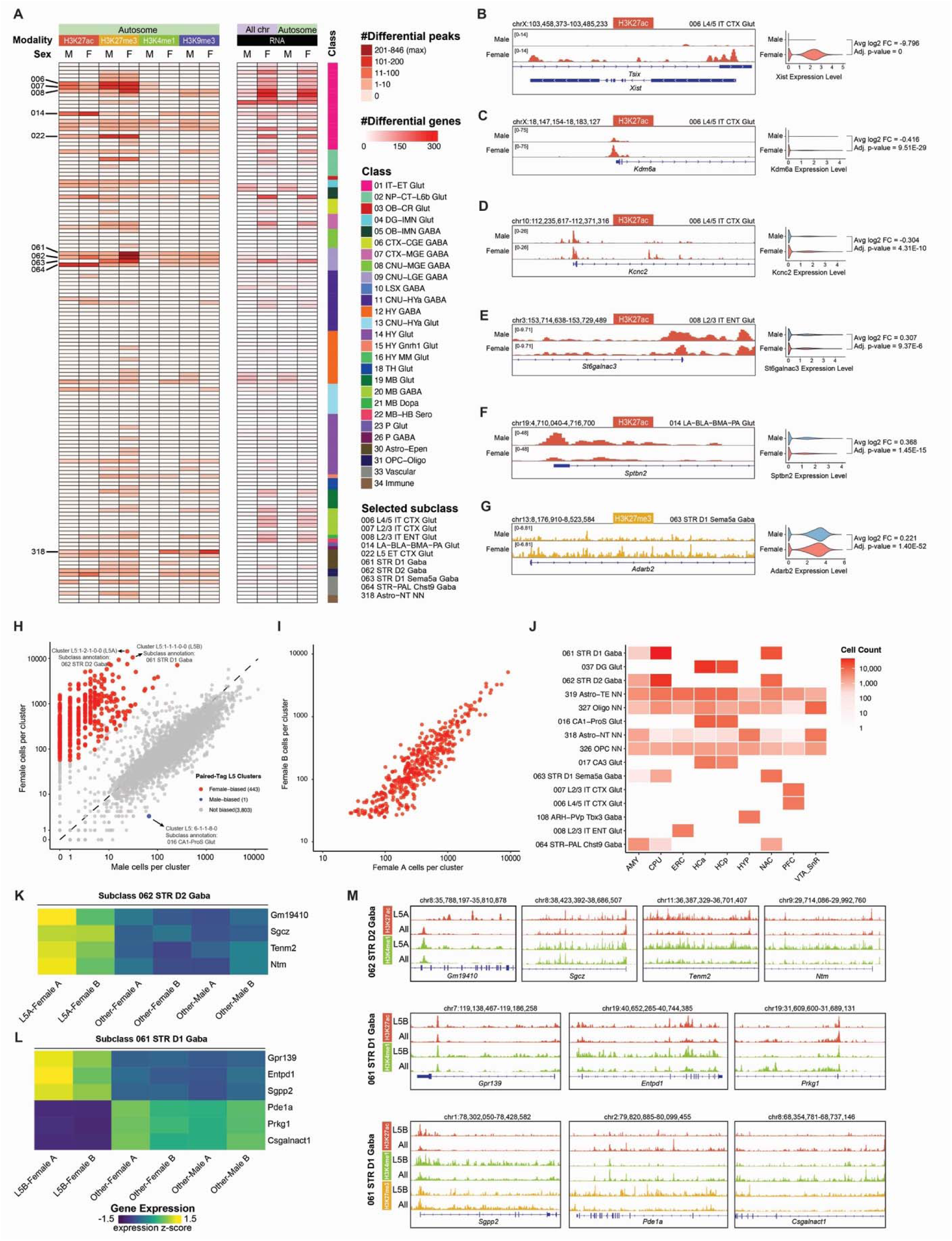
Sex-biased epigenomic features, related to Figure 6. A. Left, heatmap showing the number of differentially marked H3K27ac, H3K27me3, H3K4me1 and H3K9me3 peaks on autosomes across brain cell subclasses; Right, heatmap showing the number of sex-biased genes across brain cell subclasses, stratified by male-preferential and female-preferential genes on all chromosomes and autosomes only. B-D. Left, genome browser tracks showing sex-differential H3K27ac signal at the *Xist* (B), *Kdm6a* (C), and *Kcnc2* (D) gene loci in 006 L4/5 IT CTX Glut subclass. Tracks are shown for male and female samples. Right, violin plots showing sex-differential expression of *Xist* (B), *Kdm6a* (C), and *Kcnc2* (D) in male and female cells of the same subclass. Statistical significance was assessed using Wilcoxon rank-sum test followed by Bonferroni correction. E. Left, genome browser tracks showing sex-differential H3K27ac signal at the *St6galnac3* gene locus in 008 L2/3 IT ENT Glut subclass. Tracks are shown for male and female samples. Right, violin plot showing sex-differential expression of *St6galnac3* in male and female cells of the same subclass. Statistical significance was assessed using Wilcoxon rank-sum test followed by Bonferroni correction. F. Left, genome browser tracks showing sex-differential H3K27ac signal at the *Sptbn2* gene locus in 014 LA-BLA-BMA-PA Glut subclass. Tracks are shown for male and female samples. Right, violin plot showing sex-differential expression of *Sptbn2* in male and female cells of the same subclass. Statistical significance was assessed using Wilcoxon rank-sum test followed by Bonferroni correction. G. Left, genome browser tracks showing sex-differential H3K27ac signal at the *Sptbn2* gene locus in 014 LA-BLA-BMA-PA Glut subclass. Tracks are shown for male and female samples. Right, violin plot showing sex-differential expression of Sptbn2 in male and female cells of the same subclass. Statistical significance was assessed using Wilcoxon rank-sum test followed by Bonferroni correction. H. Scatter plot showing the number of male and female cells in each Paired-Tag L5 cluster. Significant female-biased and male-biased clusters are highlighted in red and blue, respectively. Dashed line represents the overall sex ratio of the dataset. I. Scatter plot showing the number of cells from each female biological replicate in each female-biased Paired-Tag L5 cluster. J. Heatmap showing the distribution of female-biased L5 clusters across subclasses and brain regions. Color intensity represents the cell count for each subclass-brain region combination. K, L. Heatmaps showing selected genes that are differentially expressed in L5A cluster (L5 cluster 1-2-1-0-0, from subclass 062 STR D2 Gaba) (K) and L5B cluster (L5 cluster 1-1-1-0-0, from subclass 061 STR D1 Gaba) (L), across biological replicates. M. Genome browser tracks showing histone modification Paired-Tag data tracks on genomic loci encoding L5A (top) and L5B (middle and bottom) differentially expressed genes, comparing female-enriched L5 clusters with the entire subclass.

**Figure S13.**
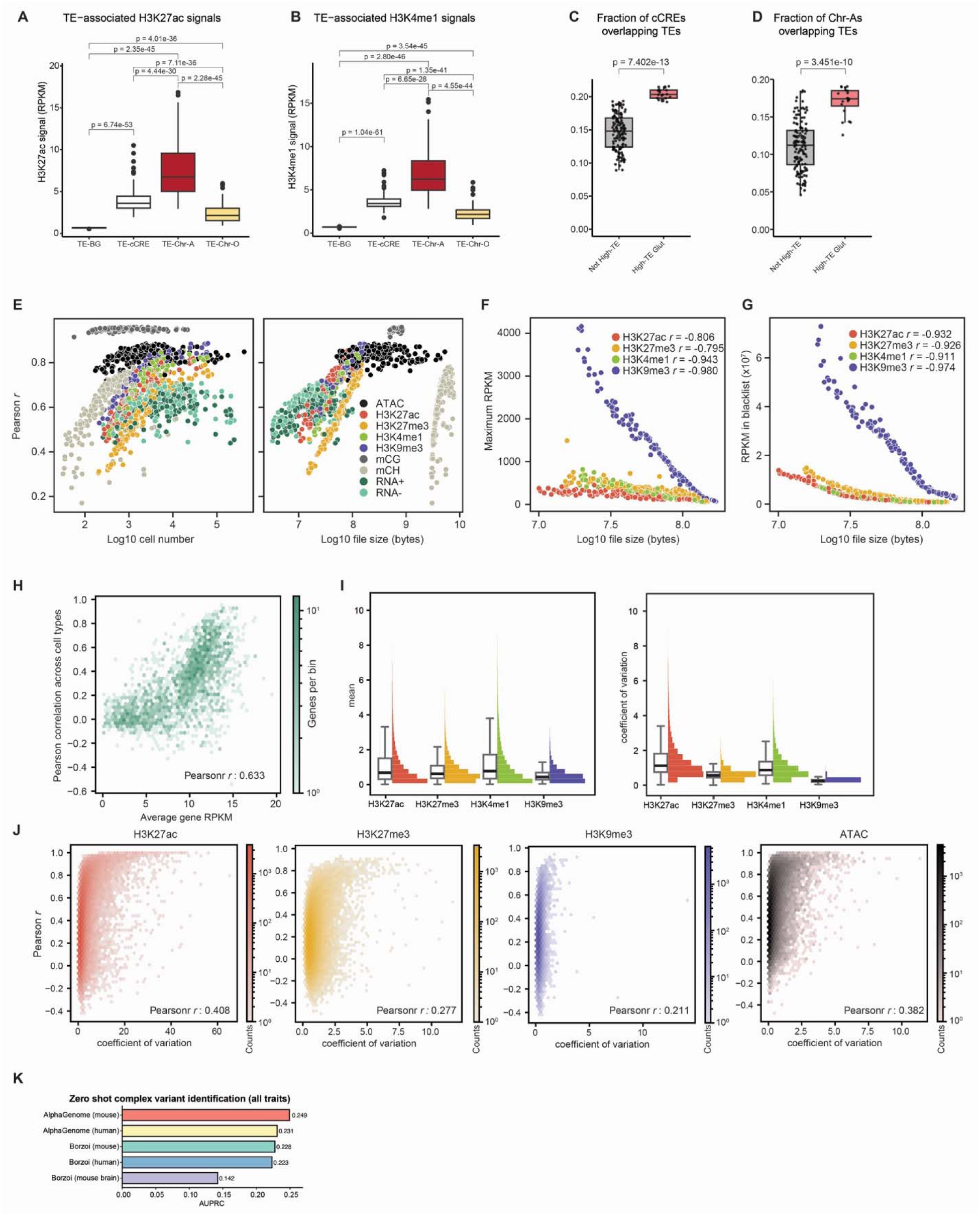
Transposable elements with regulatory signatures in the mouse brain, and mouse brain Borzoi model, related to Figures 6 and 7. A. Box plot showing H3K27ac signals on TEs stratified by genomic context: TEs not overlapping brain cCRE (TE-BG), TEs overlapping brain cCREs (TE-cCRE), TEs overlapping Chr-As (TE-Chr-A), and TEs overlapping Chr-Os (TE-Chr-O). Statistical significance was assessed using two-sided *t* test with Benjamini-Hochberg correction. B. Box plot showing H3K4me1 signals on TEs stratified by genomic context: TEs not overlapping brain cCRE (TE-BG), TEs overlapping brain cCREs (TE-cCRE), TEs overlapping Chr-As (TE-Chr-A), and TEs overlapping Chr-Os (TE-Chr-O). Statistical significance was assessed using two-sided *t* test with Benjamini-Hochberg correction. C. Box plot showing the fraction of cCREs overlapping TEs in High-TE Glut subclasses compared with other, not High-TE subclasses. Statistical significance was accessed using Wilcoxon rank-sum test. D. Box plot showing the fraction of Chr-As overlapping TEs in High-TE Glut subclasses compared with other, not High-TE subclasses. Statistical significance was accessed using Wilcoxon rank-sum test. E. Left, scatter plot showing the log10 number of cells in a cell type compared to the Pearson correlation between measurements and predictions in the test set. Right, scatter plot showing the log10 file size and the test set Pearson correlation for model predictions. F. Scatter plot showing the log10 file size in bytes and the maximum RPKM value in a track for each histone modification track used in model training. G. Scatter plot showing the log10 file size and the total RPKM values mapped to blacklist regions for each histone modification track used in training. H. Hexbin plots showing the correlation between model predictions and measurements across subclasses and the coefficient of variation for each peak region in the test set. From left to right, plots correspond to H3K27ac, H3K27me3, H3K9me3, and ATAC. I. Hexbin plot showing the Pearson correlation between predicted and measured average gene-level RPKM values for each gene in the test set. J. Left, boxplots and histograms showing the distribution of mean RPKM across subclasses for each peak in the test set, stratified by modality. Right, boxplots and histograms showing the coefficients of variation for each peak in the test set. K. Zero-shot performance of the mouse brain fine-tuned Borzoi model developed in this study on all traits in the TraitGym benchmark, compared with AlphaGenome mouse and human models and untuned Borzoi mouse and human models.

## RESOURCE AVAILABILITY

### Lead contact

Requests for further information and resources should be directed to and will be fulfilled by the lead contact, Bing Ren.

### Materials availability

This study did not generate new unique reagents.

### Data and code availability

All software, sequencing data from this study, and the sources of publicly available data used in this study are listed in the key resources table. Raw data have been deposited to the NeMO Archive (RRID: SCR_016152) with the following collection IDs (nemo:col-wgpj7cd; nemo:col-xt1ksye, landing page: https://assets.nemoarchive.org/grant/nemo:grn-f309ksd). Processed data are available through CATlas web portal (https://catlas.org/catlas/amb-pt, RRID: SCR_018690) and Gene Expression Omnibus (https://www.ncbi.nlm.nih.gov/geo/, RRID: SCR_005012) under the accession number: GSE317574, and will be made publicly available upon acceptance of the manuscript. Custom codes and scripts used for analysis are available at GitHub (https://github.com/beyondpie/amb_pairedtag).

## ACKNOWLEDGEMENTS

We thank Dr. Chris Jeans and the QB3 Macrolab at UC Berkeley for the purification of recombinant proteins used in this study, Dr. Kristen Jepsen and the IGM Genomics Center at UC San Diego for the Illumina Sequencing, Dr. Kai Zhang for guidance on bioinformatic analysis, and the San Diego Supercomputer Center for support with the Triton Shared Computing Cluster (TSCC) high-performance computing resources. We thank Drs. Yanxiao Zhang, Jie Xu, and Guojie Zhong, as well as all members of the Ren, Behrens, and Ecker laboratories, for scientific input, feedback, and discussions. Some schematics were created with BioRender. This work was supported by the National Institute of Mental Health (1RF1MH128838-01 to B.R. and M.M.B.). This work includes data generated at the UC San Diego IGM Genomics supported by a National Institutes of Health SIG grant (S10 OD026929). Z.W. was a DDBrown Awardee of the Life Sciences Research Foundation and was supported by the National Institute of Health through a Pathway to Independence Award (1K99HL183669-01).

## AUTHOR CONTRIBUTIONS

B.R., M.M.B., Z.W. and C.Z. conceived and designed the study. B.R. and M.M.B. supervised the study. Z.W., T.H.L., J.A.R, N.D.J., J.L., J.K.W., S. Cho, S. Cao, A.S.B, N.E. and M.M.B. performed the experiments. Z.W., Y.X., L.C. and C.Z. developed the new Paired-Tag experimental protocol and analytical pipelines. Z.W., S.Z., E.J.A., Y.X., K.W., B.S., Z.A.G., X.G., S.X. and H.L. analyzed the data. E.J.A. developed the deep learning models and the SeqNN web portal. Z.W., S.Z., D.G. and Z.T. participated and contributed to the development of deep learning models. W.W. and Y.L. developed the CATlas web portal. B.R., M.M.B., J.R.E., Z.W. and J.A.R coordinated the research. Z.W., S.Z., E.J.A., M.M.B. and B.R. wrote and edited the manuscript, with input from all authors. All authors have read, edited, and approved the final version of the manuscript.

## DECLARATION OF GENERATIVE AI AND AI-ASSISTED TECHNOLOGIES IN THE WRITING PROCESS

During the preparation of this work, the authors used OpenAI’s ChatGPT to improve the language of the manuscript. After using this tool, the authors reviewed and edited the content as needed and take full responsibility for the content of the published article.

## Key Resources Table

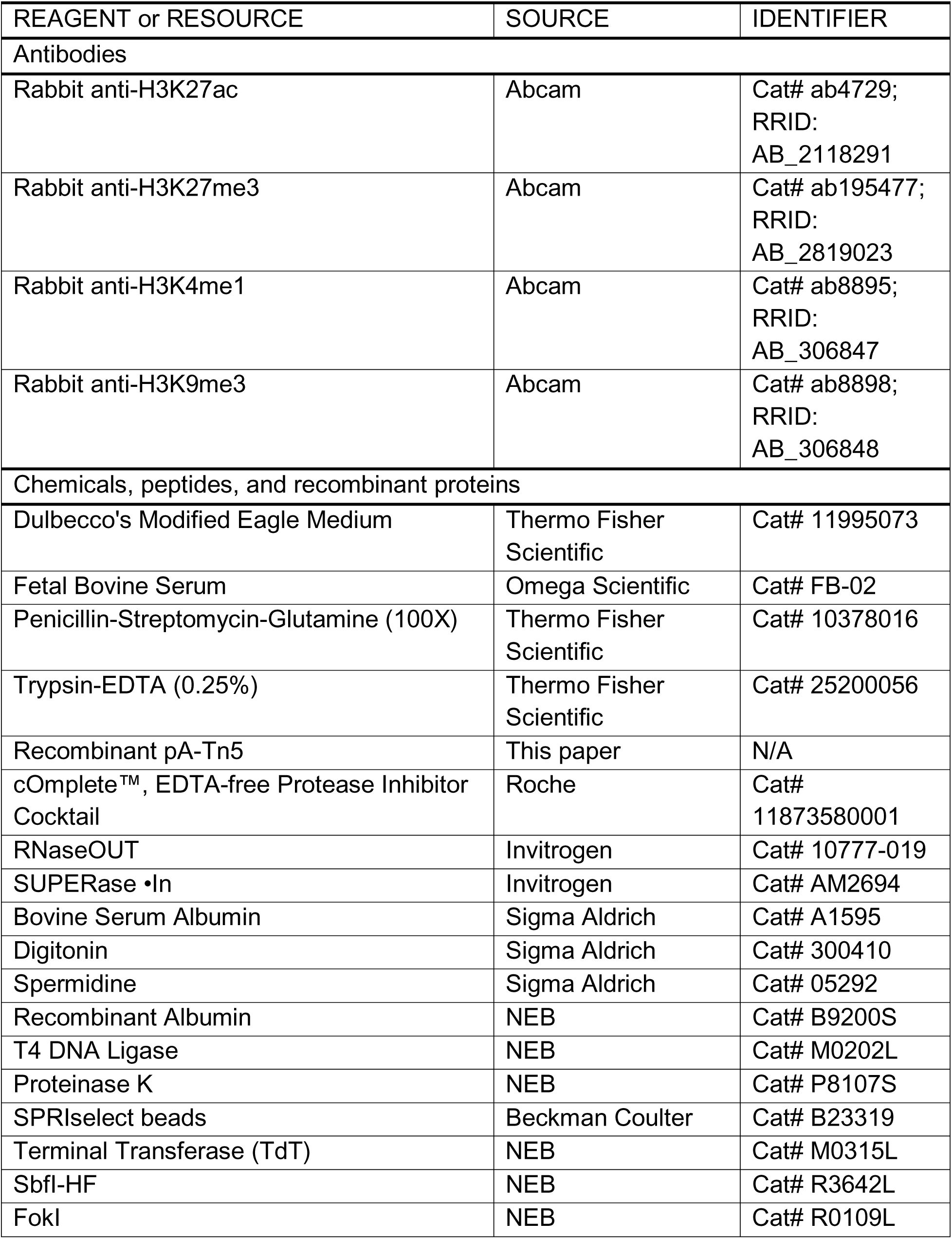
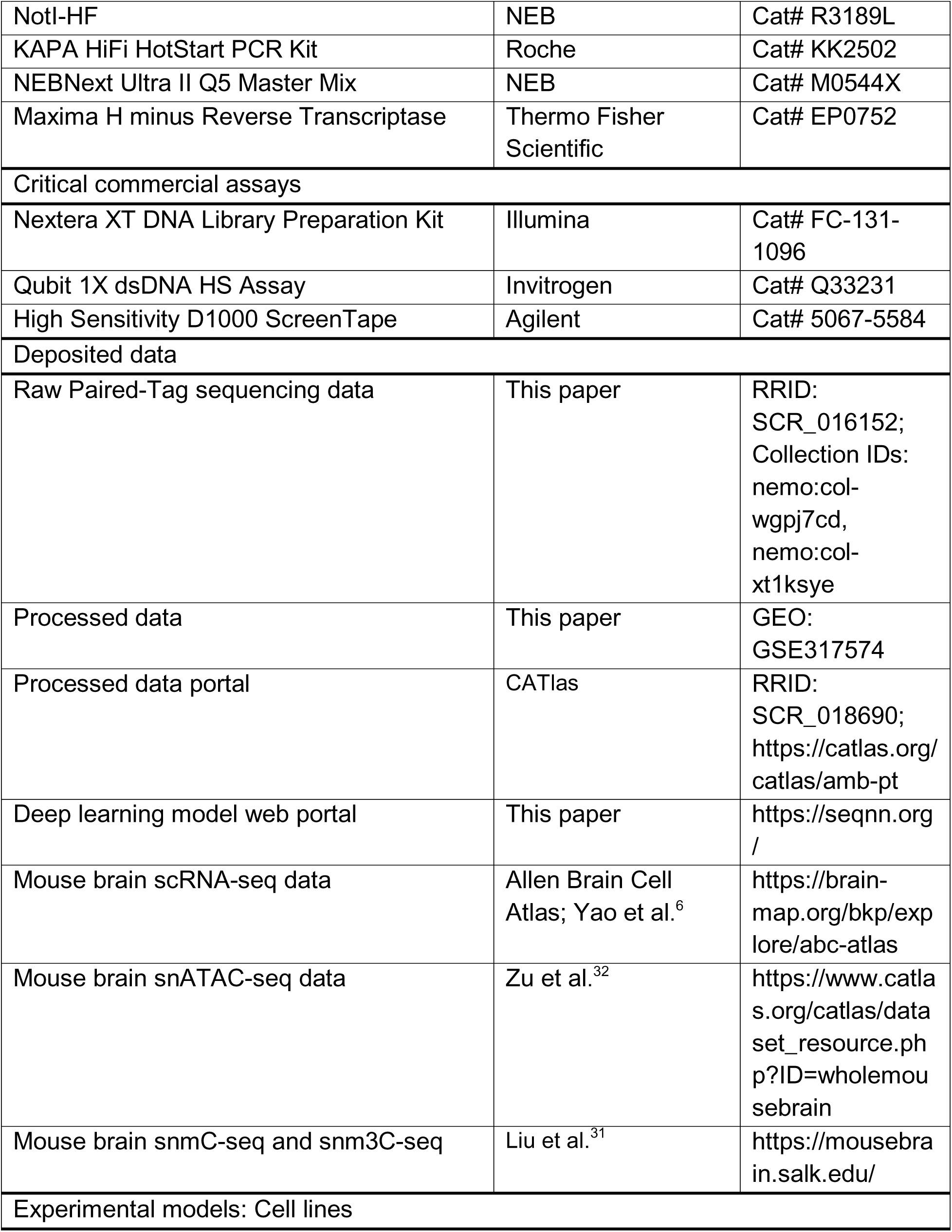
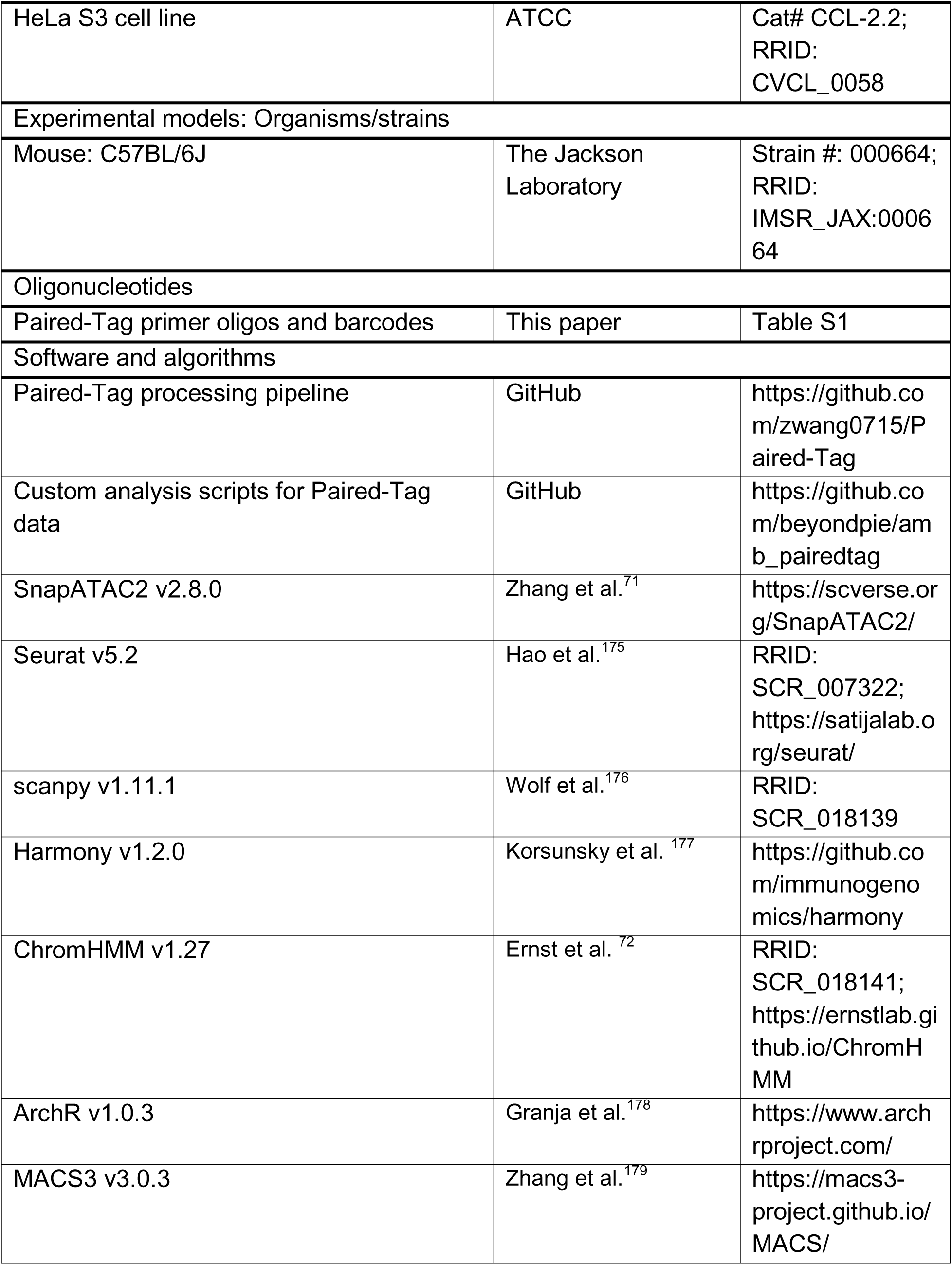
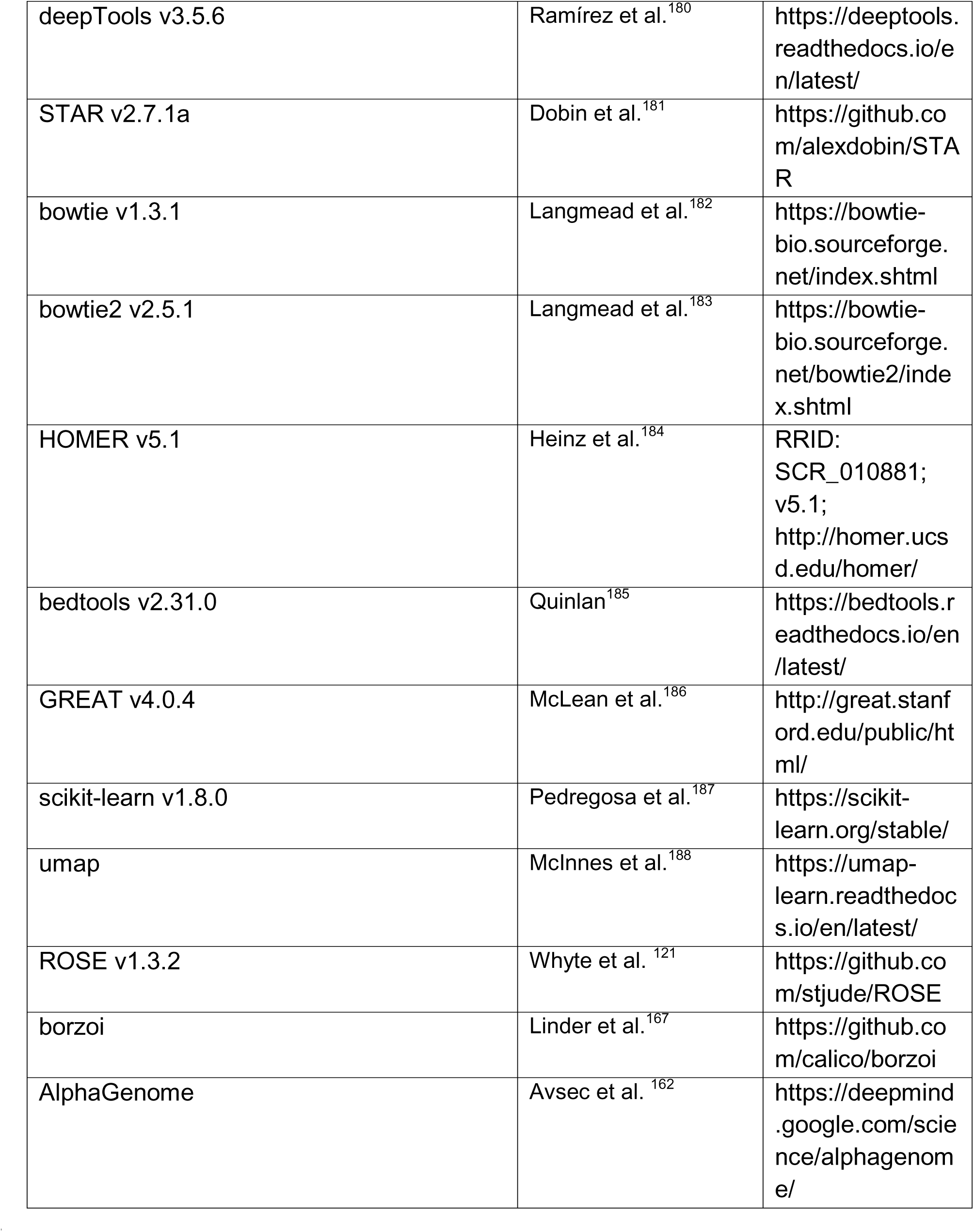

## Experimental model and subjects details

### Experimental animals

Animal work described in this manuscript has been approved and conducted in accordance with the Institutional Animal Care and Use Committee (IACUC) protocols at the Salk Institute. C57BL/6J mice (000664, RRID: IMSR_JAX:000664) were purchased from the Jackson Laboratories at 6 weeks of age and maintained in the Salk animal facility on a 12-h dark/light cycle with controlled temperature (20-22 C) and humidity (30-70%), and with *ad libitum* access to food and water.

### Cell lines and cell culture

HeLa S3 (ATCC, CCL-2.2) cells were cultured in Dulbecco’s Modified Eagle Medium (ThermoFisher Scientific, 11995073) supplemented with 10% fetal bovine serum (Omega Scientific, FB-02) and 1% penicillin-streptomycin-glutamine (ThermoFisher Scientific, 10378016), at 37 C with 5% CO2. Cells were not authenticated or tested for mycoplasma.

## Method details

### Mouse tissue preparation and nuclei isolation

Brains of C57BL/6J mice were collected from 8-week-old mice and sectioned into 600 μm coronal sections along the anterior-posterior axis in ice-cold dissection medium [20 mM Sucrose, 28 mM D-Glucose (Dextrose), 0.42 mM NaHCO3, in HBSS]^35,163^. Specific brain regions were dissected according to the Allen Brain Reference Atlas^71^, and stored at -80°C, as previously described^34,35^. For nuclei isolation of each brain region, dissected brain tissues were pooled from 2-8 animals of the same sex to obtain enough nuclei for Paired-Tag experiment for each biological replica, and two biological replicas were performed for both male and female samples. Frozen brain tissue samples were first dounce-homogenized with pestle A x 5-10 times and pestle B x 15-20 times in ice-cold Douncing Buffer [0.25 M sucrose (Sigma, S7903), 25 mM KCl (Invitrogen, AM9640G), 5 mM MgCl2 (Invitrogen, AM9530G), 10 mM Tris-HCl pH 8.0 (Invitrogen, 15568025), 1 mM DTT (Sigma, 43816), 1× cOmplete Protease Inhibitor cocktail (Roche, 11873580001), 0.5 U μL^−1^ RNase OUT (Invitrogen, 10777-019) and 0.5 U μL^−1^ SUPERase Inhibitor (Invitrogen, AM2694)] supplemented with 0.1% Triton X-100 (Sigma, 93443). Homogenates were then filtered through 30-µm Cell-Trics filters (Sysmex, 04-004-2326) and spun down for 10 min at 1,000 x g and 4°C. The nuclei pellets were resuspended with an equal volume of ice-cold Douncing Buffer and spun down again. The washed nuclei pellets were permeabilized in ice-cold Nuclei Isolation Buffer (NIB) [5% bovine serum albumin (Sigma, A1595) in PBS (ThermoFisher Scientific, 14190250), 0.2% IGEPAL CA-630 (Sigma, I8896), 1× cOmplete Protease Inhibitor cocktail, 1 mM DTT, 0.5 U μL^−1^ RNase OUT, and 0.5 U μL^−1^ SUPERase Inhibitor] for 5 min. The nuclei were counted on a cell counter (RWD, C100) upon DAPI staining and subjected to Paired-Tag experiments immediately.

### Spike-in HeLa cell preparation

HeLa S3 cells from passage 1 to 25 were used as spike-in control cells in the Paired-Tag experiment. HeLa cells were dissociated with 0.25% Trypsin-EDTA (ThermoFisher Scientific, 25200056), washed with PBS, and incubated in ice-cold NIB for 5 min. The HeLa nuclei were counted on a cell counter upon DAPI staining and subjected to Paired-Tag experiments immediately.

### Paired-Tag experimental procedures

Paired-Tag experiments were performed as previously described^62^, with major modifications in barcode design to improve the throughput, as detailed below.

#### Oligo sequences and assembly of transposomes

All oligo sequences used can be found in Supplementary Table 1. Specifically, 48 newly designed barcoded DNA adaptor oligos were reconstituted with nuclease-free H2O (ThermoFisher Scientific, AM9930), mixed with a pMENTs oligo individually (final concentration of 50 μM) and annealed with the following program using a thermocycler: 95°C x 5 min, then slowly ramp down to 4°C at a speed of -0.1°C s^−1^. Then, annealed adaptor oligos were mixed with unloaded Protein A-Tn5 (0.7 mg mL^−1^) in a 1:6 volume ratio. The mixtures were briefly vortexed and spun down with a desktop centrifuge, incubated at room temperature for 30 min then at 4°C for an additional 10 min. The assembled transposome complexes (also referred to as barcoded pA-Tn5) were stored at -20°C for up to 6 months. For barcoded reverse transcription (RT) primers, 48 newly designed barcoded poly-dT RT primers and 48 newly designed barcoded random hexamer RT primers were reconstituted at 100 μM, then mixed and diluted with nuclease-free water correspondingly to make 48 barcoded RT primer mixes, with both poly-dT and random hexamer RT primers at a final concentration of 12.5 μM.

#### Antibody-pA-Tn5 incubation and targeted tagmentation

For each Paired-Tag experiment, usually 16-32 reactions were included, where each reaction profiled one sample and one histone modification. The following histone modification antibodies were used: anti-H3K27ac (Abcam, ab4729), anti-H3K27me3 (Abcam, ab195477), anti-H3K4me1 (Abcam, ab8895), anti-H3K9me3 (Abcam, ab8898).

For each Paired-Tag reaction, 2 μg antibody and 1 μL barcoded pA-Tn5 were diluted in 30 μL MED1 buffer [1% BSA, 300 mM NaCl, 20 mM HEPES pH 7.5 (Invitrogen, 15630106), 1× cOmplete Protease Inhibitor cocktail, 0.01% Digitonin (Sigma, 300410), 0.5 mM Spermidine (Sigma, 05292), 2 mM EDTA (ThermoFisher Scientific, 15575020), 0.01% IGEPAL CA-630, 1 mM DTT, 0.5 U μL^−1^ RNase OUT, and 0.5 U μL^−1^ SUPERase Inhibitor] and pre-incubated on a tube rotator for 1 h at room temperature. 500-800k nuclei from each sample were resuspended in 45 μL MED1 buffer, mixed with pre-incubated antibody-pA-Tn5-MED1 mix, and further incubated overnight on a tube rotator at 4°C. Within each Paired-Tag experiment, we usually spike in ∼10% HeLa nuclei (50-80 k) to mouse brain nuclei in one or two reaction tubes to estimate the barcode collision rate of the entire experiment. After this incubation, the nuclei were spun down at 500 x g at 4°C for 10 min, washed twice with 75 μL MED2 buffer (2% BSA, 300 mM NaCl, 20 mM HEPES pH 7.5, 1× cOmplete Protease Inhibitor cocktail, 0.01% Digitonin, 0.5 mM Spermidine, 0.01% IGEPAL CA-630, 1 mM DTT, 0.5 U μL^−1^ RNase OUT, and 0.5 U μL^−1^ SUPERase Inhibitor), and resuspended in 75 μL MED2 buffer. The tagmentation reaction was activated by adding 3 μL 250 mM MgCl_2_ and was carried out at 550 rpm, 37°C for 1 h in a ThermoMixer (Eppendorf). Following tagmentation, 24.8 μL of 40.4 mM EDTA was added to quench the reaction. Nuclei were then spun down at 500 x g at 4°C for 10 min and proceeded to reverse transcription immediately.

#### Reverse transcription

For each Paired-Tag reaction, nuclei were resuspended in 24 μL RT mix [6 μL 5x RT buffer, 6.9375 μL nuclease-free H_2_O, 6 μL PBS, 1.5 μL 10 mM dNTPs (NEB, N0447S), 0.375 μL SUPERase Inhibitor, 0.1875 μL RNase OUT, 3 μL Maxima H minus reverse transcriptase (ThermoFisher Scientific, EP0752)], and 6 μL of corresponding barcoded RT primer mix was added to each reaction with gentle pipetting to mix. RT reaction was performed in a thermocycler using the following program: Step 1, 50°C × 10 min; Step 2, 8°C x 12 s, 15°C x 45 s, 20°C x 45 s, 30°C x 45 s, 42°C x 2 min, 50°C x 5 min, repeat for 3 total cycles; Step 3, 50°C × 10 min; Step 4, hold at 12°C. After RT reaction, the nuclei were pooled to a pre-chilled 1.5 mL Maxymum recovery tube (Axygen, MCT-150-L-C), and 5% Triton X-100 were added to the nuclei pool to a final concentration of 0.1%. Nuclei were then spun down at 500 x g at 4°C for 10 min and proceeded to combinatorial barcoding immediately.

#### Ligation-based combinatorial barcoding

We designed new barcode sequences to expand the barcoding space of the Paired-Tag combinatorial indexing. 384 R02 barcodes and 384 R03 barcodes were designed and synthesized (Table S1). Combinatorial barcodes were prepared as previously described^62^, and 5 μL annealed combinatorial barcode was distributed to each well of a DNA lobind 96-well plate (Eppendorf, 0030129504) using a Biomek i7 automated liquid handler (Beckman Coulter). Combinatorial barcoding oligo sequences can be found in Supplemental Table 1. In total, 4 x R02 barcoding plates (384 different R02 barcodes) and 4 x R03 barcoding plates (384 different R03 barcodes) were prepared and stored at -20°C. R02 and R03 barcoding plates were thawed and equilibrated to room temperature before use.

Post-RT nuclei were resuspended in 2 mL of 1x NEBuffer 3.1 (NEB, B6003S) and then transferred to the ligation mix [4500 μL H_2_O, 1000 μL 10x T4 Ligase Buffer (NEB, B0202S), 200 μL 10x NEBuffer 3.1, 100 μL Recombinant Albumin (NEB, B9200S), T4 DNA Ligase (NEB, M0202L)]. Each 20 μL of ligation mix with nuclei was distributed to R02 barcoding plates using a multichannel pipette and incubated at 300 rpm for 30 min at 37°C in a ThermoMixer. Next, 5 μL of R02 blocking solution (prepared with 528 μL 100 μM R02 Blocking oligo, 500 μL 10x T4 Ligase Buffer, 972 μL H_2_O) was dispensed to each well using a multichannel pipette and the reaction was further incubated at 300 rpm for 30 min at 37°C in a ThermoMixer. The nuclei were then pooled and spun down at 500 x g at 4°C for 10 min. The next round of ligation-based combinatorial barcoding was performed using R03 barcoding plates similarly as the first round, except that after 30 min of the ligation reaction, 5 μL of R03 terminating solution (prepared with 528 μL 100 μM R03 Termination oligo, 1000 μL 500mM EDTA pH 8.0, 472 μL H_2_O) was added to each well to quench the ligation reaction. The nuclei were then collected by centrifugation at 500 x g, 4°C for 10 min. Post-barcoding nuclei were washed once in 1 mL PBS, resuspended in 1 mL PBS, and counted. Nuclei were aliquoted to sublibraries each containing 2,500 to 5,000 nuclei diluted to 35 μL in PBS. For each sublibrary, 5 μL 4M NaCl, 5 μL 10% SDS (Corning, 46-040-CI), and 5 μL Proteinase K (NEB, P8107S) were added, and the nuclei were lysed at 850 rpm, 55°C for 2 h to overnight in a ThermoMixer. The PK-digested sublibraries were cooled to room temperature, purified with 1x SPRIselect beads (Beckman Coulter, B23319), and eluted in 25 μL of H_2_O. The purified sublibraries were stored at -20°C, or proceeded for library preparation immediately.

#### Preamplification of barcoded DNA/cDNA

For each 25 μL purified sublibrary from the previous step, 3 μL 10x TdT Buffer and 1 μL 1 mM dCTP (NEB, N0446S) were added and the reaction was incubated at 95°C for 5 min, and then immediately chilled on ice for 5 min. Next, 1 μL Terminal Transferase (NEB, M0315L) was added, and the reaction mixture was incubated at 37°C for 30 min followed by 75°C for 20 min in a thermocycler. Subsequently, 30 μL Anchor Mix [14.4 μL H_2_O, 12 μL 5x KAPA HiFi Fidelity Buffer, 1.2 μL KAPA dNTPs Mix, 1.2 μL 10 μM Anchor-FokI-GH oligo, and 1.2 μL KAPA HiFi HotStart DNA Polymerase (Roche, KK2502)] was added to each reaction and the following thermocycling steps were performed in a thermocycler: Step 1, 98°C x 3 min; Step 2, 98°C x 15 s, 47°C x 1 min, 68°C x 2 min, 47°C x 1 min, 68°C x 2 min, repeat for a total of 16 cycles; Step 3, 72°C × 10 min; Step 4, hold at 12°C. Immediately afterwards, 40 μL Preamplification Mix (22 μL H_2_O, 8 μL 5x KAPA HiFi Fidelity Buffer, 4 μL 10 μM PA-F oligo, 4 μL 10 μM PA-R oligo, 1 μL KAPA dNTPs Mix, 1 μL KAPA HiFi HotStart DNA Polymerase) was added to each reaction and the following thermocycling steps were performed: Step 1, 98°C x 3 min; Step 2, 98°C x 20 s, 62°C x 20 s, 72°C x 2 min, repeat for a total of 10 cycles; Step 3, 72°C x 2 min; Step 4, hold at 12°C. Preamplified sublibraries were purified with double size selection (0.2x + 0.65x, 20 μL + 65 μL beads) using SPRIselect beads, and eluted in 40 μL of H_2_O. From purified sublibraries, 1 μL of eluted DNA was used in a Qubit 1x dsDNA High Sensitivity Assay (Invitrogen, Q33231) to measure DNA concentration. For the remaining purified sublibraries, 4.5 μL CutSmart Buffer (NEB, B6004S) was added to each reaction, from which 19 μL of the reaction mixture was transferred to a new tube (hereafter referred to as the RNA sublibrary tube). The original tube holding the remaining reaction is referred to as the DNA sublibrary tube from this point onward.

#### Restriction enzyme digestion and purification of DNA and RNA sublibraries

For each DNA sublibrary tube, 1 μL SbfI-HF (NEB, R3642L) and 1 μL FokI (NEB, R0109L) were added. For each RNA sublibrary tube, 1 μL NotI-HF (NEB, R3189L) was added. The restriction enzyme digestion reactions were incubated at 37°C for 2 h to overnight, and further purified using 1.25x SPRIselect beads (25 μL beads into each RNA sublibrary tube, 31.3 μL beads into each DNA sublibrary tube). Both post-digestion DNA and RNA sublibraries were eluted in 10 μL H_2_O.

#### DNA sublibrary preparation

For each 10 μL purified, post-digestion DNA sublibrary, 10 μL P5 adaptor ligation mix (5 μL H_2_O, 2 μL 10x T4 Ligase Buffer, 1.5 μL 10 μM P5 Adaptor mix, 1.5 μL T4 DNA ligase) was added and the ligation reaction was carried out using the following thermocycling program: 4°C × 10 min, 10°C × 10 min, 16°C x 15 min, 25°C x 1h, hold at 4°C. P5 Adaptor mix was prepared by: (1) mixing 25 μL 100 μM P5-FokI oligo with 25 μL 100 μM P5c-NNDC-FokI oligo; (2) mixing 25 μL 100 μM P5A-FokI oligo with 25 μL 100 μM P5Ac-NNDC-FokI oligo; (3) mixing 25 μL 100 μM P5C-FokI oligo with 25 μL 100 μM P5Cc-NNDC-FokI oligo; (4) mixing 25 μL 100 μM P5T-FokI oligo with 25 μL 100 μM P5Tc-NNDC-FokI oligo; (5) annealing oligo mixtures from (1)-(4) with the following program using a thermocycler: 95°C x 5 min, then slowly ramping down to 4°C at a speed of -0.1°C s^−1^; (6) mixing four annealed oligo mixtures in a 1:1:1:1 ratio; then diluting from 50 μM to 10 μM final concentration.

The ligation product was then purified by 1.25x (25 μL) SPRIselect beads purification and eluted in 20 μL H_2_O. Next, 25 μL NEBNext Ultra II Q5 Master Mix (NEB, M0544X), 2.5 μL 10 μM TruSeq i7 indexing primer, and 2.5 μL 10 μM TruSeq i5 indexing primer were added to the eluted DNA sublibraries. The indexing PCR reaction was conducted with the following thermocycling program: Step 1, 98°C x 3 min; Step 2, 98°C × 10 s, 63°C x 30 s, 72°C x 1 min, repeat for a total of 12 cycles; Step 3, 72°C x 1 min; Step 4, hold at 12°C. The indexing PCR reaction was purified using 0.7x (35 μL) SPRIselect beads and eluted in 20 μL H_2_O. The size distribution of the final DNA sublibrary was examined using a High Sensitivity D1000 screentape (Agilent, 5067-5584) on a TapeStation 4200 device (Agilent), and library yield was examined using qPCR. Sequencing was performed with an Illumina NextSeq 2000, NovaSeq 6000 or NovaSeq X sequencer using the following setting: Read 1 – 100 cycles, Index 1 – 8 cycles, Index 2 – 8 cycles, Read 2 – 100 cycles.

#### RNA sublibrary preparation

From each 10 μL post-digestion RNA sublibraries, 5 μL was transferred to a new tube for downstream library preparation. On ice, 10 μL Tagment DNA buffer (TD) and 5 μL Amplicon Tagment Mix (ATM) from the Nextera XT DNA Library Preparation Kit (Illumina, FC-131-1096) were added to 5 μL RNA sublibrary with gentle pipetting to mix, and the reaction mix was incubated at 55°C for 5 min. Immediately after the tagmentation reaction, 5 μL Neutralize Tagment Buffer (NT) was added to each reaction, and the reaction was incubated at room temperature for 5 min. Next, 5 μL H_2_O, 15 μL Nextera PCR Mastermix (NPM), 2.5 μL 10 μM TruSeq i7 indexing primer, and 2.5 μL 10 μM Nextera i5 indexing primer were added to the reaction, and the indexing PCR reaction was conducted with the following thermocycling program: Step 1, 72°C x 3 min; Step 2, 95°C x 30 sec; Step 3, 95°C × 10 s, 55°C x 30 s, 72°C x 30 s, repeat for a total of 14 cycles; Step 4, 72°C x 5 min; Step 5, hold at 12°C. The indexing PCR reaction was purified using 0.7x (35 μL) SPRIselect beads and eluted in 20 μL H_2_O. The size distribution of the final RNA sublibrary was examined using a High Sensitivity D1000 screentape on a TapeStation 4200 device and library yield was examined using qPCR. Sequencing was performed with an Illumina NextSeq 2000, NovaSeq 6000 or NovaSeq X sequencer using the following setting: Read 1 – 100 cycles, Index 1 – 8 cycles, Index 2 – 8 cycles, Read 2 – 100 cycles.

### Processing and alignment of sequencing reads

Cellular barcodes from the sequencing reads were first extracted by matching the linker sequences adjacent to the cellular barcodes, which were then mapped to the cellular barcodes reference using bowtie^164^. Reads with more than 1 nucleotide mismatch were discarded. The adapter sequences were trimmed from 3′ of DNA and RNA libraries, with Poly-dT and random hexamer sequences further trimmed from 3′ of RNA libraries. Cleaned reads were mapped to the mouse GRCm38/mm10 reference genome with STAR^165^ for RNA or bowtie2^166^ for DNA. Duplicated reads were removed based on the mapped position, cellular barcode, PCR index, and UMI. RNA alignment files were converted to a matrix with cells as columns and genes as rows. DNA alignment files were converted to a matrix with cells as columns and 5-kb genomic bins as rows. A detailed, step-by-step Paired-Tag data processing pipeline can be found at: https://github.com/zwang0715/Paired-Tag.

### Quality control and doublet removal

For each sublibrary, cellular barcodes with 500 - 30,000 DNA features and 200 - 30,000 RNA features were retained. The initial double rate in each sublibrary was estimated using the spike-in Hela S3 cells as described in ^167^. Species-specific barcodes were defined as barcodes with at least 75% of RNA reads mapped to the genome of the corresponding species. We next applied a modified version of Scrublet implemented in SnapATAC2^74^ to remove potential doublets in each sublibrary, using the estimated initial doublet rates described above. Briefly, more than 8,000 differentially expressed genes identified from a previous adult whole mouse brain single-cell RNA-seq study were used as features^6^. Spectral embedding was performed across barcodes from all sublibraries using *snapatac2.tl.spectral* with the parameters *n_comps* as 50 and *weighted_by_sd* as “TRUE”. Doublet scores were then predicted for each barcode within individual sublibraries using *snapatac2.preprocessing.scrublet*. Because the predicted doublet probabilities were generally small, consistent with the relatively low doublet rates in the Paired-Tag experiments, applying the default Scrublet thresholds resulted in few cells being flagged as doublets. Therefore, we manually removed barcodes with the highest predicted doublet scores such that the proportion of removed cells in each sublibrary matched the corresponding estimated doublet rate. In addition to the quality control steps described above, additional low-quality cells were removed during downstream clustering and integration, as described in the corresponding Methods sections.

### Iterative cell clustering

We implemented a five-round iterative clustering strategy using single-nucleus RNA expression profiles from Paired-Tag^6^. Briefly, all 2.5 million single nucleus RNA-seq profiles from Paired-Tag were used for the first round of clustering (L1 level) using a standard clustering workflow. In the second round (L2 level), independent clustering was performed for each of the 17 L1-level clusters. In the third round (L3 level), independent clustering was carried out for each of the 182 L2-level clusters. In the fourth round (L4 level), independent clustering was performed for 972 of the 1,041 L3-level clusters. In the fifth round (L5 level), independent clustering was performed for 400 of the 3,339 L4-level clusters. In total, this hierarchical procedure identified 4,302 cell clusters. Detailed procedures are described below.

#### Feature selection

For L1-level clustering, we used more than 8,000 differentially expressed genes defined in the previous adult mouse brain scRNA-seq study^6^. For subsequent clustering levels, highly variable genes were selected from this gene set using *<u>scanpy.pp</u>.highly_variable_genes* in *scanpy*^168^, with the parameter *flavor* as “seurat_v3”, which emulates the *FindVariableFeatures* function with the parameter *method* as “vst” in Seurat^169,170^. Because cell clusters became progressively smaller with increasing clustering depth, we selected the top 3,000 genes for L2-level clustering and the top 2,000 genes for L3-, L4-, and L5-level clusterings.

#### Dimensionality reduction

We applied *snapatac2.tl.spectral* from SnapATAC2 to project high-dimensional, sparse gene expression profiles into low-dimensional space. This approach performs spectral embedding of the normalized graph Laplacian defined by the cell-to-cell similarity matrix based on the cosine distance of highly variable genes’ expression. For L1-level clustering, the dimensionality of the embedding was set to 50, whereas a dimensionality of 30 was used for subsequent clustering levels. The parameter *weighted_by_sd* in the function was set to “TRUE”, as recommended.

#### Graph-based clustering

We constructed k-nearest neighbor graphs using *snapatac2.pp.knn* from SnapATAC2 with the parameter *method* as “kdtree”. For L1- and L2-level clusterings, the parameter *n_neighbors* was set to 50, whereas a value of 30 was used for subsequent clustering levels. Clustering was then performed using *snapatac2.tl.leiden* with *objective_function* as “modularity”. Because the clustering resolution strongly influences the number of identified clusters, we evaluated a range of resolution values from 0.1 to 2 in increments of 0.1. Clustering performance at each resolution was assessed using the Silhouette coefficient as implemented in scikit-learn^171^, together with visual inspection of two-dimensional UMAP projections. UMAP embeddings were computed from the spectral embeddings using the umap package^172^ with parameters a as 1.8956, b as 0.8005, and init as “spectral”. For each clustering on different levels, the final resolution was selected based on a combination of Silhouette scores and UMAP visualization.

#### Histone modification-based clustering

Single-cell histone modification profiles generated by Paired-Tag were processed using SnapATAC2 for dimensional reduction and clustering. Briefly, the genome was segmented into fixed-width bins, with a bin size of 5 kb for H3K27ac and H3K4me1, and 50 kb for H3K27me3 and H3K9me3, reflecting the distinct genomic distribution patterns of active and repressive histone marks. Blacklisted genomic regions were excluded, and only autosomal regions were retained for downstream analysis. For each histone modification, cell-by-bin count matrices were constructed and highly variable genomic bins were selected. Specifically, the top 250,000 most variable bins were retained for H3K27ac and H3K4me1, whereas the top 25,000 most variable bins were retained for H3K27me3 and H3K9me3. Dimensional reduction was then performed using spectral embedding as implemented in SnapATAC2. A k-nearest neighbor graph was constructed based on the spectral embedding, followed by Leiden clustering to identify cell clusters. For visualization, two-dimensional embeddings were generated using UMAP based on the same low-dimensional representation. To assess the concordance between histone mark-based embeddings and RNA-defined cell-type annotations, we computed normalized mutual information (NMI) and k-nearest neighbor (kNN) label transfer accuracy. NMI was calculated between Leiden cluster assignments and RNA-based cell-type labels using the *normalized_mutual_info_score function* in the *scikit-learn* package, with arithmetic mean normalization. kNN label transfer accuracy was computed by assigning each cell the majority RNA-based label of its k nearest neighbors in the SnapATAC2 spectral embedding space (k=15), excluding the cell itself. Only cells with valid labels in both modalities were included in the analysis.

#### Batch effect estimation and clustering quality control

To assess potential batch effects across biological replicates, we quantified replicate mixing within each major cell class using k-nearest neighbor Batch Effect Test (kBET) and Local Inverse Simpson’s Index (LISI). Cells were represented by their coordinates in the UMAP embedding generated from Paired-Tag data. For each cell class, cells from the four biological replicates (MaleA, MaleB, FemaleA, and FemaleB) were downsampled to equal numbers, and classes with fewer than 50 cells in any replicate were excluded. kBET was performed using a neighborhood size of 50 cells to compare local replicate composition against the expected global distribution, and LISI was calculated using replicate identity as the batch label. Higher kBET acceptance rates and higher LISI values indicate improved mixing of cells across biological replicates.

### Transfer label-based annotation with scRNA-seq data

We performed the transfer label-based annotation for more than 4,000 Paired-Tag-derived cell clusters using a published adult whole mouse brain scRNA-seq dataset as reference^6^. The nine brain regions profiled by Paired-Tag were mapped to their corresponding anatomical regions in the scRNA-seq dataset as follows: prefrontal cortex (PFC) to ACA and PL-ILA-ORB regions in the scRNA-seq study, hippocampus anterior and posterior (HCa and HCp) to HIP, entorhinal cortex (ERC) to ENT, nucleus accumbens (NAC) to STRv,caudate putamen (CPU) to STRd, amygdala (AMY) to STR-sAMY and CTXsp, ventral tegmental area-substantia nigra (VTR_SnR) to MB-VTA_SN, and Hypothalamus (HYP) to HY. Only cells from both the mapped regions and generated using the 10x Genomics single cell 3’ gene expression v3 platform were retained from the scRNA-seq dataset, resulting in 656,346 cells used for downstream analysis while preserving all relevant cell clusters. Transfer label analysis was performed separately for neuronal and non-neuronal cells. Neuronal and non-neuronal populations were distinguished based on *Snap25* expression in L1-level Paired-Tag clusters.

For non-neuronal cells, we randomly sampled 30 cells in each of more than 1,300 L5-level clusters in the Paired-Tag dataset and 1,000 cells in each of L4-level clusters in the scRNA-seq data (including non-neuronal and immature-neural cells) to keep the cells from each dataset roughly balanced. More than 8,000 genes previously defined from differential expression analysis in the scRNA-seq study were used as variable features. We applied the *FindTransferAnchors* function in Seurat using canonical component analysis (CCA) to co-embed the two datasets into a joint low-dimensional space, followed by *TransferLabel* to annotate Paired-Tag cell with the supertype-level label from the scRNA-seq data. For each L5-level cluster, the most frequently assigned supertype label was selected as the cluster annotation. Co-embedding results were visualized using Uniform Manifold Approximation and Projection (UMAP), computed from the joint embedding generated by *FindIntegrationAnchors* as recommended in Seurat. Annotations were further validated by manual inspection of canonical marker gene expression.

For neuronal cells, a similar transfer label procedure was applied but performed separately within each brain region, reflecting the strong regional specificity of neuronal cell types. Within each region, cells from the reference scRNA-seq dataset with region-mismatched annotations were first ignored, and L5-level clusters (about 0.69% cells) from Paired-Tag dataset with small number of cells were flagged and excluded from our later analysis. To retain representative populations while balancing cell numbers, we down-sampled L5-level clusters from both Paired-Tag and reference scRNA-seq datasets, including immature neuronal cells. Variable features were then selected independently for each dataset using *FindVariableFeatures* in Seurat, retaining the top 3,000 genes per dataset. The intersection of these features was used for subsequent integration. L5-level clusters were annotated using subclass-level labels transferred from the scRNA-seq data. Clusters containing fewer than ten cells in each region were removed from further analysis. A small fraction (less than 0.01%) of Paired-Tag neuronal cells lacked corresponding co-embedded cells in the reference scRNA-seq dataset, likely reflecting differences in sample collection strategies between the two studies. These cells were instead annotated using transfer label results from a region-agnostic integration, following the same procedure applied to non-neuronal cells.

Additional manual quality control was performed for neuronal populations. For each brain region, major cell classes were projected onto the L1-level UMAP embedding of the Paired-Tag data. Cells deviating from the dominant cluster structure for their assigned class were flagged and excluded from downstream analyses (about 4.26% of cells). After transfer label-based annotations, we had 2,571,700 cells and 4247 clusters.

### Joint co-embedding of multiple single-cell modalities

To evaluate the integration of multiple single-cell modalities, we visualized their joint embeddings following Seurat-based integration. Because of the large scale of the datasets analyzed in this study, we applied a customized Python implementation of the Seurat integration workflow: (https://github.com/scverse/SnapATAC2/tree/main/snapatac2-contrib/snapatac2_contrib/integration)^169^.

We first confined all datasets to cells derived from the nine major brain regions described in the transfer-label analysis. After filtering, the datasets included 656,346 cells from scRNA-seq^6^, 63,792 cells from snmC-seq3^34^, 472,528 from snATAC-seq^35^, and 2,571,700 cells from Paired-Tag.

Next, gene expression levels were simulated for the snATAC-seq and snmC-seq3 datasets. For snATAC-seq, gene activity scores were computed from the number of fragments overlapping each gene body and its 2kb upstream region from the TSS, using the function *snapatac2.pp.make_gene_matrix* in SnapATAC2. For snmC-seq3 data, distinct strategies were used depending on cell type. In neuronal cells except granule neurons (DG Glut), gene expression is inversely correlated with the mCH fraction within the gene body. In non-neuronal cells (NN), immature neurons (IMN) and granule neurons, gene expression is inversely correlated with the mCG fraction in the gene body. Gene bodies were defined as regions spanning 2kb upstream of the TSS to 2kb downstream of the TES. After grouping cells according to these categories, the natural logarithm of the corresponding mCH or mCG fractions was z-score transformed, and the negative values were used as gene activity scores, following previously described methods^34^.

For data integration, the adult mouse brain scRNA-seq dataset was used as the reference, while snATAC-seq, snmC-seq3, and Paired-Tag datasets were treated as queries. Integration anchors were identified using canonical correlation analysis (CCA) with the function *integration.SeuratIntegration.find_anchor*, corresponding to *FindIntegrationAnchors* in Seurat. Only highly variable features shared between the reference and query datasets were used. To mitigate biases introduced by unequal cell numbers across subclasses, we performed downsampling at the cell subclass level for all datasets before integration.

Finally, the identified anchors were used to correct principal components for each query dataset using the function *integration.SeuratIntegration.integrate*, corresponding to *IntegrateData* in Seurat v4. Principal component analysis was performed independently for each modality using more than 1,500 highly variable genes shared across all datasets. The corrected principal components from all four modalities were then merged, and Harmony was applied to further reduce batch effects^173^. UMAP was performed on the Harmony-corrected embeddings to visualize cells from all modalities in a shared two-dimensional space.

### Identification of reproducible peaks for each histone modality

Peak calling was performed following the ENCODE pipeline separately for each sex, histone modification (four in total), and cell subclass. Sex-specific peaks were first identified and then merged to generate subclass-specific peak sets. These subclass-specific peaks were subsequently combined across cell subclasses to derive histone modification–specific peak sets. Detailed procedures are described below.

#### Sex-specific peak calling

For each combination of sex, histone modification, and cell subclass, we first generated a pooled BAM file containing reads from all barcodes within that group. This pooled BAM was then split into four BAM files: two biological replicates and two shuffled pseudo-replicates generated by randomly and evenly splitting barcodes. Replicates with insufficient read depth were excluded from peak calling based on scatter plots of peak number versus read count across BAM files for different histone modifications. Minimum read thresholds were set to 10,000 reads for H3K27ac and 10^4.5^ (31,622) reads for the remaining histone modifications. Peak calling was performed separately for each retained bam files using MACS3^174^ with the following parameters: *--shift -100 --extsize 200 --nomodel --nolambda --keep-dup--all -g mm -q 0.05*. We compared peak calling results obtained with and without the *--nolambda* parameter. Visual inspection showed that inclusion of *--nolambda* retained the majority of biologically relevant peaks without compromising specificity, whereas exclusion of this parameter resulted in fewer detected peaks and loss of signal, likely due to the sparsity of DNA fragments in the dataset. Therefore, *--nolambda* was used in all downstream analyses. Given the distinct chromatin characteristics of different histone modifications, narrow peaks were called for H3K27ac by adding the *--call-summits* parameter, whereas broad peaks were called for the remaining histone modifications using *--broad --broad-cutoff 0.1*. Reproducible peaks were defined as those that satisfied both of the following criteria: (1) a MACS3 negative log10 q-value of at least 0.01, and (2) detection in every replicate or absence from no more than one replicate, as defined by overlap with at least one peak in the corresponding replicate peak sets.

#### Peak merging

Peaks from different cell subclasses were merged using an iterative way implemented in ArchR^175^. This approach uses SPM scores, defined as normalized MACS peak scores or negative log10 q-values, to select representative peaks among overlapping regions, resulting in a non-overlapping and high-confidence peak set. For H3K27me3 and H3K9me3, peaks located on chromosome X were excluded. Peaks overlapping ENCODE blacklist regions were also removed. This merging strategy was first applied to generate unified peak sets for each cell subclass within each histone modification and was then applied again to generate a final unified peak set for each histone modification across all cell populations.

### Chromatin state annotation using ChromHMM

We applied ChromHMM^75^ to annotate chromatin states across the mouse autosomes for each cell subclass by integrating chromatin accessibility from single-nucleus ATAC-seq with histone modification profiles of H3K27ac, H3K27me3, H3K4me1, and H3K9me3 generated by Paired-Tag. Sex chromosomes were excluded from this analysis due to chromosomal-wide processes, such as X-inactivation, on sex chromosomes. Peaks from each modality were first used to generate binarized genomic tracks using the BinarizeBed utility in ChromHMM. Chromatin state models were then inferred using the LearnModel utility. To determine the optimal number of chromatin states, we evaluated a series of models with different state numbers. A 25-state model was selected as a “full” model, which captured all potential chromatin states with redundancy based on empirical observations. Models with 5 to 24 states were then compared with the full model using two complementary approaches, as described previously^54^. First, Pearson correlations were computed between emission probability vectors of each state in a query model and each state in the full model. For each state in the full model, the maximum correlation with any state in the query model was used to quantify how well that state was represented. The average of these maximum correlations across all full-model states was used as a summary metric, with a value of 1 for the full model. Query models with average correlations greater than 0.85 were considered acceptable, and models with average correlations greater than 0.99 were considered satisfactory. Second, k-means clustering was applied to group states within each query model and the full model using emission probabilities, with k equal to the number of states in the model. Clustering performance was quantified using the ratio of between-cluster sum of squares to total sum of squares. The full model served as a reference for maximal separation, and query models achieving at least 95% of the full model’s separation were considered acceptable. The minimal number of states (18) satisfying both criteria was selected as the optimal ChromHMM model.

To facilitate biological interpretation and downstream analyses, we collapsed the original 18-state ChromHMM model into a simplified 8-state chromatin annotation scheme with descriptive state names. State consolidation was guided by the emission probabilities of epigenomic marks, signal patterns observed in genome-wide coverage heatmaps, and the functional association of each state with gene activity. Specifically, Original states 3, 4, 5, 9, 12, and 13 were classified as active chromatin (Chr-A). These states were characterized by strong H3K27ac enrichment accompanied by co-present or flanking ATAC-seq and H3K4me1 signals, consistent with active regulatory elements. Although state 5 did not exhibit a prominent H3K27ac signal in the emission heatmap, inspection of the coverage heatmap revealed a diffuse yet robust H3K27ac signal across these regions, supporting its inclusion as active chromatin. State 9 displayed detectable H3K27me3 emission; however, this state also showed substantially higher ATAC, H3K27ac, and H3K4me1 signals compared with other states and was strongly associated with loci harboring highly expressed genes. Based on these combined features, state 9 was likewise classified as Chr-A. Original states 6 and 8 were classified as poised chromatin (Chr-Po) due to the co-occurrence of H3K4me1 and H3K27me3 signals, together with either chromatin accessibility or H3K27ac enrichment, a pattern consistent with poised regulatory elements. Original state 2 was classified as primed chromatin (Chr-Pr) based on the exclusive emission of H3K4me1, indicative of enhancer priming in the absence of overt activation. Original state 11 was classified as open chromatin (Chr-O) due to its exclusive ATAC-seq signal, reflecting accessible chromatin without accompanying histone modifications. Original states 7, 10, and 14 were grouped as repressed chromatin (Chr-R). These states were defined by strong H3K27me3 enrichment together with the presence of a single active chromatin-associated mark: H3K4me1 in state 7, ATAC-seq in state 10, and H3K27ac in state 14. This combination suggests a repressive chromatin environment with limited or context-dependent accessibility. Original states 15 and 16 were classified as heterochromatin–Polycomb (Hc-P) due to strong H3K27me3 emission in the absence of additional active chromatin marks. Although state 16 also exhibited H3K9me3 emission, it encompassed very few genomic regions and was therefore grouped with Polycomb-associated heterochromatin. Original state 17 was classified as heterochromatin–H3K9me3 (Hc-H), as it was uniquely defined by exclusive H3K9me3 enrichment. State 0 was designated as not determined (ND) because no epigenomic marks were detected. State 18 was also classified as ND due to the simultaneous detection of all assayed epigenomic marks, including ATAC-seq, H3K27ac, H3K27me3, H3K4me1, and H3K9me3, which we were not able to confidently assign to a single descriptive chromatin state category. Notably, this state covered less than 0.1% of the genome and was preferentially localized at highly expressed gene loci. We reasoned that this state represents a rare and negligible genomic fraction arising from the intrinsic open chromatin bias of Tn5-based single-cell genomic assays, and merged it into the ND chromatin state for accuracy. After consolidating the chromatin states, the corresponding emission matrix was recalculated by mapping original state assignments to the 8-state model and re-estimating emission probabilities using binarized signals from each modality.

### Variations of ChromHMM States across cell subclasses

To quantify conservation of ChromHMM states across cell subclasses, we calculated the proportion of overlapping base pairs between subclasses for each chromatin state. For a given ChromHMM state, cell subclass A was treated as the reference and subclass B as the comparison. For each genomic interval belonging to that state in subclass A, the most overlapping interval with the same state in subclass B was identified, and the number of overlapping base pairs was recorded, with a value of zero assigned if no overlap was detected. The conservation proportion between subclasses A and B was defined as the sum of overlapping base pairs divided by the total length of all intervals for that state in subclass A. State variation for subclass A was then quantified as the variance of conservation proportions across all other subclasses.

### Annotation of *cis-*regulatory elements (CREs) using ChromHMM states

For each cell subclass, *cis-*regulatory elements were annotated based on the ChromHMM state assigned to the peak summit, defined as the center of the CRE. The majority of CREs were annotated as Chr-O or Chr-A, followed by Chr-R and Chr-Po. Only a small fraction of CREs were annotated as Hc-P or Hc-H, and no CREs were assigned to the Chr-Pr state.

### Analysis of differential regions using SnapATAC2

Differential region analysis was performed using *snapatac2.tl.diff_test* on peaks identified for each cell subclass and each DNA modality, including ATAC, H3K27ac, H3K4me1, H3K27me3, and H3K9me3. To identify cell subclass–specific differential regions, the *direction* parameter was set to “*positive*” for ATAC, H3K27ac, and H3K4me1, focusing on regions exhibiting significantly higher signal in the target subclass compared with all other cells; the *direction* parameter was set to “*both*” for H3K27me3 and H3K9me3 to characterize regions exhibiting significantly higher or lower signals in the target subclass/class compared with all other cells. The parameters *min_pct = 0.01* was used during differential testing. Differential regions were then defined using an adjusted p-value threshold of 0.05 and a minimum log fold change of 0.2 for each DNA modality. Differential regions were subsequently integrated with ChromHMM annotations. Using this framework, 470,176 cCREs were classified as distal subclass-specific Chr-As, defined as Chr-A overlapping subclass-specific differentially accessible CREs located more than 1 kb upstream or downstream of transcription start sites in a given cell subclass. Gene ontology analysis on differential regions was performed using the Genomic Regions Enrichment of Annotations Tool (GREAT, http://great.stanford.edu/public/html/)^176^.

### Bigwig generation for Paired-Tag sequencing data

BigWig files were generated from merged BAM files using the bamCoverage utility in deepTools^177^ with the parameters: *-e 100 -bs 100 --normalizeUsing RPKM*. The *smoothLength* parameter was set to 300, corresponding to three times the bin size, for H3K27ac, H3K4me1, and H3K27me3 signals; and to 10,000 for H3K9me3 to account for the broader enrichment profiles of this modification; no smoothing was performed for RNA signals.

### Single cell analysis of histone modification and gene expression co-variance

Single-cell covariance between histone modification signal and gene expression was analyzed using a strategy adapted from the peak-gene linking framework implemented in Signac *LinkPeaks* and from the cis-association analysis in SHARE-seq^178,179^. For targeted analyses, candidate regulatory regions located within ±1.25 Mb of the transcription start site of each gene of interest were considered. For each candidate region-gene pair, histone modification signal at the regulatory region and RNA expression of the corresponding gene were quantified across the same cells, and Pearson correlation coefficients were calculated. To estimate background correlations, 100 control regions with matched GC content were sampled from other chromosomes for each candidate region. A background-corrected z-score was then calculated by comparing the observed region-gene correlation to the distribution of correlations obtained from the matched control regions. Candidate links were evaluated based on both the observed correlation and the background-corrected z-score.

To evaluate the relationship between Hotairm1 expression and local chromatin state at single-cell resolution, we analyzed a candidate cCRE upstream of the Hotairm1 locus (chr6:52,144,557-52,173,410). Hotairm1-positive cells were defined as astrocytes (subclasses 318 and 319) with detectable Hotairm1 RNA expression. As a negative-control background, Hotairm1-negative astrocytes were selected and matched to Hotairm1-positive cells within each histone modification dataset. Matching was performed separately for each histone mark to account for the fact that the four histone modification profiles were generated from distinct cell sets. Negative cells were matched based on available technical covariates, including RNA depth and feature number. For each histone modification dataset, reads/fragments overlapping the fixed cCRE region were counted. cCRE signal was normalized by total histone-modification fragment depth per cell as log1p(cCRE count / total fragments × 10,000). Single-cell cCRE detection was defined as cCRE count > 0. Detection fractions were computed separately for Hotairm1-positive and Hotairm1-negative cells, and Wilson 95% confidence intervals were used to estimate uncertainty in detection proportions. Statistical association between Hotairm1 status and cCRE signal detection was further assessed using logistic regression, with cCRE detection as the binary response and Hotairm1 status as the predictor, adjusting for total histone-modification fragment depth and available RNA quality covariates. Odds ratios and 95% confidence intervals were reported for Hotairm1-positive versus Hotairm1-negative cells. To complement the single-cell detection analysis, we performed a matched pseudo-bulk bootstrap analysis. cCRE counts and total fragment depths were aggregated within each bootstrap sample, and pseudo-bulk log2 fold change was calculated as the normalized cCRE signal in Hotairm1-positive cells relative to matched Hotairm1-negative controls. Bootstrap distributions were used to estimate median log2 fold change, 95% bootstrap intervals, and directional probabilities for enrichment or depletion.

### Comparative analysis of Chr-A and Chr-O cCREs

PhastCons conservation scores were computed using the computeMatrix function from deepTools^177^, with the parameters *scale-regions*, *--regionBodyLength 500 –missingDataAsZero --binSize 10*. This analysis was performed separately for Chr-A cCREs, Chr-O cCREs, and randomly shuffled genomic regions within each cell subclass. Conservation profiles were summarized by averaging PhastCons scores at each bin across all cell subclasses, yielding a representative conservation signal for each group of genomic regions. Random genomic regions were generated using the *shuffleBed* function from bedtools^180^ with the parameters *-chrom* and *-noOverlapping*. All cCREs identified from the snATAC-seq data were used as the foreground, and regions overlapping ENCODE mouse representative DNase-hypersensitive sites (rDHS) in the mm10 genome were excluded. Genomic distance analysis was performed only for pairs of cCREs located within the same chromosome. For such pairs, distances were defined as the genomic distance between their peak centers (summits). Distance distributions were calculated separately for Chr-A and Chr-O cCRE pairs within each cell subclass.

To assess the enhancer potential of Chr-A and Chr-O genomic regions, we used results from Allen Institute AAV-based enhancer assays as ground truth. Each tested genomic region was annotated as Chr-A-major or Chr-O-major within the corresponding cell types assayed in the Allen experiments, and these annotations were compared with experimental outcomes. For each Chr-A- or Chr-O-major group, true positives were defined as genomic regions showing positive or putative enhancer activity in the AAV assays, while false positives were regions with negative experimental results. Precision was calculated as the number of true positives divided by the sum of true and false positives. Recall was defined as the number of true positives divided by the total number of regions showing positive enhancer activity in the experiments. Specificity was defined as one minus the ratio of false positives to the product of the total number of genomic regions and the number of tested cell types. Chr-A regions identified by Paired-Tag and brain cCREs defined by snATAC-seq^35^ were also evaluated for enrichment of experimentally validated brain enhancers from the VISTA Enhancer Browser^96^ (https://enhancer.lbl.gov/vista/). Validated brain enhancers were defined as enhancers showing positive LacZ staining in the forebrain, midbrain, hindbrain, or neural tube. Enrichment of the overlap was quantified using odds ratios (OR) calculated from 2×2 contingency tables summarizing the number of regions that overlapped or did not overlap validated enhancers. Statistical significance was assessed using Fisher’s exact test.

### Super-enhancer analysis

Super-enhancers were identified for each brain cell subclass using ROSE with default parameters^118^, based on ranked H3K27ac Paired-Tag signal. To evaluate putative enhancer RNA (eRNA) signal associated with super-enhancers, we first restricted super-enhancers, Chr-A regions, and Chr-O regions to intergenic intervals by removing GENCODE-annotated genic regions, including ±5 kb flanking regions, using bedtools subtract. Putative eRNA signal was then quantified from the corresponding cell type as bidirectional RNA abundance within these intergenic intervals. Specifically, bidirectional RNA abundance was calculated as log10(mean forward-strand RPKM per base + mean reverse-strand RPKM per base + 1), representing the combined RNA signal from both strands after log transformation.

### Motif enrichment analysis

Motif enrichment analysis was performed using HOMER (v5.1)^181^. Known motif enrichment was assessed using *findMotifsGenome.pl* on the mm10 genome, with input regions analyzed at their original lengths (*-size given*). A customized motif library derived from the JASPAR 2022 database (https://jaspar.elixir.no/) was used for known motif analysis. To reduce redundancy when summarizing motif enrichment across cell types, motifs with similar sequence patterns were merged into motif families using non-redundant transcription factor motif clustering (v2.0-beta) as described^161^.

### Identification of sex-biased Paired-Tag L5 clusters

To examine sex-associated cellular heterogeneity at a finer resolution than subclass-level annotations, we analyzed independently defined Paired-Tag L5 clusters. Clusters with fewer than 50 cells were excluded from sex-bias classification. For each remaining cluster, we counted the number of male and female cells based on annotation and used Fisher’s exact test to determine whether the observed male-to-female cell ratio deviated from the global sex ratio of the full Paired-Tag dataset. P values were adjusted across all tested clusters using the Benjamini-Hochberg method. Sex-biased clusters were defined using stringent criteria: adjusted P < 0.05 and >= 95% of cells originating from one sex. Clusters meeting these criteria were classified as male-biased or female-biased according to the predominant sex.

### Analysis of transposable elements (TEs)

Chromatin coordinates and annotations of mouse transposable elements (TEs) were obtained from HOMER using the mm10 reference genome. Only TEs located on autosomal chromosomes (chromosomes 1–19) were included in downstream analyses. Information on the differential accessibility of TEs and predicted TE-promoter connections was adopted from our previous work^35^. TE-Chr-A elements were defined as TE-overlapping cCREs with Chr-A chromatin state annotation. TE subfamily-level activities were calculated using the average histone modification signals across all TEs belonging to that subfamily.

### Deep learning dataset assembly

In addition to processed Paired-Tag histone modification and RNA bigwig tracks, ATAC bigwig tracks were downloaded from https://catlas.org, and DNA methylation bigwig tracks were downloaded from https://data.nemoarchive.org/biccn/grant/u19_cemba/ecker/epigenome/cellgroup/mCseq3/mouse/processed/other/. Train test splits were downloaded from the borzoi manuscript by extracting fold 3 for the test set and fold 4 for the validation set, with the remaining folds used for training. Given the association between low-coverage data and extreme values in track quantification, we employed more aggressive data clipping than the Borzoi manuscript (see the git repo for details)^155^. Data was processed using the *hound_data* script from Baskerville (https://github.com/calico/baskerville) with one notable modification: rather than employing a fixed quantile at blacklist regions, we excluded them along-side genome gaps. Therefore, no blacklist regions were included in training or evaluation. Furthermore, we predicted values for a larger portion of the input sequence (512kb, rather than 196,608 bp). This was achieved by preserving the train/test annotation of contiguous genomic regions used to train Borzoi. As a result, the sequences making up individual train/test pairs are different, but data leakage is prevented.

### Borzoi model training

Models were trained using a modified version of the Baskerville repository, preserving the architecture of Borzoi^155^, but with modifications to the training procedure. Notably, we shifted from a loss-coupled L2 regularization to decoupled regularization^182^, and employed separate learning rates and decay parameters for convolutional and transformer layers. Two training settings were employed: a full fine-tuning setting where the model was initialized using the mouse model from the zeroth borzoi training replicate, and a from-scratch setting to compare the utility of epigenome tracks for guiding gene expression prediction performance. Given the different demands of the tasks, we employed one set of hyperparameters for fine-tuning, and another set was shared across all randomly initialized training tasks. Models trained from scratch were trained for 100 epochs, while the fine-tuned model was trained for 200 epochs. All models were trained using 8 A100s.

### Deep learning model evaluation

Before evaluation, we further filtered lower-quality tracks from the evaluation based on file size. All within-domain model evaluations were performed on unseen test sequences, using either hound_eval, or borzoi_test_genes. For causal variant prediction the TraitGym complex variant test set was downloaded from Hugging Face, consisting of 1,140 causal variants and nine variants with matched minor allele frequency (MAF) for each trait. To evaluate our model’s performance on brain-relevant traits, we subset to variants and matched negatives in TraitGym for the following traits: ‘Alzheimer_LTFH’, ‘Glaucoma_Combined’, ‘Insomnia’, ‘Migraine_Self’, ‘Miserableness’, ‘Mood_Swings’, ‘Morning_Person’, ‘Smoking_Ever_Never’, ‘Worrier’, ‘Risk_Taking’, ‘Suffer_from_Nerves’, “Irritability”, and “Sensitivity”. Trait identification performance was evaluated zero shot using the “L2 of L2” score as described^156^. Model for all AlphaGenome folds were downloaded from hugginface^162^, and inference was performed with alphagenome-pytorch^183^.

### External datasets

1. ENCODE mouse representative DNA hypersensitive site (rDHS) regions for mm10 are obtained from https://www.encodeproject.org/annotations/ENCSR672RVL/ in the ENCODE database.
2. ENCODE mouse candidate *cis-*regulatory elements (cCREs) for mm10 are obtained from https://screen.wenglab.org/ in the SCREEN database (version 4).
3. The bigwig format of PhastCons scores for multiple alignments of multiple species’ genomes to the mouse genome are downloaded from https://hgdownload.cse.ucsc.edu/goldenpath/mm10/phastCons60way/ maintained by the Genome Institute at UCSC.
4. ENCODE mouse blacklist version 2 for mm10 is downloaded from https://github.com/Boyle-Lab/Blacklist.
5. Mouse mm10 (GRCm 38) genome annotation information (version M25 or GRCm38.p6) was downloaded from https://www.gencodegenes.org/mouse/release_M25.html in GENCODE.

### Statistics

No statistical methods were used to predetermine sample sizes. There was no randomization of the samples, and investigators were not blinded to the specimens being investigated. Low-quality nuclei and potential doublets were excluded before downstream analysis as described before. Clustering of these single-nuclei RNA expression profiles or different histone modification profiles was performed in an unbiased manner. Peak callings on different histone modifications followed the procedure used in ENCODE in an unbiased manner.

## SUPPLEMENTAL INFORMATION

Table S1. Paired-Tag oligo sequences.

Table S2. Paired-Tag cell counts.

Table S3. OPC to Oligo variable state regions.

Table S4. Subclass-specific epigenomic features.

Table S5. Subclass motif enrichment in Chr-A.

Table S6. Sex-specific epigenomic and transcriptomic features.

Table S7. Subclass-specific TE-Chr-A regions.

